# Multiomic profiling reveals pericyte and smooth muscle cell contributions to CADASIL pathology in cell-specific Notch3 mutant mice

**DOI:** 10.1101/2025.09.10.675140

**Authors:** Yazi Huang, Veronica Clementel, Mingzi Zhang, Kate Martinez, Karen Martinez, Gavin Spillard, Carina Torres-Sepulveda, Kassandra Kisler, Marcelo P. Coba, Ruslan Rust

## Abstract

Cerebral ischemic small vessel disease (SVD) is the leading cause of vascular dementia and a major contributor to stroke. Cerebral autosomal dominant arteriopathy with subcortical infarcts and leukoencephalopathy (CADASIL) is the most common monogenic form of familial SVD. CADASIL is caused by dominant missense mutations in Notch3, a receptor expressed in mural cells, including smooth muscle cells (SMCs) and pericytes. However, the cell-type specific contributions driving the CADASIL pathology remain unknown due to lack of animal models. Here, we generated two conditional knock-in mouse models carrying the CADASIL-causing *Notch3*^R170C^ mutation selectively in SMC and brain pericytes. Both *Notch3^R^*^170^*^C^*models showed perivascular accumulation of the NOTCH3 extracellular domain, yet developed distinct neurovascular changes depending on the affected cell type. Pericyte-specific Notch3^R170C^ mice displayed pronounced region-selective microglial activation and vascular changes, whereas SMC-specific Notch3^R170C^ mice showed localized perivascular gliosis with minimal vascular remodeling. Proteomic profiling of isolated brain vessels revealed largely unique cell-specific responses. Pericytes Notch3^R170C^ expression dysregulated metabolic pathways, whereas SMC Notch3^R170C^ expression induced immune signaling related pathways. Integration with single-cell RNA-seq data revealed that many of the proteomic and phosphoproteomic shifts might also include brain endothelial cells, including metabolic changes in the presence of pericyte Notch3^R170C^ and inflammatory signaling in the presence of SMC-Notch3^R170C^. Together, these findings define mural cell-specific mechanisms that contribute to the CADASIL-associated vascular pathology.

## Introduction

Cerebral autosomal dominant arteriopathy with subcortical infarcts and leukoencephalopathy (CADASIL) is the most common monogenic form of cerebral small vessel disease (SVD), affecting approximately 5 per 100,000 individuals, though it is likely underdiagnosed.^1–3^ Clinical manifestations in patients include migraine with aura, recurrent subcortical ischemic episodes, cognitive impairment, mood disturbances, and apathy.^4,5^ Neuropathologically, CADASIL is characterized by extensive white matter lesions, neuronal loss, and prominent cerebrovascular pathology involving degeneration and fibrosis of vascular smooth muscle cells (SMCs), ultimately leading to arterial lumen narrowing.^6–8^

CADASIL is caused by dominant missense mutations in the neurogenic locus notch homolog protein 3 (NOTCH3) gene. NOTCH3 encodes a cell surface receptor predominantly expressed in mural cells of the brain vasculature including smooth muscle cells (SMC) and pericytes^9–11^, which interacts with ligands expressed by neighboring endothelial cells.^12^ Mutations cause abnormal accumulation of the extracellular domain of NOTCH3 (NOTCH3ECD) around mural cells. Notably, animal studies suggest that CADASIL pathology arises from novel toxic functions associated with mutant NOTCH3 protein accumulation rather than from loss of normal NOTCH3 signaling.^13^ Previous studies have shown that transgenic mice expressing *Notch3R170C* mutation recapitulate key aspects of human CADASIL including molecular changes^14,15^, vascular pathology^15,16^, disturbed autoregulation^15,16^, white matter injury^17^, and altered neuronal activity^18^, thus enabling translational mechanistic and therapeutic studies between patients^19^ and animals models^20,21^.

However, the cell-type-specific mechanisms connecting NOTCH3 mutations to vascular degeneration remain unclear, largely due to the lack of cell-type-specific Notch3 animal models. This is important because pericytes and SMCs associate with distinct endothelial cell types (capillaries and arterioles respectively^11,22^) and may therefore differentially contribute to the CADASIL pathology. Yet, without models with selective Notch3^R170C^ expression in mural cells, it remains unknown whether pericytes and SMCs drive CADASIL through shared or distinct pathways.

Here, we generated conditional knock-in mice expressing Notch3^R170C^ selectively in pericytes and smooth muscle cells by inserting a Cre-inducible Notch3^R170C^ cassette into the ROSA26 locus. These mice were crossed with brain pericyte- and SMC-specific CreER mice, allowing precise induction of mutant Notch3 upon tamoxifen administration. In both models selective expression of mutant Notch3 in either pericytes or SMCs was sufficient to recapitulate hallmark CADASIL features, including deposition of aggregated Notch3 extracellular domain in brain vessels and distinct neuropathological alterations including vascular and inflammatory responses. Furthermore, using quantitative mass spectrometry of isolated brain vasculature, we identified 13,617 proteins and 74,218 phosphopeptides, enabling parallel proteomic and phosphoproteomic profiling. These analyses revealed divergent, cell-type-specific pathological signatures, with no changes in Notch3 signaling pathways. Mice with Notch3^R170C^ pericytes exhibited changes in metabolic pathways related to energy homeostasis and cytoskeletal processes associated to cell adhesion, whereas mice with Notch3^R170C^ SMCs activated immune and nuclear responses. This study provides, to our knowledge, the first analysis of pericyte- and SMC-specific mechanisms in CADASIL, revealing distinct pathological changes that contribute to vascular pathology.

## Results

### Validation of mural cell-type specific CADASIL mouse models

First, we generated two mural cell-type–specific CADASIL mouse models by crossing the conditional Notch3^R170C^ knock-in line, which carries a lox-STOP-lox cassette inserted into the ROSA26 locus, with tamoxifen-inducible brain pericyte-(ATP13A5-CreERT2^23^, referred to as Per-Notch3^R170C^) and smooth muscle cell-specific (Myh11-CreERT2, referred to as SMC-Notch3^R170C^) Cre driver lines (**Fig. 1a, Suppl. Fig. 1**). We induced mice with tamoxifen at 5 weeks and performed tissue analysis in 8-month old mice to assess long-term effects of Notch3^R170C^ accumulation. To validate our model and confirm cell-type specific expression and accumulation of Notch3, we analyzed Notch3 mRNA expression around small capillaries (<10 µm diameter) and larger arterioles (>10 µm diameter) using RNAscope in situ hybridization (**Fig. 1b, c**). We observed a prominent (>3-fold) increase in Notch3 mRNA accumulation specifically around capillaries in Per-Notch3^R170C^ mice, whereas in SMC-Notch3^R170C^ mice the signal was elevated around arterioles (**Fig. 1b, c**). Consistent with these findings, immunostaining against the NOTCH3 extracellular domain (Notch3^ECD^) revealed robust and specific perivascular protein deposition (**Fig. 1d, e**) mirroring the mRNA patterns in the respective mural cell populations. Importantly, endothelial and non-vascular brain parenchyma showed no signal in mRNA or protein Notch3 signal (**Fig. 1b, d**), confirming the strict mural-cell specificity of the CADASIL mouse models.

**Figure 1:**
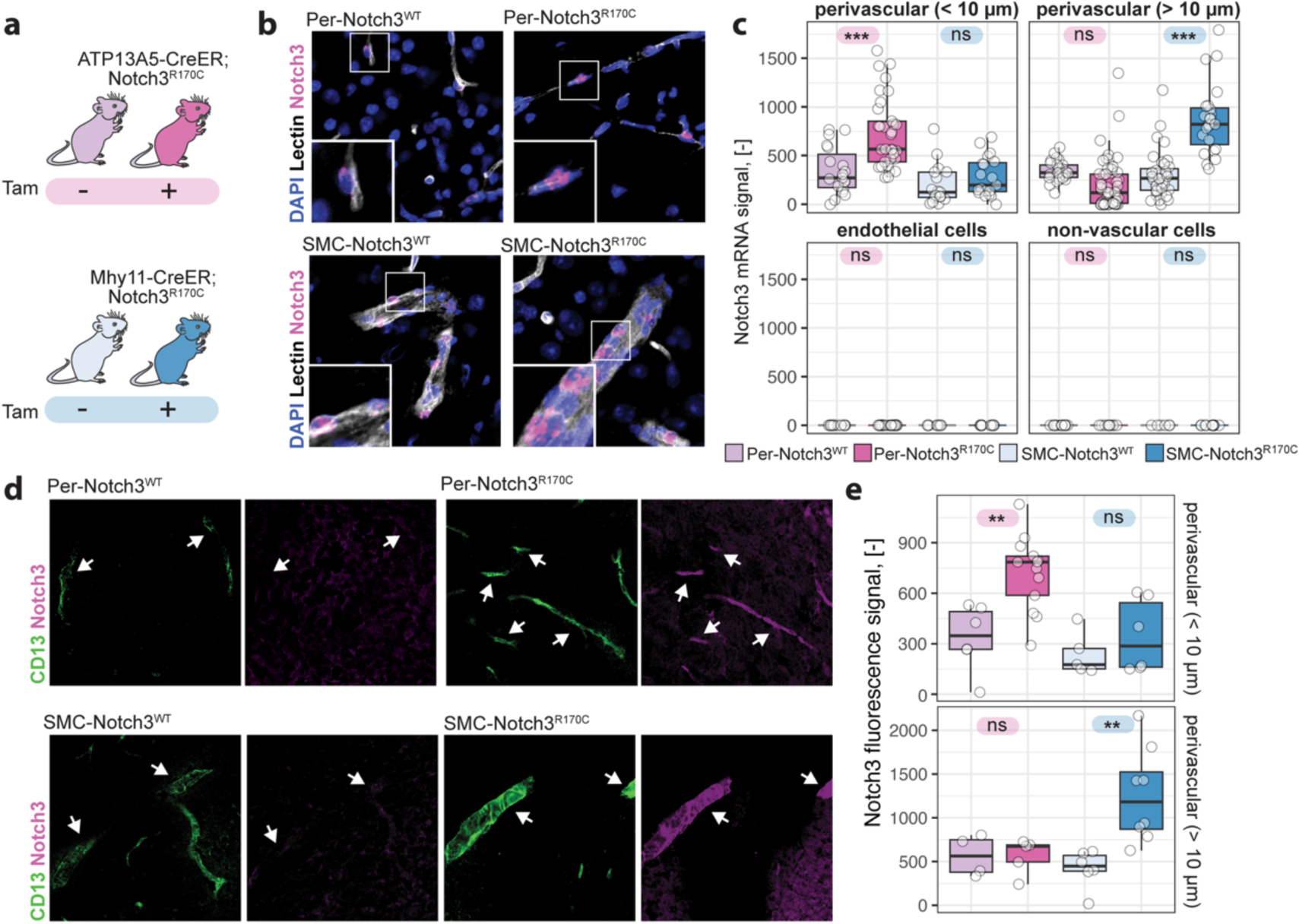
Generation and validation of mural cell-specific CADASIL mouse models. (**a**) Schematic cartoon showing the generation of pericyte (Per)- and smooth muscle cell (SMC)-specific CADASIL mice by crossing conditional Notch3^R170C^ (lox-STOP-lox) mice with ATP13A5-CreER (pericytes) or Myh11-CreER (SMCs). Tamoxifen (Tam) induces cell-specific expression of mutant Notch3. (**b**) RNAscope in situ hybridization of Notch3 mRNA (magenta) with lectin-stained vessels (white) in corpus callosum. Insets highlight cell-type-specific transcript accumulation (capillaries in Per-Notch3^R170C^, arterioles in SMC-Notch3^R170C^). DAPI (blue), nuclei. (**c**) Quantification of Notch3 mRNA confirms mural-cell-specific induction. (**d**) Immunostaining of NOTCH3 extracellular domain (ECD, magenta) and endothelial CD31 (green). Arrows indicate NOTCH3 aggregates restricted to mural cells. (**e**) Quantitative analysis of NOTCH3 protein fluorescence shows selective mural-cell accumulation. In c) Each dot represents an individual vessel from N = 4–8 mice per group. In e) each dot represents a mouse. Box plots show median ± interquartile range; whiskers, min/max; Scale bars, 10 µm. One-way ANOVA with Tukey’s test, ***, P < 0.001, **, P < 0.01, ns = not significant).

Thus, tamoxifen induction of the Notch3^R170C^ allele resulted in cell-type–specific Notch3 upregulation and Notch3^ECD^ accumulation, validating these models for investigating mural cell-specific contributions to CADASIL pathology.

### Changes in vascular and vessel-associated cells in cell-type specific Notch3 mice

Next, we assessed vascular changes in distinct brain regions of Per-specific and SMC-specific Notch3^R170C^ mice compared to their respective Notch3^WT^ control mice. Vessel morphology was comprehensively analyzed to determine vascular area fraction, vacular branching, vessel length, and nearest neighbor distance across the corpus callosum, cortex, hippocampus, and dorsal striatum (**Fig. 2a-f**), as previously established.^24,25^

**Figure 2.**
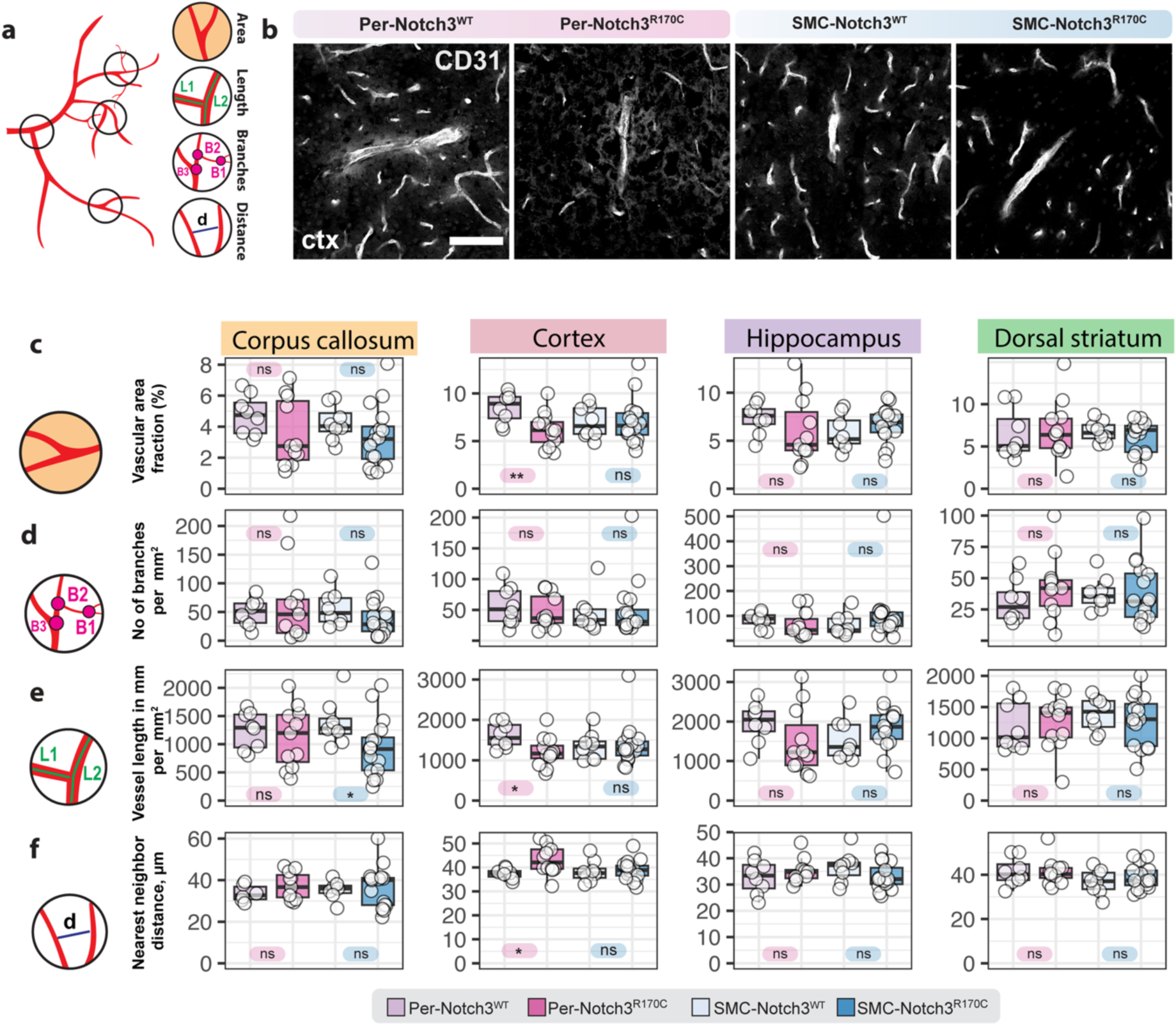
Brain vascular morphology in Notch3 mutant mice. (**a**) Schematic representation of parameters analyzed: vascular area fraction, branching density, total vessel length, and nearest neighbor distance. (**b**) Representative images of CD31-labeled cerebral vessels in the cortex from Per-Notch3 and SMC-Notch3 mice expressing WT or mutant Notch3^R170C^. Scale bar = 50 µm. (**c–f**) Quantification of vascular parameters across brain regions (corpus callosum, cortex, hippocampus, dorsal striatum). Parameters include (**c**) vascular area fraction, (**d**) branching density, (**e**) vessel length density, and (**f**) nearest neighbor distance. Box plots show median ± interquartile range; each dot represents individual mice; whiskers indicate min/max. Statistical significance: *p < 0.05, **p < 0.01, ns = not significant.

In the cortex, Per-Notch3^R170C^ mice showed significantly decreased vascular area fraction (p < 0.01) compared to controls (**Fig. 2b, c**). Additionally, these mice showed a reduction in vessel length per mm^2^ (p < 0.05, **Fig. 2e**) and an increased nearest neighbor distance (p < 0.05), indicating lower vascular density and increased distance between vessels (**Fig. 2f**). No significant changes were observed in the number of branches (**Fig. 2d**). In contrast, SMC-Notch3^R170C^ mice had a mild but statistically significant decrease in vessel length per mm^2^ only within the corpus callosum (p < 0.05), without significant changes in other vascular parameters or brain regions (**Fig. 2e**). No significant differences were observed in hippocampal or dorsal striatal regions across all groups for any of the measured vascular parameters (**Fig. 2e-f**).

Next, we assessed vessel-associated changes, incl. extravascular fibrinogen leakage and smooth muscle actin (SMA) coverage. Quantification of extravascular fibrinogen immunofluorescence showed no significant changes across any brain regions investigated between Notch3^R170C^ and control mice, indicating no increase in vascular permeability (**Fig. 3a, b**). However, analysis of SMA immunostaining revealed significantly reduced vascular coverage in the corpus callosum of Per-Notch3^R170C^ and SMC-Notch3^R170C^ mice (p < 0.05), suggesting region-specific impairment of vessel-associated SMA-expressing cells in both Per-Notch3^R170C^ and SMC-Notch3^R170C^ mice (**Fig. 3c, d**).

**Figure 3.**
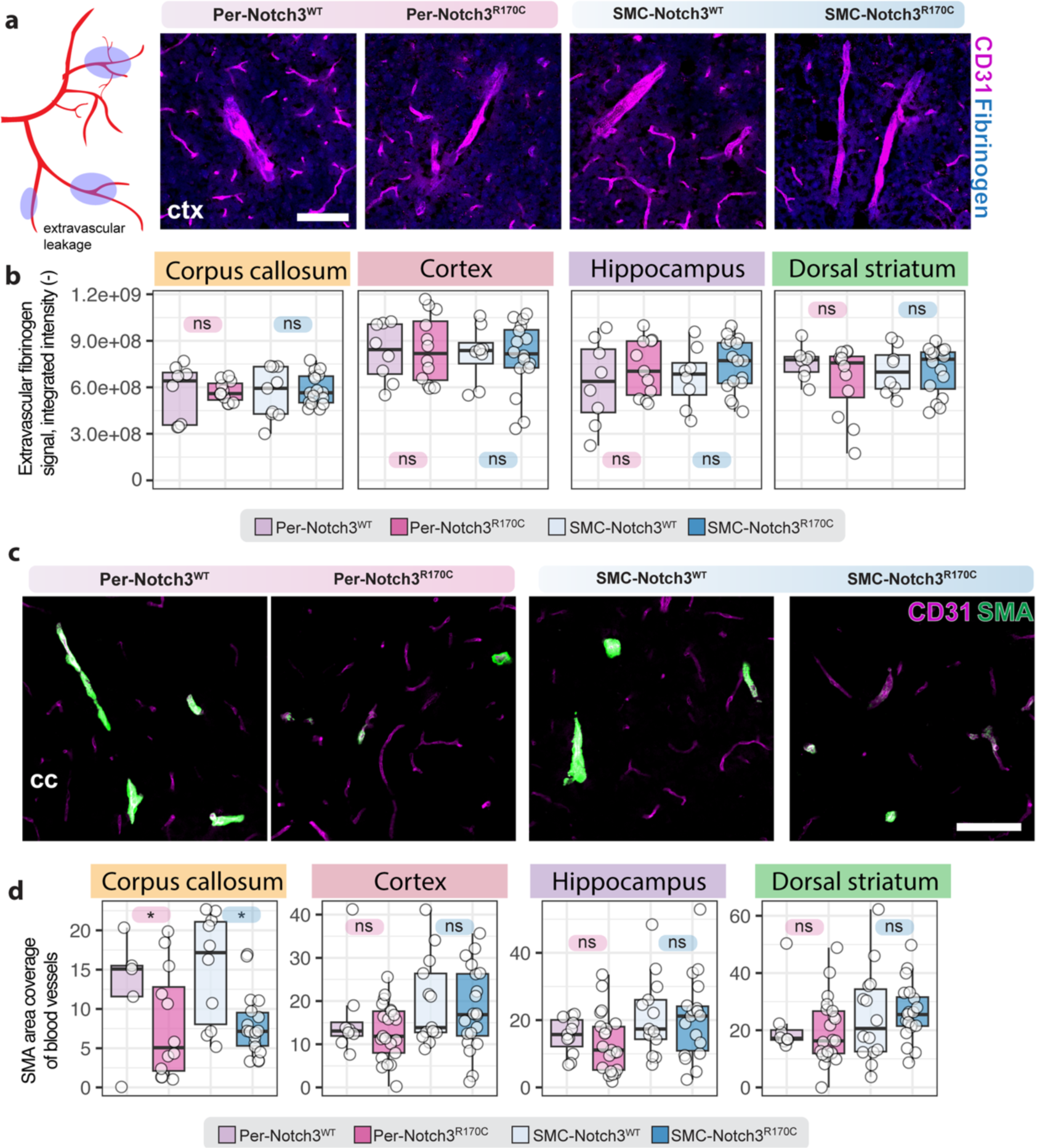
Vessel-associated changes in Notch3 mutant mice. (**a**) Representative images of extravascular fibrinogen (blue) and vascular marker CD31 (magenta) in cortical regions. Scale bar = 50 µm. (**b**) Quantification of extravascular fibrinogen immunostaining across corpus callosum, cortex, hippocampus, and dorsal striatum. (**c**) Representative images showing smooth muscle actin (SMA, green) coverage on CD31-labeled vessels (magenta) in corpus callosum regions. Scale bar = 50 µm. (**d**) Quantification of SMA coverage of blood vessels in Per-Notch3^R170C^ and SMC-Notch3^R170C^ mice compared to controls (*p < 0.05). Box plots show median ± interquartile range; each dot represents individual mice; whiskers indicate min/max. Statistical significance: *p < 0.05, ns = not significant.

Together, these data suggest that while vascular permeability remains intact in both models, Notch3^R170C^ expression in mural cells leads to region-specific reductions in vascular density and SMA coverage, with distinct patterns between pericyte- and SMC-driven Notch3^R170C^ mice.

### Pericyte-specific and SMC-specific Notch3^R170C^ expression induce regional microglial activation

Previous studies have suggested that microglial activation and neuroinflammation contribute to CADASIL-associated pathology.^26,27^ To examine neuroinflammatory changes, we quantified microglial activation markers in multiple brain regions (corpus callosum, cortex, hippocampus, and dorsal striatum; **Fig. 4a**). Per-Notch3^R170C^ mice showed microglial activation characterized by significantly increased Iba1 and/or CD68 expression in all examined brain regions including corpus callosum, cortex, hippocampus and dorsal striatum compared to littermates expressing Notch3^WT^ (**Fig. 4b, c**). By contrast, SMC-Notch3^R170C^ mice exhibited only minor regional changes, with modest CD68 and Iba1 elevation limited to the hippocampus dorsal striatum. Notably, Iba1 and CD68 upregulation did not fully overlap across all brain regions, potentially suggesting that Notch3^R170C^ expression induces distinct microglial activation states that could be associated more with morphological changes (Iba1) and/or phagocytosis (CD68) (**Fig. 4c**).

**Figure 4.**
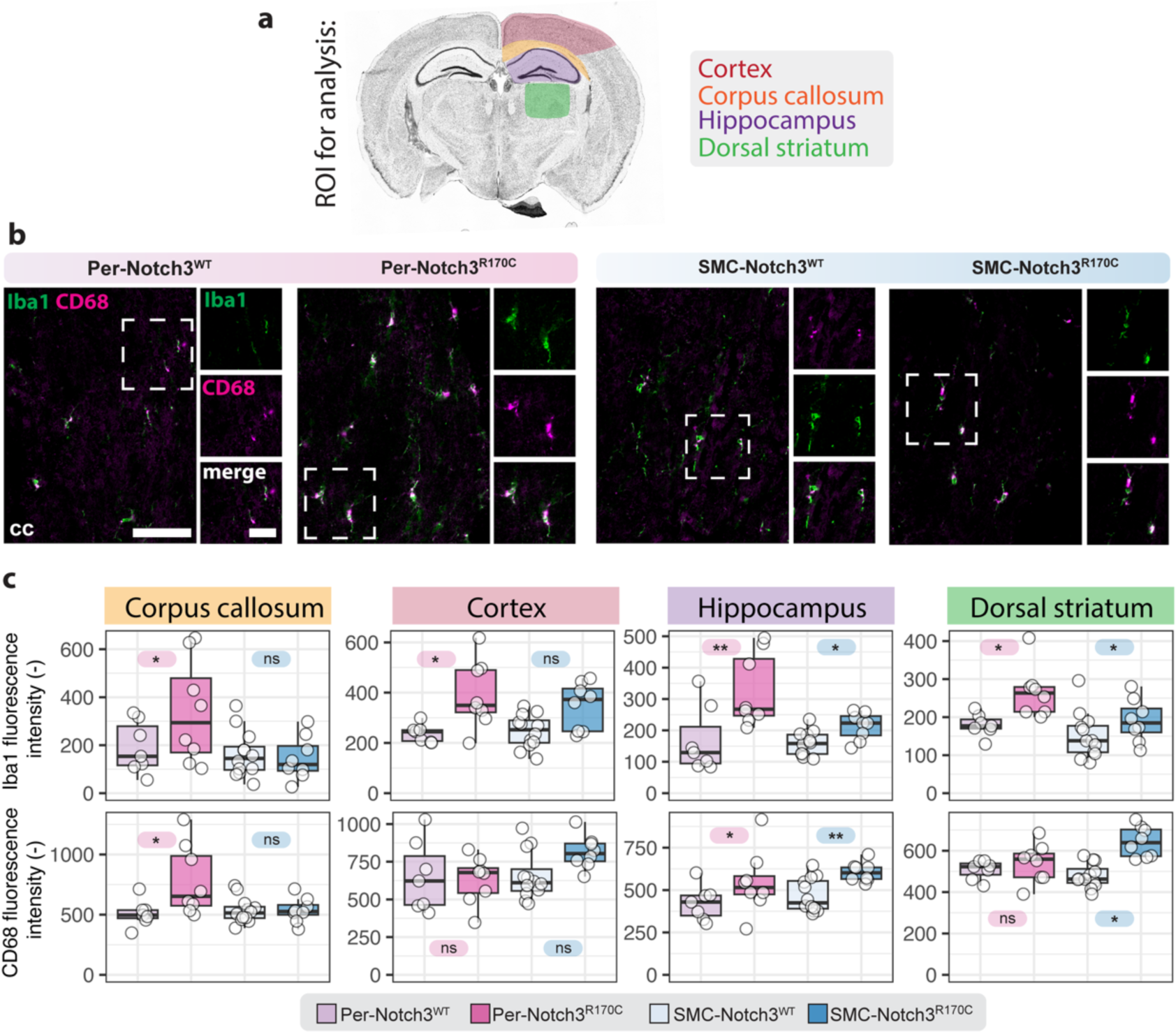
Pericyte-specific Notch3R170C expression triggers regional microglial activation. (**a**) Schematic illustrating analyzed brain regions: cortex, corpus callosum, hippocampus, and dorsal striatum. (**b**) Representative immunostaining for microglial markers Iba1 (green) and CD68 (magenta) showing increased microglial activation in corpus callosum of Per-Notch3^R170C^ mice and SMC-Notch3^R170C^ compared to littermate controls (Per-Notch3^WT^ and SMC-Notch3^WT^). Insets show magnified examples of microglial cells. Scale bars, 50 µm (overview), 20 µm (insets). (**c**) Quantification of Iba1 (upper row) and CD68 (lower row) fluorescence intensity (one-way ANOVA with Tukey’s test; P < 0.05, *P < 0.01, ns = not significant). Box plots show median ± interquartile range; each dot represents individual mice; whiskers indicate min/max.

These findings suggest that pericyte-specific Notch3^R170C^ mutations are sufficient to induce neuroinflammatory phenotypes, closely resembling those of the ubiquitous knock-in CADASIL model.^26^

### Regional and perivascular astrocytic gliosis in mural cell-specific Notch3 mutant mice

Astrocytes and glial scarring have been previously implicated in the progression of CADASIL and other small vessel diseases^15,28^, yet their activation in mural cell-specific Notch3 pathology is unknown. To evaluate astrocytic activation, we quantified GFAP immunoreactivity across multiple brain regions in both pericyte- and smooth muscle-specific Notch3^R170C^ mice (**Fig. 5a,b**). Overall, GFAP^+^ area fraction was comparable between all genotypes, suggesting that cell-type–restricted Notch3 mutations are not sufficient to induce astrocytic gliosis.

**Figure 5.**
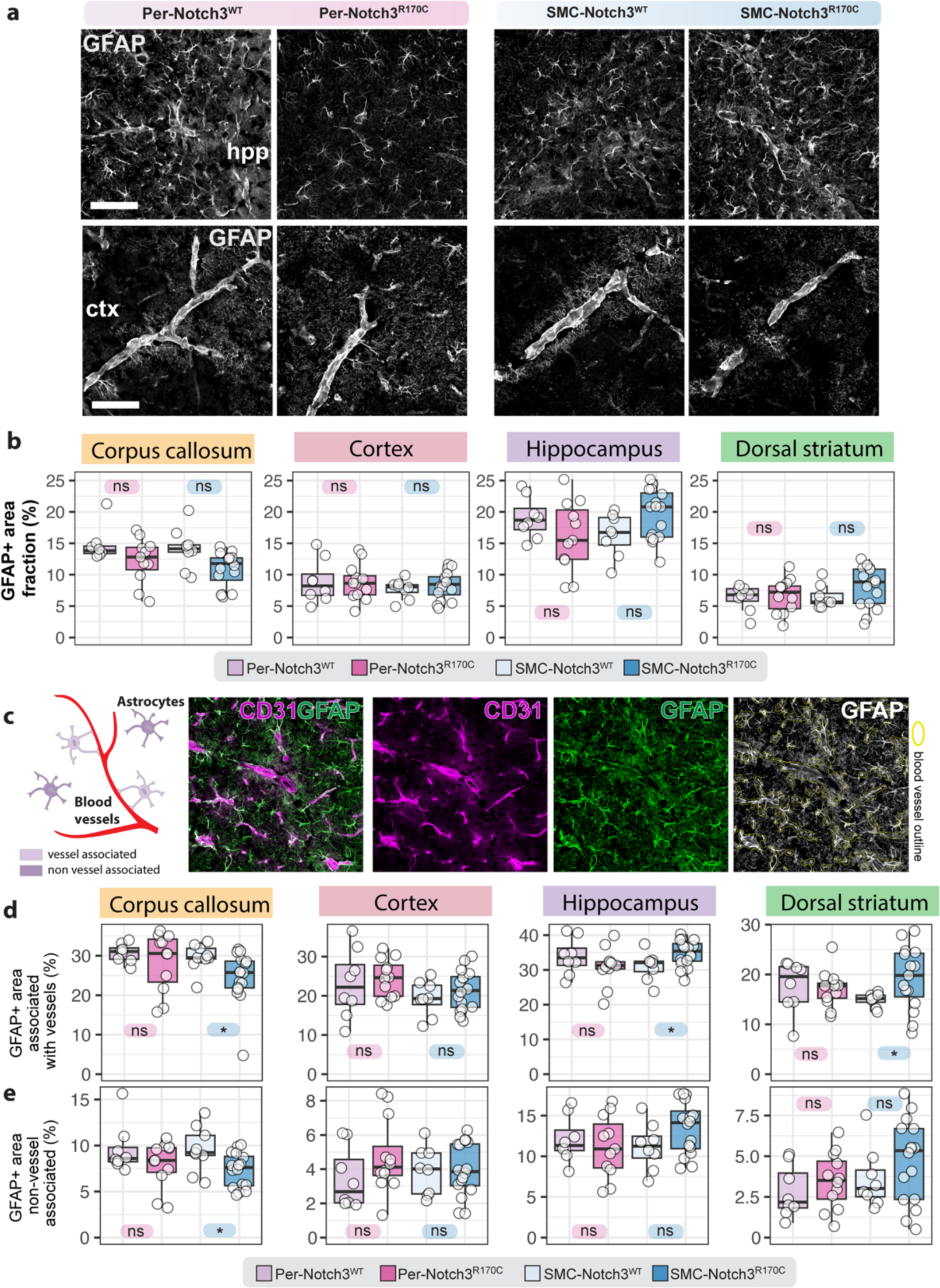
Astrocytic gliosis and vessel-associated GFAP in Notch3^R170C^ mutant mice. (a) Representative images of GFAP immunoreactivity in the hippocampus (hpp) and cortex (ctx) across genotypes. Scale bars, 50 µm. (b) Quantification of total GFAP+ area fraction in corpus callosum, cortex, hippocampus, and dorsal striatum. (c) Schematic and representative images illustrating the classification of vessel-associated and non-vessel-associated GFAP (green) signal using CD31 (magenta) co-labeling. Scale bar = 50 µm. (d) Quantification of vessel-associated GFAP signal. (e) Quantification of non-vessel-associated GFAP. Box plots show median ± interquartile range; each dot represents individual mice; whiskers indicate min/max. Statistical significance: *p < 0.05, ns = not significant.

To further understand whether astrocytic activation occurred in proximity to the vasculature, we developed a semi-automated pipeline to distinguish vessel-associated from non-vessel-associated GFAP signal based on the proximity to the endothelial marker CD31 (**Fig. 5c**). This analysis revealed that, SMC-Notch3^R170C^ mice showed a moderate reduction in GFAP signal in the corpus callosum in vessel associated and non-associated astrocytes (both p < 0.05), and increased GFAP coverage in the hippocampus and dorsal striatum (**Fig. 5c-e**), whereas Per-Notch3^R170C^ mice showed no statistically significant changes across all brain regions.

These findings suggest that while overall astrocyte reactivity remains largely unchanged, SMC-specific Notch3 mutations show a localized perivascular gliosis in subcortical regions, aligning with inflammation patterns (**Fig. 4b, c**) observed in SMC-specific Notch3^R170C^ mice.

### Myelin loss in the corpus callosum of Notch3 ^R170C^ mice

Myelin integrity is often compromised in CADASIL and other small vessel diseases.^15^ To assess potential white matter alterations, we examined myelin basic protein (MBP) immunoreactivity across selected brain regions (**Fig. 6a**).

**Figure 6.**
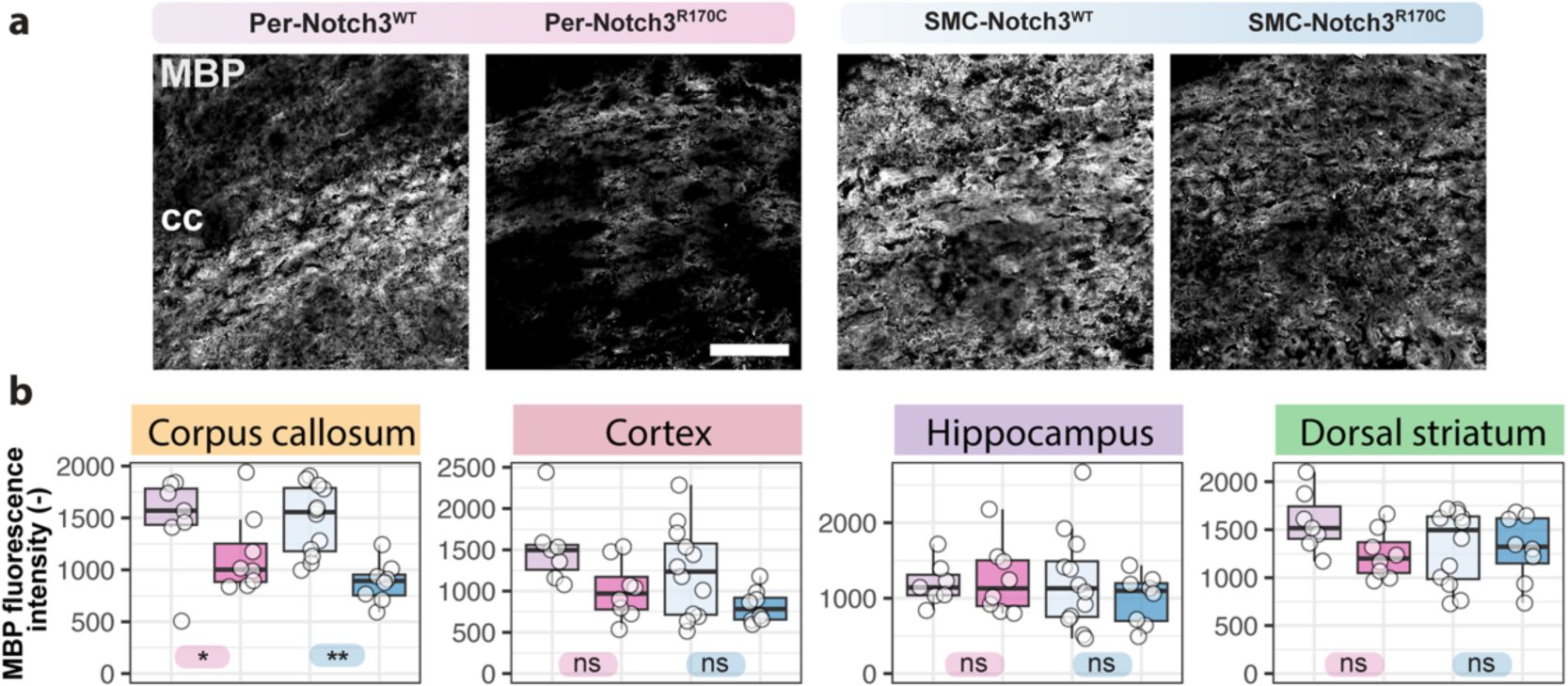
Reduction of myelin basic protein in the corpus callosum of Notch3 mutant mice. (**a**) Representative images of MBP immunostaining in the corpus callosum (cc) of pericyte- and smooth muscle cell-specific Notch3^R170C^ mice and controls. Scale bar = 50 µm. (**b**) Quantification of MBP fluorescence intensity across corpus callosum, cortex, hippocampus, and dorsal striatum. Box plots show median ± interquartile range; each dot represents individual mice; whiskers indicate min/max. Statistical significance: ** p < 0.01, *p < 0.05, ns = not significant.

Quantification revealed a significant reduction in MBP intensity within the corpus callosum of both Per-Notch3^R170C^ and SMC-Notch3^R170C^ mice compared to their respective wild-type littermates (**Fig. 6b**). No significant changes were observed in cortex, hippocampus, or dorsal striatum, suggesting region-specific white matter vulnerability in CADASIL models.

### Proteomic and vascular signatures of pericyte-specific and SMC-specific Notch3^R170C^ expression

To dissect the molecular responses of vascular mural cells to CADASIL-associated Notch3 mutation, we isolated brain vessels (including endothelial cells and their closely associated pericytes and smooth muscle cells) from Per- and SMC-Notch3^R170C^ mice and respective littermate controls (**Fig. 7a**). CD31 labeling confirmed successful enrichment of brain vasculature (**Fig. 7b**), and Western blot analysis verified the accumulation of NOTCH3^ECD^ protein specifically in vessel isolations from both Per- and SMC-Notch3^R170C^ mice (**Fig. 7c**).

**Figure 7.**
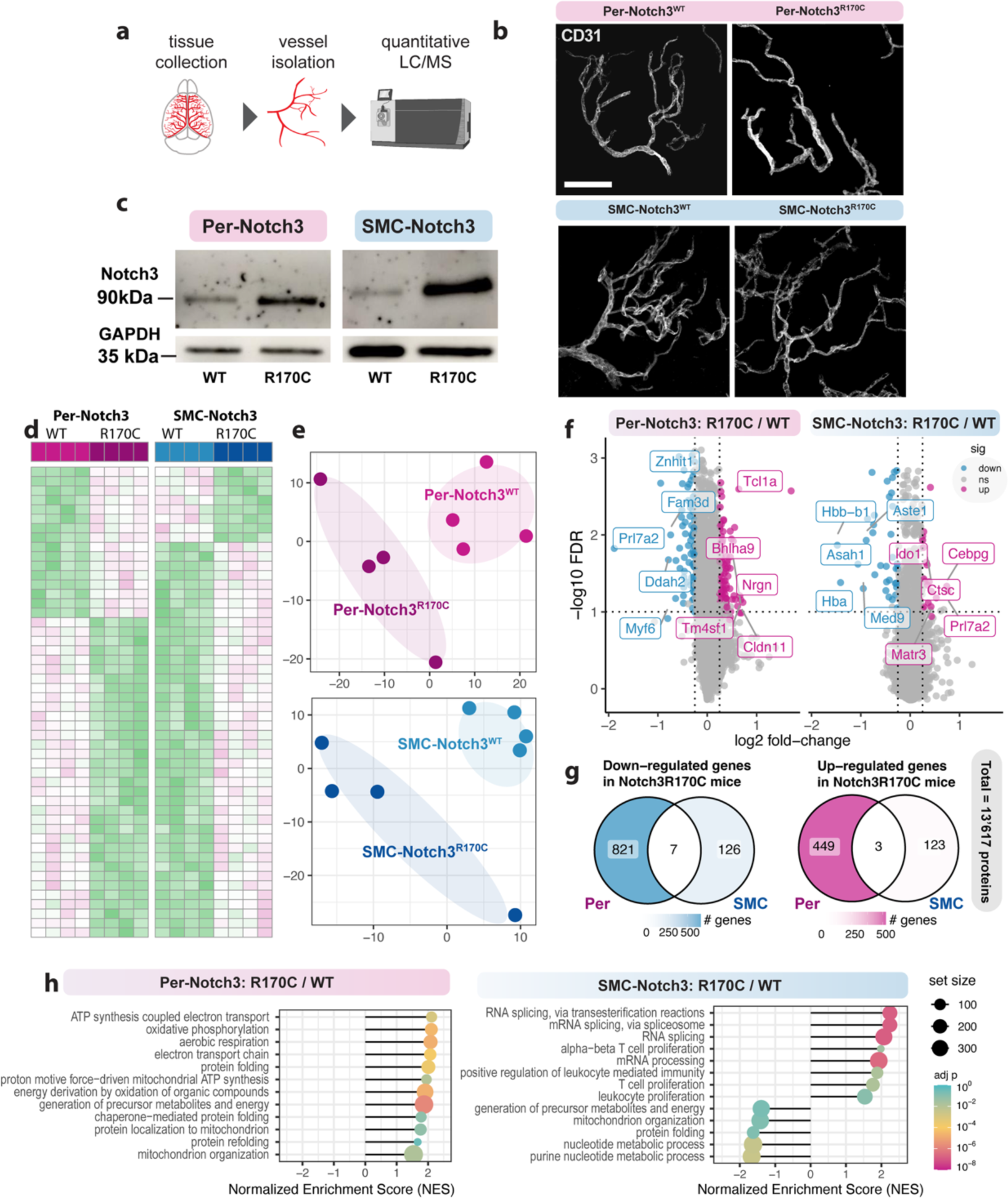
Proteomic profiling of isolated brain vessels reveals cell-type–specific signatures in Notch3 mutant mice. (**a**) Schematic of workflow: brain tissue was collected, vessels were isolated, and subjected to quantitative LC-MS/MS proteomics. (**b**) Representative images of CD31-labeled brain vasculature confirming successful isolation of vascular fragments. (**c**) Western blot of isolated vessels shows accumulation of full-length NOTCH3 (∼90 kDa) in Per-Notch3^R170C^ and SMC-Notch3^R170C^ mice; GAPDH used as loading control. (**d**) Heatmap of differentially expressed proteins reveals distinct profiles between genotypes (Differentially expressed proteins were identified separately in each comparison; the proteins represented in each heatmap are not identical across the two models.) (**e**) Principal component analysis (PCA) and partial least squares-discriminant analysis (PLS-DA) show clear separation of mutant and control groups. (**f**) Volcano plots of differentially expressed proteins (Per-Notch3^R170C^ vs WT and SMC-Notch3^R170C^ vs WT) highlighting top up- and down-regulated proteins, including shared hits LMO3 (up) and DGAT2 (down). (**g**) Venn diagram summarizing the overlap of most differentially expressed proteins (FDR < 0.05; FC > 0.5) between Per- and SMC-Notch3^R170C^ mice. (**h**) Gene set enrichment analysis (GSEA) of proteomic data reveals distinct pathway enrichment patterns: Per-Notch3^R170C^ mice show upregulation of mitochondrial respiration and protein folding pathways, while SMC-Notch3^R170C^ mice display enrichment in RNA processing, chromatin remodeling, and immune pathways. Data represent normalized enrichment scores (NES) for selected top-ranking gene sets.

Quantitative mass spectrometry revealed robust separation of mutant and control proteomes in both mural cell-specific models (**Fig. 7d**), with partial least squares discriminant analyses (PLS-DA) confirming distinct clustering between genotypes (**Fig. 7e**). We quantitated 13’617 proteins in total and identified 1280 proteins with abundance changes in Per-Notch3^R170C^ mice (452 up, 832 down) and 259 proteins in SMC-Notch3^R170C^ mice (126 up, 133 down), with minimal overlap between groups (shared proteins: 3 up, 7 down) (**Fig. 7f, g**). The upregulated shared proteins include UGT2A3, involved in metabolic processing; SLTM, a transcriptional regulator; and H1.8, a linker histone associated with chromatin organization. Among the shared downregulated proteins, ECHDC1 and DGAT2 are directly involved in lipid metabolism, NHLRC3 and GPS1 in protein degradation and signaling, KRT76 and IGSF11 in structural organization and adhesion, and ASTE1 in DNA repair.

Pathway enrichment analysis of proteomic data showed distinct pathway activation profiles between mural cell types (**Fig. 7h**). In Per-Notch3^R170C^ mice, differentially expressed proteins were enriched in pathways related to mitochondrial respiration, oxidative phosphorylation, and protein folding, suggesting a shift in metabolic and protein folding processes compared to their littermate controls. In contrast, SMC-Notch3^R170C^ mice showed enrichment of gene sets linked to spliceosome assembly, chromatin remodeling, and immune signaling, and downregulation of vesicle trafficking pathways compared to their littermate controls.

To independently validate and extend these findings, we applied GeneAgent, a recently developed knowledge-guided AI tool^29^. GeneAgent integrates curated pathway ontologies with literature-derived protein-function relationships to generate functional modules. In Per-Notch3^R170C^, we identified modules related to lipid metabolism, membrane organization, and vascular development, driven by proteins incl. DEGS1, ATP6V0C, and UGT2A3, as well as transcriptional regulators like SOX18 and ACVR1, broadly aligning with our enrichment analysis, suggesting that Per-Notch3^R170C^ dysfunction leads to metabolic and structural remodeling (**Suppl. Data 1**). For SMC-Notch3^R170C^ mice, GeneAgent AI-based analysis identified a functional network centered on leukocyte-mediated cytotoxicity and innate immune regulation, involving transcriptional regulators (SPI1, CEBPG), lysosomal proteases (CTSC), and modulators of autophagy and Toll-like receptor signaling (RUBCN, IRAK1BP1). This immune-skewed profile aligns with the pathway enrichment results and suggests that Notch3^R170C^ expression in SMCs induces proteomic changes that might promote a pro-inflammatory environment (**Suppl. Data 1**).

Overall, these proteomic findings reveal distinct mural cell-type-specific differences in protein expression and pathway enrichment in response to Notch3^R170C^ expression.

### Phosphoproteome changes in pericyte- and SMC-specific Notch3R170C mice mirrors proteomic signatures of CADASIL pathology

To understand post-translational signaling mechanisms disrupted by the Notch3^R170C^ variant in pericytes and SMCs, we performed mass spectrometry analysis of protein phosphorylation in the same fraction of isolated brain vessels used for total protein quantitation. We were able to quantitate 74,218 phosphopeptides in 13,584 proteins (**Fig. 8a, Suppl Table 1**), In the Per-Notch3^R170C^ mice we were able to quantitate a dysregulation of 1,973 phosphopeptides in 1,733 proteins, with the majority downregulated (1,371 phosphopeptides in 1,236 proteins) and only 602 upregulated in 571 proteins (Supp. Table 1, Fig. 8a). (**Supp. Table 1**, **Fig. 8a**). By contrast, in SMC-Notch3^R170C^ mice, 858 phosphopeptides in 801 proteins were altered, with more sites upregulated (617 phosphopeptides in 580 proteins) than downregulated (241 phosphopeptides in 237 proteins) (**Supp. Table 1**, **Fig. 8a**).

**Figure 8.**
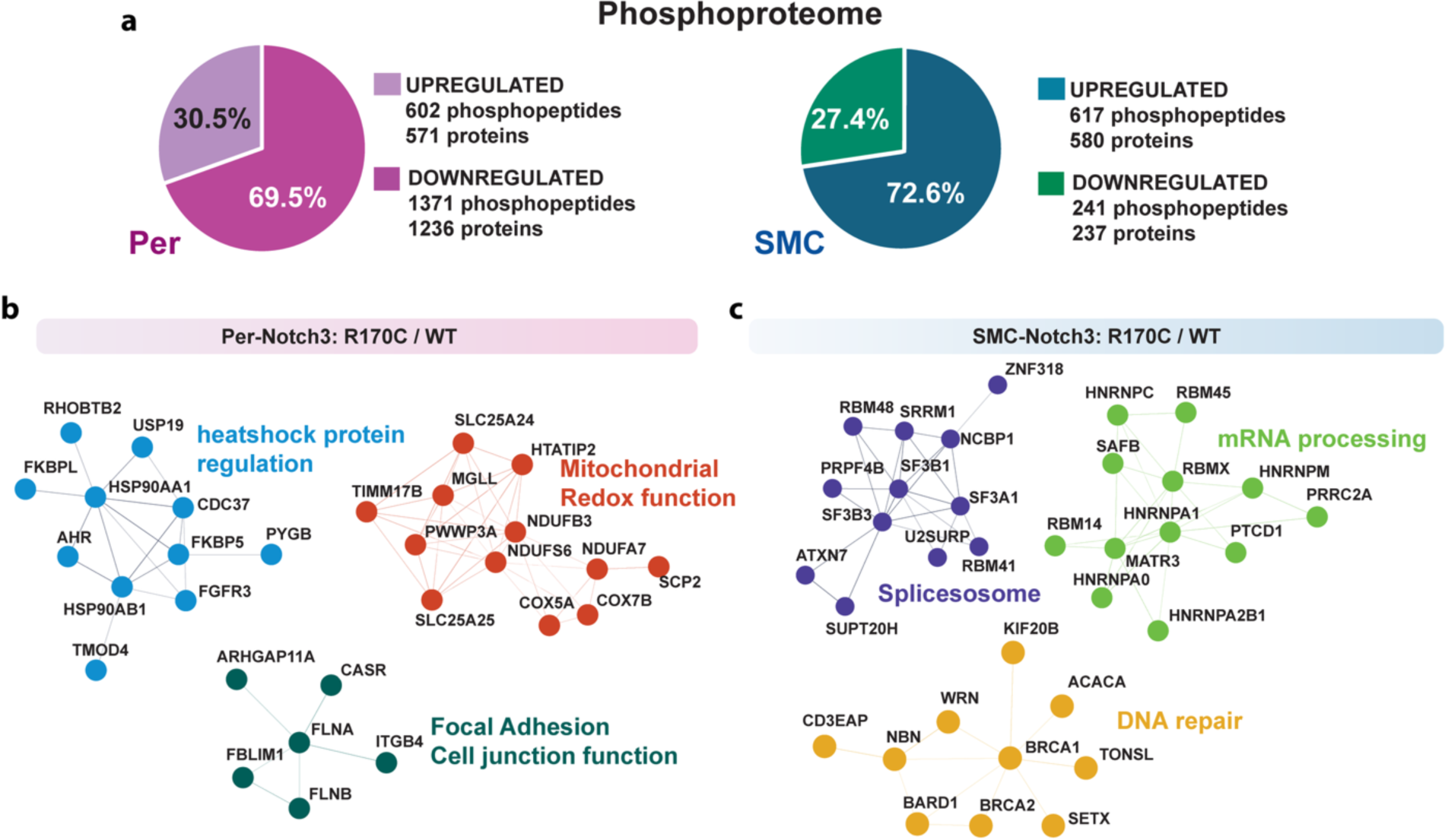
Phosphoproteomic profiling reveals mural cell-type specific signaling alterations in Notch3^R170C^ mutant mice. (**a**) Distribution of significantly up- and downregulated phosphopeptides in isolated brain vessels from pericyte-specific (Per-Notch3^R170C^, left) and smooth muscle cell–specific (SMC-Notch3^R170C^, right) mutant mice compared to respective Notch3^WT^ controls. Percentages indicate the proportion of phosphopeptides exhibiting increased or decreased phosphorylation. (**b**) Network analysis of differentially phosphorylated proteins in Per-Notch3^R170C^ mice highlights functional modules related to heat shock protein regulation (blue), mitochondrial redox function (red), and focal adhesion/cell junction function (green). (**c**) Network analysis of SMC-Notch3^R170C^ phosphoproteome reveals clusters associated with mRNA processing (green), spliceosome assembly (purple), and DNA repair (yellow). Node size reflects the number of connections; edges represent experimentally supported protein– protein interactions.

To gain functional insight, we examined the ontology of proteins exhibiting differential phosphorylation. In Per-Notch3^R170C^ mice, a larger number of cellular pathways were affected than in SMC-Notch3^R170C^ mice, with ontology terms associated with cytoskeletal-microtubule organization and intracellular transport. To determine specific signaling modules that might be affected in each genotype, we first identified protein-protein interactions for each dysregulated proteins using the BioGrid database and then performed a protein interaction network analysis. This analysis has the capacity to identify a set of protein-protein interactions with shared dysregulation in protein phosphorylation. Using this analysis, we were able to identify large, coherent modules associated to mitochondrial metabolism, redox function, protein folding (heat shock proteins), and focal adhesion, alongside smaller disconnected clusters dominated by cytoskeletal proteins (**Fig 8b, Suppl Fig. 2**).

In contrast, SMC-Notch3^R170C^ mice, enriched terms converged on RNA/DNA-related processes, including spliceosome assembly, mRNA processing, and DNA repair. Protein interaction network analysis further supported this shift toward nuclear and post-transcriptional regulation, as many of the dysregulated proteins in these categories formed connected subnetworks (**Fig 8c, Suppl. Fig. 2**).

To explore potential upstream regulators of these phosphoproteomic changes, we next profiled the vascular kinome. Kinome analysis identified protein kinases across all major branches of the vascular kinome, including AGC, CAMK, CMGC, CK1, STE, TK, and TKL groups. Per-Notch3^R170C^ mice displayed a broader spectrum and higher number of significantly altered kinases compared to SMC-Notch3^R170C^ mice (**Suppl. Fig. 3**).

To perform all the protein phosphorylation analysis, we only considered phosphopeptides from proteins with un-altered total protein levels. However, total protein analysis and protein phosphorylation indicated dysregulation of similar molecular pathways in both phenotypes. In Per-Notch3^R170C^ mice, phosphorylation changes recapitulated the metabolic and protein-folding alterations observed in the proteome and further extended to cytoskeletal organization and intracellular transport, whereas in SMC-Notch3^R170C^ mice, phosphoproteomic patterns closely mirrored their proteomic profile, enriched in RNA/DNA-related and nuclear regulatory processes.

### Canonical NOTCH3 proteomic pathways remain largely unaltered in pericyte-specific and SMC-specific Notch3^R170C^ mice

Recent studies suggested that the CADASIL arises from toxic gain-of-function mechanisms associated with mutant NOTCH3 accumulation rather than loss of canonical signaling^13^, we therefore systematically profiled the core components of the NOTCH3 pathway in the isolated brain vessels.

Quantitative proteomics of isolated brain vasculature revealed comparable abundance of NOTCH3 upstream ligands (e.g., Dll4, Jag1), core transcriptional mediators (Rbpj, Maml1), and canonical downstream targets (Hes1, Hey1, Hey2, Nrip2) across all genotypes (**Fig. 9a**). Mapping of the NOTCH3 interactome phosphoproteome showed widespread site coverage in both Notch3^R170C^ models (**Fig. 9b**), with stable phosphorylation detected at five conserved NOTCH3 residues (**Fig. 9c**). All quantified interactors showed consistent total protein levels (**Fig. 9d**). As an example, RBPJ, the key transcriptional mediator of canonical NOTCH signaling, displayed unchanged phosphorylation at three conserved sites (Y65, T37, S176) (**Fig. 9e**)

**Figure 9.**
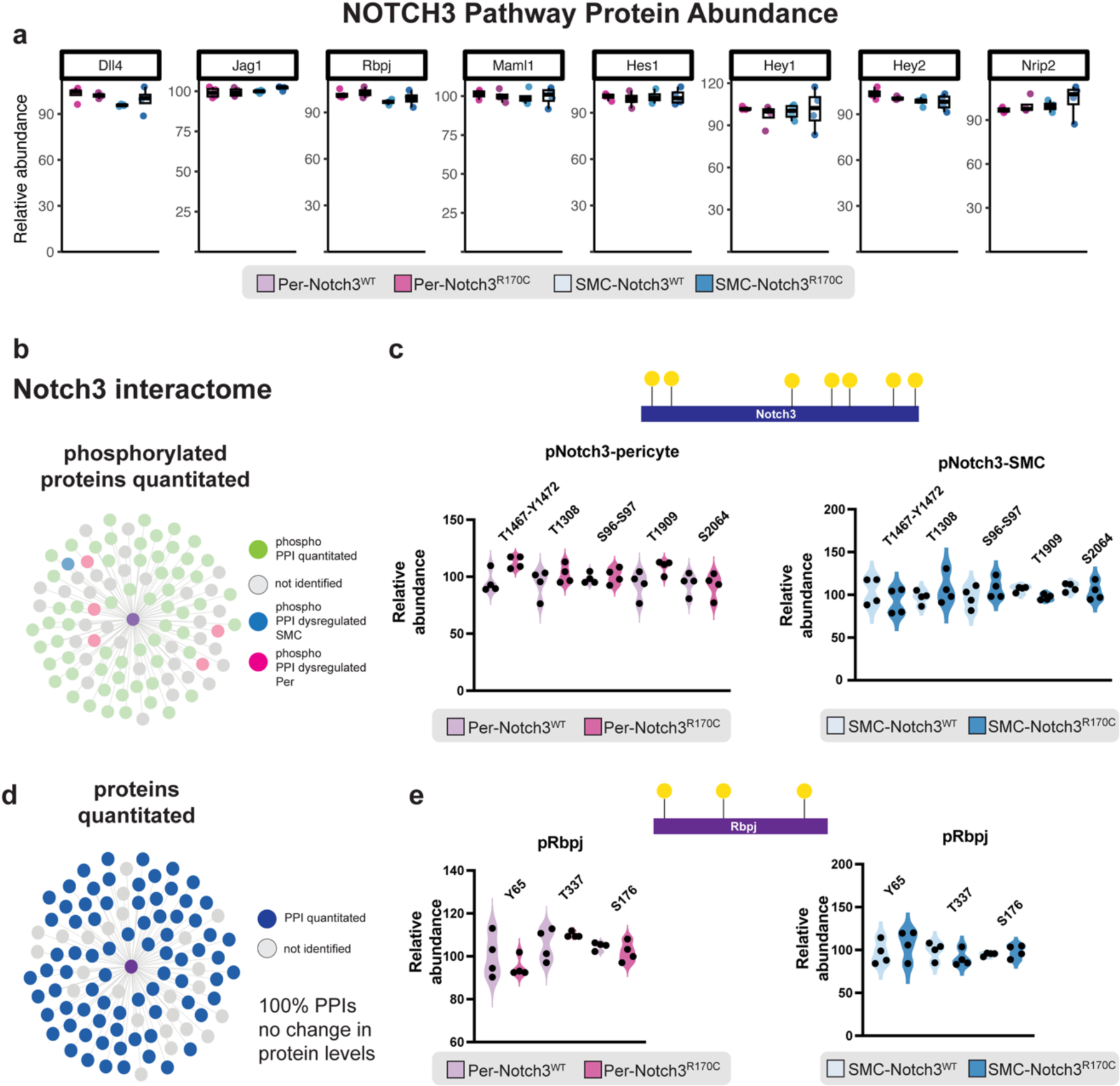
Proteomic and phosphoproteomic analysis of NOTCH3 signaling and interactors. (**a**) Boxplot of relative abundance of canonical NOTCH3 pathway components measured by quantitative proteomics in pericyte- and SMC-specific Notch3^R170C^ mice and respective controls. (**b**) Network diagram of the NOTCH3 interactome displaying phosphoproteins quantified in pericytes (pink), SMCs (blue), both (green), or not identified (gray). (**c**) Dot plots showing relative abundance of phosphorylation at five sites along the NOTCH3 protein in pericytes (left) and SMCs (right), with schematic indicating phosphosite positions. (**d**) Network of proteins within the NOTCH3 interactome with successfully quantified protein-protein interactions (PPIs) shown in blue and non-identified proteins in gray. (**e**) Dot plots showing relative abundance of phosphorylation at three sites on Rbpj (Y65, T37, S176) across mural cell types and genotypes, with schematic indicating phosphosite locations.

These data indicate that canonical NOTCH3 signaling is maintained in brain vessels despite mural cell-specific expression of the CADASIL-associated Notch3^R170C^ variant.

### Single cell RNA-seq guided annotation reveals endothelial-associated proteomic shifts in pericyte and SMC-specific Notch3^R170C^ mice

To explore whether proteomic alterations in brain vessels from Notch3^R170C^ mutant mice might also involve other cell types, we integrated our proteomics data with a curated single-cell RNA-seq dataset of the murine brain vasculature. Using the canonical marker matrix from a previously published brain vasculature atlas^11^, we re-clustered our proteomic dataset and assigned proteins to predicted vascular cell types based on transcriptomic enrichment (z-score > 1, corresponding to expression more than one standard deviation above the mean across cell types, similar to previous studies^30^) (**Fig. 10a,b**). Cell-type predictions were validated by confirming that the average expression of assigned proteins aligned with the appropriate cell clusters in UMAP space (**Fig. 10c**).

**Figure 10.**
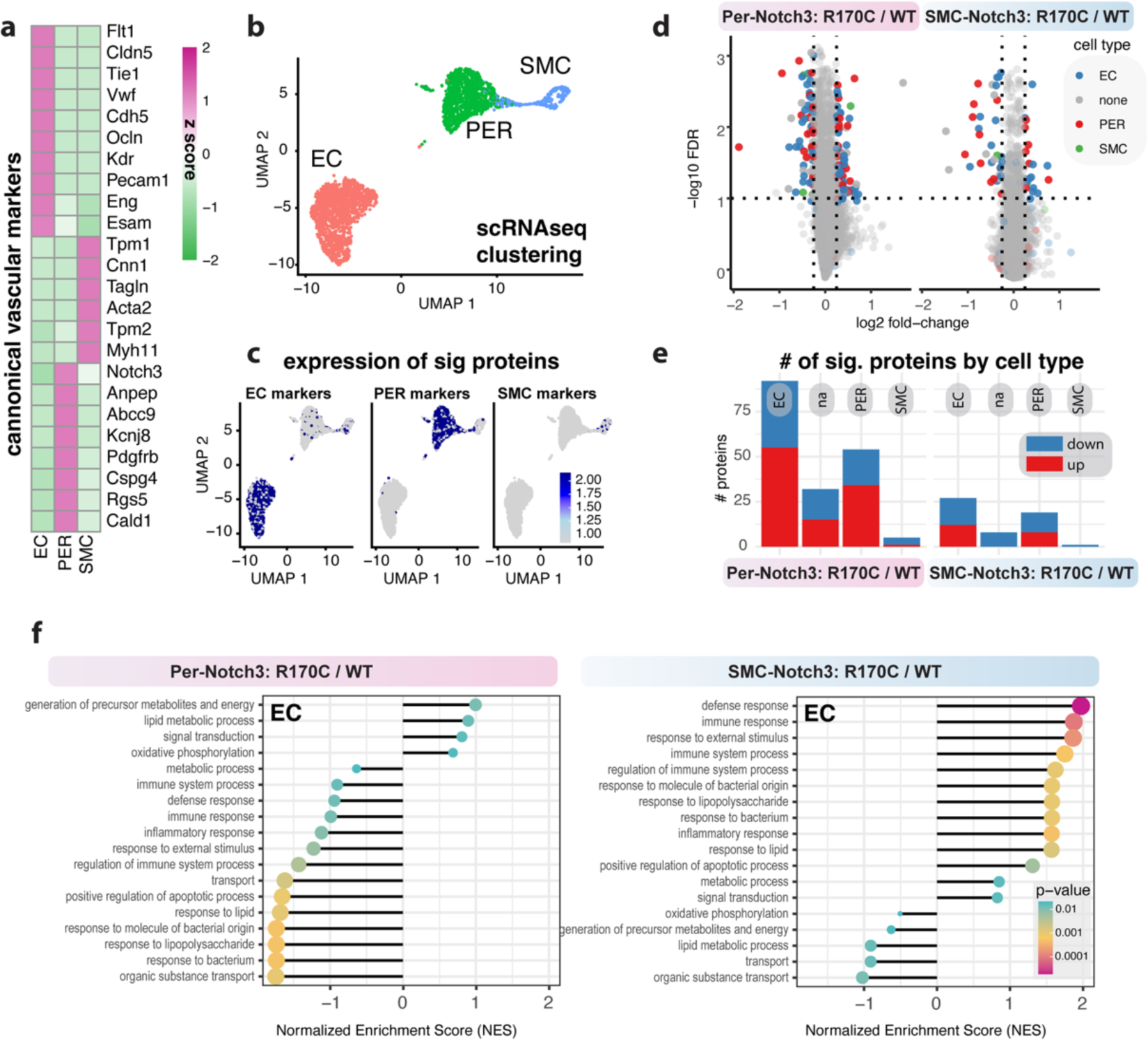
RNA-seq–guided annotation identifies vascular cell-type–specific proteomic profiles in Notch3 mutant mice. (**a**) Heatmap of canonical vascular marker genes from a reference BBB atlas used to assign proteomic profiles to predicted vascular cell types. (**b**) Re-clustering of proteomics data using single-cell RNA-seq–based annotation delineates endothelial cells (EC), pericytes (PER), and smooth muscle cells (SMC) as primary clusters. (**c**) UMAP plots showing the expression distribution of cell-type–assigned proteins. (**d**) Volcano plots of differentially expressed proteins overlaid with predicted cell-type assignments. (**e**) Bar graph summarizing the number of significantly altered proteins by predicted cell type. (**f**) Gene set enrichment analysis (GSEA) of EC-assigned proteins with selected pathways shown. NES = normalized enrichment score; selected top pathways shown.

Next, we mapped the significantly altered proteins from each mutant line onto the annotated clusters. This analysis suggested that many of differentially expressed proteins and phosphopeptides in both Per- and SMC-Notch3^R170C^ mice were enriched in endothelial cells (ECs), with comparatively fewer changes detected in pericytes or SMC-enriched clusters (**Fig. 10d-e**). This suggests that although the Notch3 mutation is restricted to mural cells, many of the proteomic alterations involve proteins with relatively high expression in ECs, suggesting that endothelial responses may also contribute to the observed vascular changes.

To better understand EC-associated changes, we performed pathway analysis specifically on EC-assigned proteins (**Fig. 10f**). In Per-Notch3^R170C^ mice, these proteins showed enrichment in pathways related to mitochondrial ATP synthesis, oxidative phosphorylation, and ER-associated protein processing, reflecting a shift in metabolic activity. In contrast, proteins assigned to ECs from SMC-Notch3^R170C^ mice exhibited enrichments in proteins linked to immune activation, cell adhesion, and cytokine signaling. These findings have also been independently validated using GeneAgent that suggested EC-associated shifts in signal transduction, lipid remodeling and metabolism for Per-Notch3^R170C^ mice and immune regulation pathways in SMC-Notch3^R170C^ (**Suppl. Data 1**).

For proteins assigned to pericytes, Per-Notch3^R170C^ mice showed a reduction in pathways related to metabolic processes, whereas in SMC-Notch3^R170C^ mice, pericyte-assigned changes were more related to DNA- and RNA-associated regulation pathways. In both models, SMC-assigned protein changes were less pronounced and were predominantly associated with broad, non-specific regulatory processes (**Suppl. Fig 4**).

Similarly, in the phosphoproteomic analysis, the networks examined (**Fig. 8b,c**) including heat shock protein regulation, mitochondrial redox, and focal adhesion in Per-Notch3^R170C^, as well as spliceosome, mRNA processing, and DNA repair in SMC-Notch3^R170C^ were predominantly composed of proteins enriched in endothelial cells, indicating that these phosphorylation changes may be at least partly associated with endothelial pathways (**Suppl. Fig 5).**

These findings suggest that some of the proteomic and phosphoproteomic changes might also a contribution from dysregulated pathways in endothelial cells, with distinct patterns emerging in Per- and SMC-Notch3^R170C^ mice.

## Discussion

We generated two mouse models with the Notch3^R170C^ mutation restricted to either pericytes or SMC to define cell type-specific contributions and associated vascular changes to CADASIL pathology. Both models showed Notch3 ECD deposition but distinct cerebrovascular pathology: pericyte Notch3^R170C^ mutation contributed to microglial activation and vascular remodeling, while SMC Notch3^R170C^ mutation induced localized gliosis. Proteomic and transcriptomic analyses suggested that many of these changes do not occur solely in the targeted mural cell but also might have a contribution from dysregulation of endothelial cells function. Within this context we observed that pericyte Notch3 mutations associated with metabolic shifts and SMC Notch3 mutations linked to immune-related signaling.

Previous mouse models have relied on either overexpression of mutant Notch3 or knock-in approaches introducing human or mouse CADASIL mutations.^13,15,26^ While these models have provided critical insights into CADASIL disease pathology, their systemic Notch3 expression patterns did not allow the dissection of mural cell–specific contributions. One difficulty has been the lack of reliable genetic markers to selectively target pericytes. The recent identification of Atp13a5 as a CNS-specific pericyte marker and generation of Atp13a5-Cre mice now enable targeted genetic manipulation of brain pericytes.^23^ Using these mice we restricted the *Notch3^R^*^170^*^C^* allele to either pericytes or SMCs, providing the first evidence that mutations in these mural cell types and the endothelial segments they interact with, lead to distinct patterns of CADASIL pathology. For example, pericyte-specific expression led to more pronounced inflammatory responses and vascular remodeling, whereas SMC-specific expression caused perivascular gliosis with minimal vascular remodelling. Interestingly, some pathological features, such as myelin loss, were observed in both models, suggesting also shared downstream mechanisms independent of the mural cell subtype.

Leakage of the blood–brain barrier (BBB) in CADASIL and sporadic small vessel disease (SVD) has not been consistently observed. While some studies have reported subtle or transient BBB alterations in patients and animal models^16,31,32^, others have found no evidence of overt barrier disruption.^15,33^ In CADASIL mouse models and patients, including those expressing cysteine-altering mutations such as R169C, ultrastructural analyses, tracer permeability assays, and immunostaining for tight junction markers have consistently showed no increased BBB leakage.^15,33^ Our observations are in agreement with these findings, as we did not detect evidence of barrier compromise in our model. However, we cannot exclude the possibility that the absence of detectable BBB leakage reflects a milder or earlier disease stage or is influenced by the cell-type-specific expression of the pathogenic Notch3 allele.

The relationship between Notch3 signaling and extracellular domain (ECD) accumulation remains a central question in CADASIL pathogenesis. While some earlier studies have suggested that altered Notch3 activity may contribute to disease^34,35^, recent work has challenged this idea by showing preserved signaling despite substantial ECD aggregation.^13^ Consistent with these findings, we observed no major changes in canonical Notch3 downstream signaling protein and phosphoprotein expression across our pericyte- and SMC-specific Notch3^R170C^ models, suggesting that downstream signaling remains largely intact. However, proteomic and phosphoproteomic profiling of isolated brain vessels revealed marked differences in Notch3ECD accumulation and associated pathway activation depending on the mural cell type affected. These results support a model in which CADASIL pathology arises not from disrupted Notch3 signaling per se, but from cell type-specific consequences of aberrant Notch3^ECD^ deposition. These findings are further supported by clinical studies showing that the severity of CADASIL mutations correlates with Notch3^ECD^ aggregation, but not with changes in signaling activity.^36,37^ Together with our findings, this reinforces the view that toxic ECD accumulation, rather than impaired canonical signaling, is the key pathological driver in CADASIL.

Our single-cell RNA-seq–guided mapping shows that proteins with dysregulated total levels and phosphorylation signatures, are not only expressed in mural cells, but also in endothelial cells. Importantly, our analysis shows that the RNA expression of these proteins is also relatively higher in endothelial cells, when compared to pericytes and SMCs. Therefore, our findings support the idea that endothelial dysfunction is an early and central feature of CADASIL pathology.^38^ However, the underlying mechanisms of endothelial impairment differ depending on the mural cell type expressing Notch3^R170C^, with pericyte-specific mutations triggering metabolic adaptation, and SMC-specific mutations inducing inflammatory priming. Future studies may further investigate the context-dependent mural-endothelial crosstalk in CADASIL.

In this study mice were analyzed at only one time point (8 months). In future studies, longitudinal assessments, including cognitive and cerebrovascular function, may help to understand the overall disease progression. While vessel isolates were enriched for brain vasculature, we cannot exclude that minor contamination by non-vascular cells could influence proteomic profiles. Our analyses focused exclusively on the brain, and peripheral vascular beds were not examined. Finally, while the *Atp13a5*-Cre and *Myh11*-CreERT2 models enable precise mural cell targeting in mice, the extent to which these patterns mirror human mural cell heterogeneity remains to be determined.

Our findings not only improve our understanding of CADASIL pathology but also could have broader clinical relevance. It has been previously shown that discoveries in CADASIL patients typically translate to sporadic SVD^39–44^ and therefore offer a valuable platform to dissect pathways that are likely relevant beyond the monogenic disorder and into the broader spectrum of age-related cerebrovascular disease. Therefore, our findings reveal cell-type specific pathways that could help identify biomarkers and therapeutic targets relevant not only to CADASIL but also to more common forms of SVD.

## Methods

### Experimental design

We generated a tamoxifen-inducible conditional knock-in mouse model expressing the CADASIL-associated Notch3^ArgR170Cys^ (Notch3^R170C^) mutation in a mural cell–type–specific manner (**Suppl. Fig. 1**). Mutant Notch3 expression was driven from the Rosa26 locus upon Cre recombination and was restricted to either pericytes (ATP13a5-CreER) or smooth muscle cells (Myh11-CreER) using cell-specific CreER driver lines. Mice received tamoxifen (1 mg/day, i.p., for 7 days) between 3 and 5 weeks of age to induce Notch3^R170C^ expression selectively in the target population. Tissues were harvested at 8–10 months post-induction for histological analysis, immunostaining, western blotting, and brain vessels isolation for proteomic profiling.

### Mice

A conditional Notch3^R170C^ (Kozak mutation) knock-in mouse line was generated on a C57BL/6 background (Cyagen Biosciences) using CRISPR/Cas9-mediated genome engineering. A “CAG-loxP-STOP-loxP-Notch3^R170C^-polyA” cassette was inserted into intron 1 of the Rosa26 locus. To achieve mural cell-type–specific expression, these mice were crossed with tamoxifen-inducible CreER driver lines targeting either smooth muscle cells (SMMHC-CreER; JAX #019079) or brain pericytes (ATP13a5-CreER, generated in-house based on our identification of ATP13a5 as a pericyte-specific marker) (**Suppl. Fig. 1**). Double-transgenic offspring (SMMHC-CreER;Notch3^R170C^ or ATP13a5-CreER;Notch3^R170C^) were genotyped and treated with tamoxifen (1 mg/day, i.p.; 10 mg/mL in corn oil) for 7 consecutive days beginning at 3–5 weeks of age to induce mutant Notch3 expression in SMCs or pericytes, respectively. All animal procedures were approved by the Institutional Animal Care and Use Committee of the University of Southern California (Protocol #20678) and conducted in accordance with ARRIVE 2.0 guidelines.^45^

### Brain tissue processing

Mice were anaesthetized with ketamine (100 mg kg⁻¹) and xylazine (10 mg kg⁻¹) followed by transcardially perfused with ice-cold 0.01 M PBS–EDTA (pH 7.4). Brains were dissected, embedded in optimal cutting temperature (OCT) compound, snap-frozen on dry ice, and stored at –80 °C. Coronal sections (20 µm) were cut on a cryostat, mounted onto glass slides (VWR, 48311-703), and stored at –80 °C until use.

### Western blot

Isolated brain vessels pellets were kept on ice and lysed in RIPA buffer (Cell Signaling Technology, 9806) supplemented with protease inhibitors (Roche, 11836153001), followed by probe sonication (20 kHz, 5 s on/10 s off, 3 cycles). Lysates were cleared by centrifugation (12,000 rpm, 15 min, 4 °C), and total protein content in the supernatant was quantified using a BCA assay (Thermo Fisher Scientific, 23228) according to the manufacturer’s instructions. Equal amounts of denatured protein were separated on 10% SDS–PAGE gels and transferred to PVDF membranes (Invitrogen, IB34002). Membranes were blocked in 5% non-fat dry milk (Bio-Rad, 1706404) in TBS-T for 1 h at room temperature, incubated overnight at 4 °C with primary antibodies against NOTCH3 ECD (clone 1E4) (MilliporeSigma, MACB594) and GAPDH (Cell Signaling Technology, 2118), and then with HRP-conjugated secondary antibody (Cell Signaling Technology, 7076, 7074) for 2 h at room temperature. Immunoreactive bands were visualised using SuperSignal West Pico PLUS chemiluminescent substrate (Thermo Fisher Scientific, 34577).

### RNAscope fluorescent in situ hybridization for Notch3 messenger RNA

RNAscope fluorescent in situ hybridization was performed on frozen brain sections using the Notch3 probe set (Advanced Cell Diagnostics, 323110) according to the manufacturer’s instructions. Sections were fixed in 4% paraformaldehyde for 1 h at 4 °C, dehydrated through graded ethanol (50%, 70%, 100%, 100%; 5 min each at room temperature), treated with hydrogen peroxide for 10 min and Protease Plus for 5 min at room temperature. Hybridization with the Notch3-specific probe was carried out for 2 h at 40 °C in a HybEZ oven (ACD, 240200). Signal amplification was achieved sequentially using AMP1 (30 min), AMP2 (30 min), and AMP3 (15 min), each at 40 °C. HRP-C1–based detection was followed by incubation with a fluorescent dye (1:800 dilution, 30 min, 40 °C) and quenched with HRP blocker (15 min, 40 °C). Sections were counterstained with DAPI (1 µg/mL; Invitrogen, D1306) and imaged by confocal microscopy.

### Brain vessel isolation

Brain capillaries were isolated using dextran gradient centrifugation followed by sequential cell-strainer filtrations, as we have previously described (PMID 14583404; PMID 39772387). Briefly, mice were anaesthetized with ketamine (100 mg kg⁻¹) and xylazine (10 mg kg⁻¹) and transcardially perfused with ice-cold PBS. Brains were rapidly dissected into cold Buffer B. Each brain was bisected along the midline using a sterile scalpel (Fine Science Tools, 1003-12), and the brainstem, cerebellum, and hippocampus were excluded. Residual white matter was removed using fine forceps and tweezers (Fine Science Tools, 11051-10 and 11251-35). The remaining brain tissue was minced into uniform flakes and gently homogenised in a glass pestle apparatus (DWK Life Sciences, 885300-002). The homogenate was mixed with 1 mL of 30% dextran (70-kD, Sigma-Aldrich, 31390), vortexed thoroughly, and centrifuged at 6,000 × g for 20 min at 4 °C. The upper layer containing capillary-depleted parenchyma was discarded, and the vessel-rich pellet was passed through 40 µm nylon cell strainers (Corning, 352340). Vessels retained on the surface of the 40 µm strainer were collected and used for downstream staining or proteomic analysis.

### Immunofluorescence staining

Frozen tissue sections were thawed at room temperature and rinsed in PBS to remove OCT, fixed in 4% paraformaldehyde (PFA; ChemCruz, sc-281692) for 10 min at room temperature and blocked in 5% donkey serum in PBS (SouthernBiotech, 0030-01) for 30 min. Sections were incubated overnight at 4 °C with primary antibodies against the following targets: CD31 (BD Pharmingen, 550274, rat, 1:100), CD13 (R&D Systems, AF2335, goat, 1:100), FITC-conjugated α-smooth muscle actin (SMA; Sigma-Aldrich, F3777, mouse, 1:1,000), fibrinogen (Dako, A0080, rabbit, 1:200), GFAP (Millipore, MAB3402, mouse, 1:200), myelin basic protein (MBP; BioLegend, 808402, mouse, 1:100), Iba1 (Biocare Medical, CP290A, rabbit, 1:100), and CD68 (Abcam, ab53444, rat, 1:100). Following primary antibodies, sections were washed in PBS and incubated with the following secondary antibodies (1:500, 2–3 h at room temperature) as appropriate: Alexa Fluor 488–conjugated donkey anti-rat IgG (Thermo Fisher Scientific, A-21208), Alexa Fluor 647– conjugated goat anti-rat IgG (A-21247), Alexa Fluor 568–conjugated donkey anti-mouse IgG (A-10037), Alexa Fluor 568–conjugated donkey anti-goat IgG (A-11057), Alexa Fluor 647– conjugated donkey anti-goat IgG (A-21447), and Alexa Fluor 568–conjugated donkey anti-rabbit IgG (A-10042). Nuclei were counterstained with DAPI (1 µg/mL; Invitrogen, D1306) for 30 s, and slides were mounted with coverslips (VWR, 48404-453). For staining of isolated brain vasculature, vessels were cytospun onto glass slides, fixed in 4% PFA for 10 min, blocked with 5% donkey serum in PBS for 30 min, and incubated with CD31 (rat, 1:100) overnight at 4 °C. The following day, slides were incubated with Alexa Fluor 647–conjugated goat anti-rat IgG (Thermo Fisher Scientific, A-21247, 1:500) for 2–3 h at room temperature, counterstained with DAPI (30 s), and mounted for imaging.

### Confocal microscopy

Brain sections and isolated brain vasculature were imaged on a Nikon A1R HD laser-scanning confocal microscope fitted with 405-, 488-, 561- and 640-nm solid-state lasers. After system start-up, all laser lines were allowed to stabilize for ≈15 min. Slides were mounted on the motorised stage and brought into focus using the objective’s automated collar correction. For each fluorophore, laser power, detector gain and digital offset were adjusted to maximise dynamic range while preventing pixel saturation; the resulting settings were kept constant for all samples within a batch. Four 1024 × 1024-pixel z-stacks (1 µm optical step size) were acquired per section from anatomically matched fields in cortex, corpus callosum, hippocampus and dorsal striatum. Raw image files (.nd2) were archived and subsequently processed in FIJI/ImageJ for quantitative analysis.

### Microscopic analysis

#### Extravascular fibrinogen quantification

To assess perivascular fibrinogen deposition, sections were immunostained with an antibody detecting both fibrinogen and fibrin polymers. Images were acquired from ROIs in the cortex, hippocampus, corpus callosum and dorsal striatum. Using FIJI (ImageJ), fibrinogen and lectin channels were manually thresholded to generate binary masks. To isolate extravascular fibrinogen signal, the binarized lectin-positive vessel mask was subtracted from the fibrinogen mask, and the remaining perivascular signal area was measured using the “Analyze Particles” function. Measurements were averaged across 3–5 ROIs per region per mouse.

#### Intensity analysis of Iba1 and CD68 inflammation

For Iba1 and CD68, fluorescence intensity was quantified directly in defined anatomical ROIs using the “Measure” tool in FIJI. To normalize the signal uniformly across samples, background subtraction was applied.

#### Analysis of glial scarring GFAP

GFAP signal was thresholded to generate a binary mask. GFAP positive area was expressed as a percent of the total ROI area. For vessel-associated GFAP analysis, binary masks were generated from the CD31 channel, dilated by ∼2 µm, and applied to segment GFAP signal into vessel-associated and non-vessel-associated compartments prior to quantification.

#### MBP intensity

For MBP, fluorescence intensity was quantified directly in defined anatomical ROIs using the “Measure” tool in FIJI. To normalize the signal uniformly across samples, background subtraction was applied.

#### Vascular analysis of area fraction, branches, length and nearest neighbor distance

CD31-labeled sections were used to extract vascular parameters including area fraction, total vessel length density, branchpoint count, and nearest neighbor distance (NND). Images were binarized in FIJI using a fixed threshold across conditions, and skeletonized vessel maps were generated using the “Skeletonize” and “Analyze Skeleton” plugins. Vessel area and length density were measured from the original binarized images, whereas branchpoints and NND were extracted from the skeletonized mask. Four fields per region were analyzed and averaged per individual animal, as previously described.^24,25,46,47^

#### SMA coverage analysis

To quantify smooth muscle actin (SMA) coverage of blood vessels, CD31 and SMA channels were thresholded individually, and binary masks were created. To quantify the SMA signal the CD31-mask was dilated by 2 µm to analyze the perivascular area. The resulting SMA-positive vessel area was expressed as a percentage of total CD31+ area per ROI.

### Proteomics

Brain vessels were isolated from capillary-enriched cortical tissue of 4 mice/group (Per-Notch3^R170C^, Per-Notch3^WT^, SMC-Notch3^R170C^, SMC-Notch3^WT^) at 8-10 months of age and frozen in liquid nitrogen.

### Sample preparation

Samples were received in dry ice and dissolved in 500 µL lysis buffer (0.5M triethylammonium bicarbonate, 0.05% sodium dodecyl sulfate). Lysates were subjected to tip sonication (Q700, QSonica, amplitude = 10, 2 sec on/2 sec off pulses, 20 sec total processing time per sample, on ice) and then centrifuged at 15,000 rpm at 4 °C for 10 min. The supernatant was transferred to a fresh tube, and protein concentration of each sample was determined using the Qubit Protein Assay Kit (Thermo, Q33211) and Qubit 4.0 fluorometer per manufacturer’s instructions.

Equal amount of protein (40 µg) per sample was transferred to a fresh tube adjusted to a final volume of 90 µL with lysis buffer. Reducing reagent (4 µL; Sigma, 4381664) was added, and samples were incubated at 60°C for 1hr. Alkylating reagent (2 µL; Sigma, 4381664) was then added, followed by incubation at room temperature for 15 min. Trypsin/LysC (2 µg; Promega, V50703) was added, and samples were incubated overnight at room temperature in the dark.

TMTpro reagents (Thermo, A44522) were equilibrated at room temperature, and 20 µL of anhydrous acetonitrile (Sigma, 900644) were added to each label and the contents were then transferred to each sample. Samples were incubated at room temperature for 1hr, followed by the addition of 8 µL 5% hydroxylamine and a further 15 min incubation at room temperature. Samples were then combined and dried-up using a speedvac (Eppendorf 5301 vacufuge concentrator).

Phosphopeptide enrichment was performed on the combined TMTpro-labeled peptides by sequentially applying the High-Select TiO₂ Phosphopeptide Enrichment Kit (Thermo, A32993) and the High-Select Fe-NTA Phosphopeptide Enrichment Kit (Thermo, A32992), according to the manufacturer’s guideline. The resulting TiO2 and Fe-NTA enriched phosphopeptide fractions were subjected to reverse-phase fractionation (Thermo Scientific Pierce, 84868). The flow-through from the phosphoenrichment, containing native peptides, was subjected to offline alkaline C8 reverse phase fractionation. Twenty-three fractions were collected in a time-dependent manner and dried-up using a speedvac. These were desalted using C18 tips with 100 µL bed per manufacturer’s instructions (Thermo Scientific Pierce, 87784).

### Quantitative proteomic analysis with LC-MS

The desalted peptide fractions were reconstituted in water 0.1% formic acid and analyzed using LC-MS with FAIMS ion mobility pre-separation (nano-easy LC 1200, Thermo Orbitrap Exploris 480).

### Proteomic data processing

For the native peptides, raw data files of the LC-MS analysis were submitted to Proteome Discoverer 2.5 (Thermo) for target decoy search using Sequest against the mus musculus canonical swissprot database (TaxID= 10090, v2021-07-30). The search allowed for up to two missed cleavages, a precursor mass tolerance of 20ppm, a minimum peptide length of six and a maximum of three equal dynamic modifications of oxidation (M), deamidation (N, Q) or phosphorylation (S, T, Y). Methylthio (C) and TMTpro (K, peptide N-terminus) were set as static modifications. Peptide level confidence was set at q<0.05 (<5% FDR). Normalization mode of total peptide amount was applied. Only unique peptides were considered for protein quantitation. All mass spectrometry proteomics data will be deposited to the ProteomeXchange Consortium via the PRIDE partner repository.

### Differential-abundance analysis

Processed files from Proteome Discoverer v2.4 (13,617 proteins, 4 × 4 biological replicates) were log₂-transformed in R v4.4. Differential abundance between mutant and control samples was tested separately for pericyte and SMC datasets with an unpaired moderated t-test (genefilter::rowttests), followed by Benjamini–Hochberg correction. Proteins with |log₂FC| > 0.25 and FDR < 0.10 were taken forward to Gene-Ontology enrichment (clusterProfiler). Protein identities were mapped to endothelial, pericyte or smooth-muscle bins using a single-cell vascular atlas (transcript z ≥ 0.7) to enable cell-type-resolved pathway analysis using Seurat v5, similar to previous analysis pipeline^30,48–50^. Cell-type–specific hit lists were subjected to independent GO-BP ORA and visualized as dot plots.

We used Gene Agent, a self-verification language agent for gene-set analysis^29^, to independently validate the pathway enrichment results. For Per- and SMC-Notch3^R170C^ mice, the analyses can be found as: <u>GeneAgent: SMC-Notch3</u> and <u>GeneAgent: Per-Notch3</u>; The corresponding analyses for annotated endothelial cells are provided separately under: <u>GeneAgent: SMC-Notch3-EC</u> and <u>GeneAgent: Per-Notch3-EC</u> (**Suppl. Info 1**)

### Statistical analysis

Sample sizes were estimated using nQuery Advisor (Statsols) based on a two-sided α = 0.05, 80% power, and effect size estimates derived from prior studies of cerebrovascular dysfunction in mouse models.^30,48,51–53^ For normally distributed data with equal variances, one-way ANOVA followed by Bonferroni post hoc comparisons was used. For two-group comparisons, unpaired two-tailed Student’s t-tests were applied. All statistical analyses were conducted using GraphPad Prism 10.2 and R v4.3. A two-sided p-value < 0.05 was considered statistically significant. Exact tests, sample sizes, and p-values are detailed in the figure legends.

## Data availability

All mass spectrometry proteomics data will be deposited to the ProteomeXchange Consortium via the PRIDE partner repository upon publication of the peer-reviewed article. Single cell RNAseq data were used from GSE98816. All other supporting data are available for reproducibility purposes from the corresponding author upon request.

## Supporting information

All Suppl. Figures

## Acknowledgement

This work is supported by the funding from Swiss 3R Competence Center (OC-2020-002), the Swiss National Science Foundation (PZ00P3_216225) and USC Dean’s Pilot Funding Award to RR.

## Author contributions statement

M.P.C., R.R. contributed to overall project design. Y.H. performed brain vessel isolations. V.C., M.C., and R.R. prepared samples for LC/MS and performed all proteomics analyses. R.R. performed scRNAseq integration. Y.H., V.C., M.Z., Kat. M., Kar.M., G.S., C.T.S., R.R. conducted and analyzed experiments. K.K., M.P.C., R.R. supervised the study. M.P.C., R.R. wrote and edited the manuscript with input from all authors. All authors read and approved the final manuscript.

