## Supplementary material for "Multiomic profiling reveals pericyte and smooth muscle cell contributions to CADASIL pathology in cell-specific Notch3 mutant mice": All Suppl. Figures

### Title

### Authors, TBD

Yazi Huang[1,2], Veronica Clementel[1,2], Mingzi Zhang[1,2], Kate Martinez[1,2], Karen Martinez[1,2], Gavin Spillard[1,2], Carina Torres-Sepulveda[1,2], Kassandra Kisler[1,2], Marcelo P. Coba[1,2,3,*], Ruslan Rust[1,2,*]

* co-senior authors

### Affiliations

[1] Department of Physiology and Neuroscience, University of Southern California, 90033, Los Angeles, USA
[2] Zilkha Neurogenetic Institute, Keck School of Medicine, University of Southern California, 90033, Los Angeles, USA.
[3] Department of Psychiatry and Behavioral Sciences, University of Southern California, 90033, Los Angeles, USA

### Correspondance

Ruslan Rust, Ph.D.
Assistant Professor
The Zilkha Neurogenetic Institute,
Department of Physiology and Neuroscience
Keck School of Medicine of the University of Southern California
1501 San Pablo Street
Los Angeles, CA 90033

ORCID: 0000-0003-3376-3453

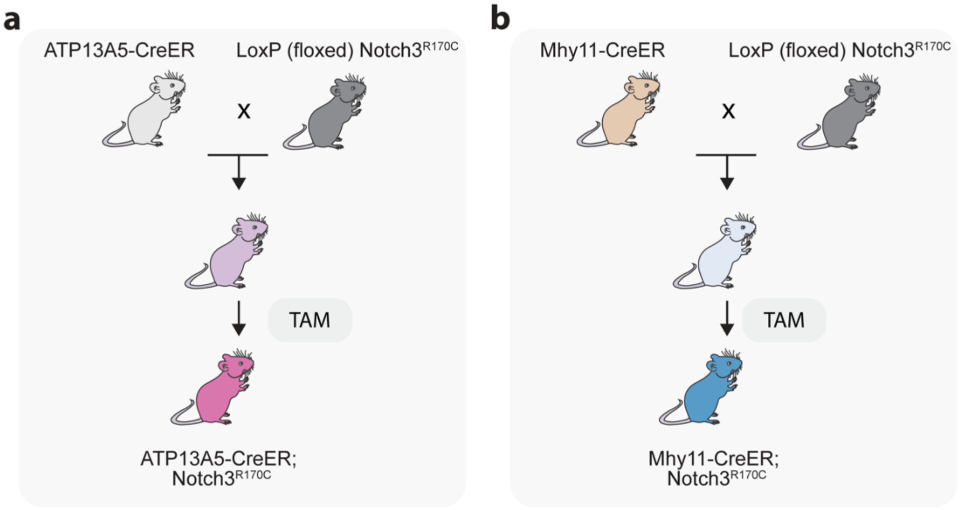


**Supplementary Figure 1. Generation of mural cell-specific mouse models**. ATP13A5-CreER (pericyte-specific) or Myh11-CreER (SMC-specific) mice were crossed with LoxP-Notch3$^{R170C}$ mice. Upon tamoxifen (TAM) administration, Cre-mediated recombination induces expression of mutant Notch3 in mural cell subtypes**.**

a

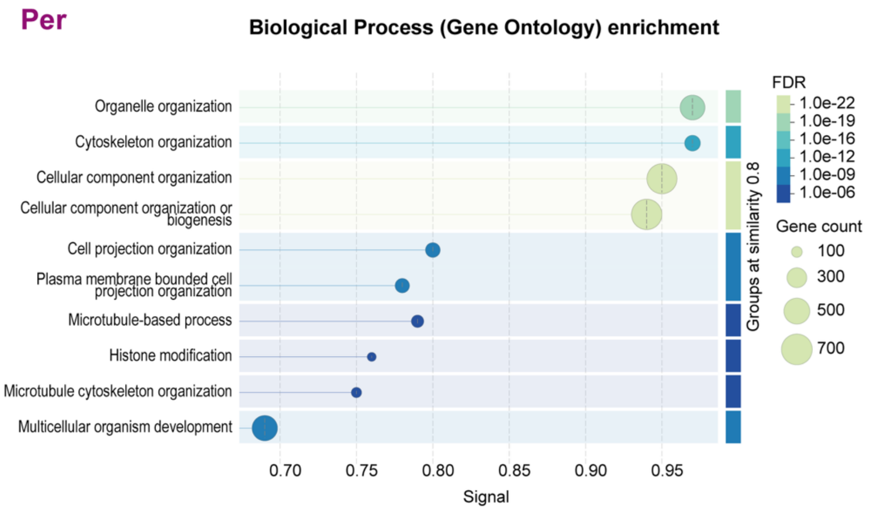


b

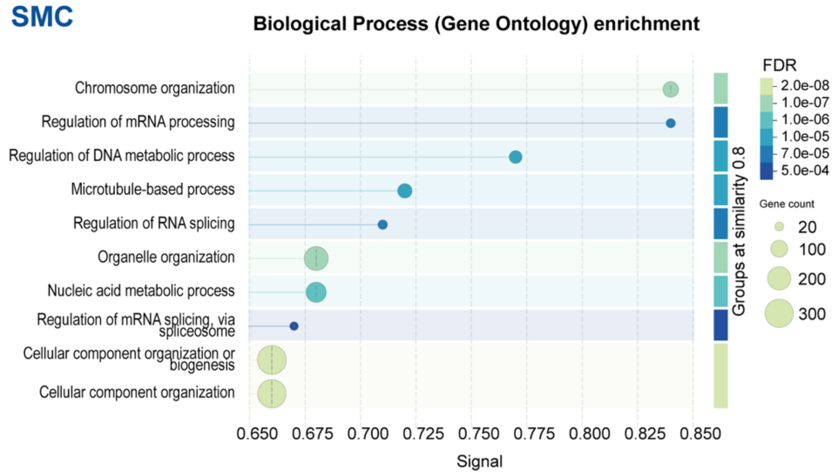


**Supplementary Figure 2. Gene ontology and network analysis of differentially phosphorylated proteins in mural cell-specific Notch3$^{R170C}$ mice.** (a,b) Top Gene Ontology (GO) biological processes enriched among differentially phosphorylated proteins in (a) Per-Notch3$^{R170C}$ mice or (b) SMC-Notch3$^{R170C}$ mice compared to Notch3$^{WT}$ controls. Dot size indicates the number of proteins in each GO term; color scale represents FDR-corrected significance. (b) Equivalent enrichment analysis for SMC-Notch3$^{R170C}$ mice reveals enrichment of nuclear and RNA metabolism–related processes.

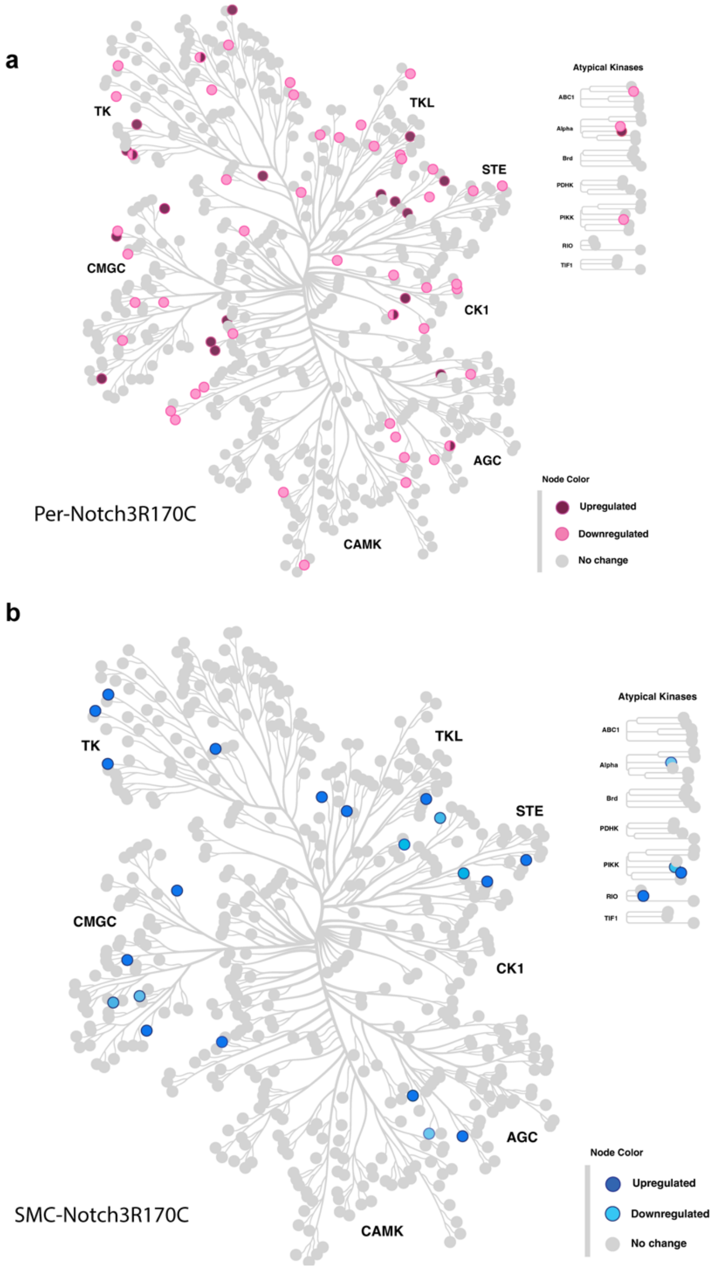


**Supplementary Figure 3. Kinome maps of differentially regulated protein kinases in Per- and SMC-Notch3$^{R170C}$ mice**. Kinases identified in brain vessel proteomes were mapped onto the human kinome phylogenetic tree and colored according to direction of change relative to littermate controls. (a) Per-Notch3$^{R170C}$ mice show a broader distribution of altered kinases across all major kinase groups, including AGC, CAMK, CMGC, CK1, STE, TK, and TKL families. (b) SMC-Notch3$^{R170C}$ mice display fewer changes, with altered kinases distributed across multiple groups. Node color indicates upregulated (pink), downregulated (dark pink in a, light blue in b), or unchanged (grey) kinases; insets show atypical kinase group distribution.

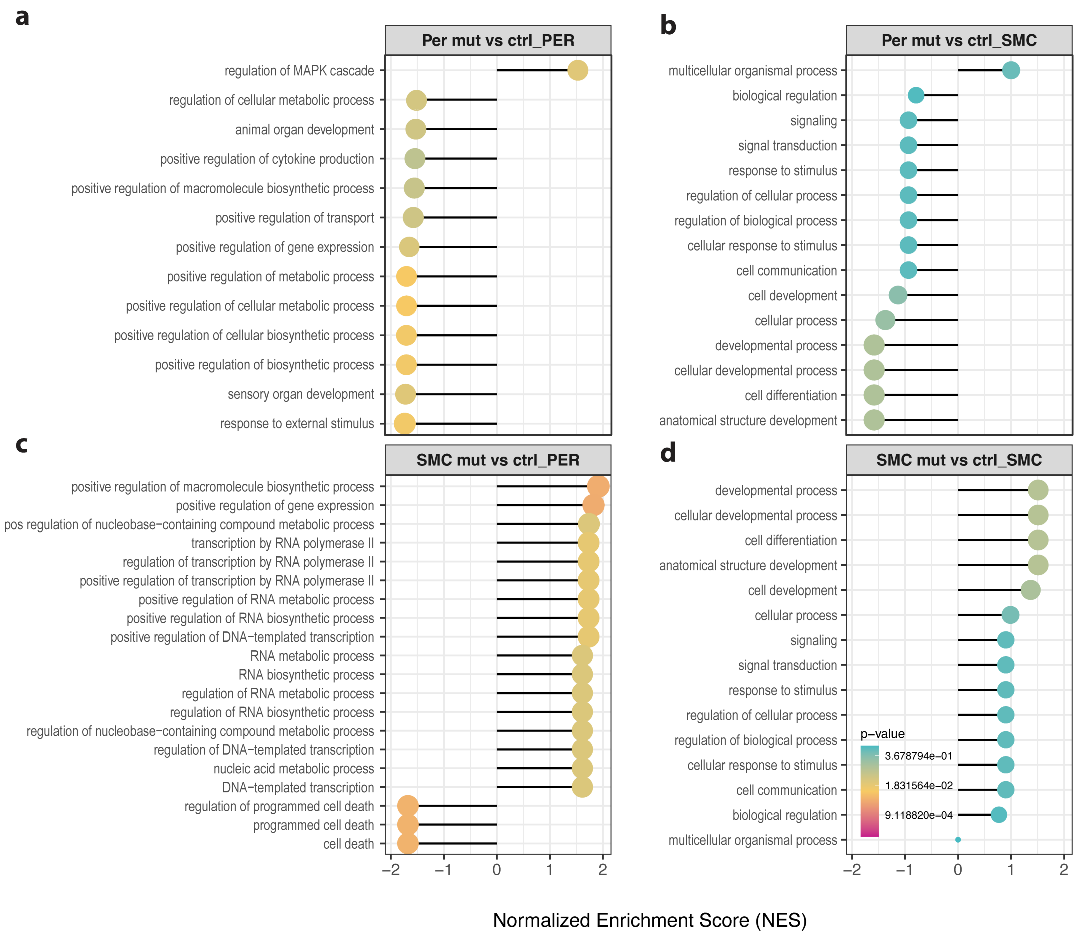


**Supplementary Figure 4. Gene set enrichment analysis (GSEA) cell type–assigned proteins in mouse brain vessel proteomes from Per- and SMC-Notch3$^{R170C}$ mice.** Proteins were annotated to vascular cell types by integration with a single-cell RNA-seq reference atlas. (a) Top enriched biological processes for proteins assigned to pericytes in Per-Notch3$^{R170C}$ vs. control mice. (b) Top enriched biological processes for proteins assigned to smooth muscle cells in Per-Notch3$^{R170C}$ vs. control mice. (c) Top enriched biological processes for proteins assigned to pericytes in SMC-Notch3$^{R170C}$ vs. control mice. (d) Top enriched biological processes for proteins assigned to smooth muscle cells in SMC-Notch3$^{R170C}$ vs. control mice. NES = normalized enrichment score; color scale represents adjusted p-values for selected top pathways.

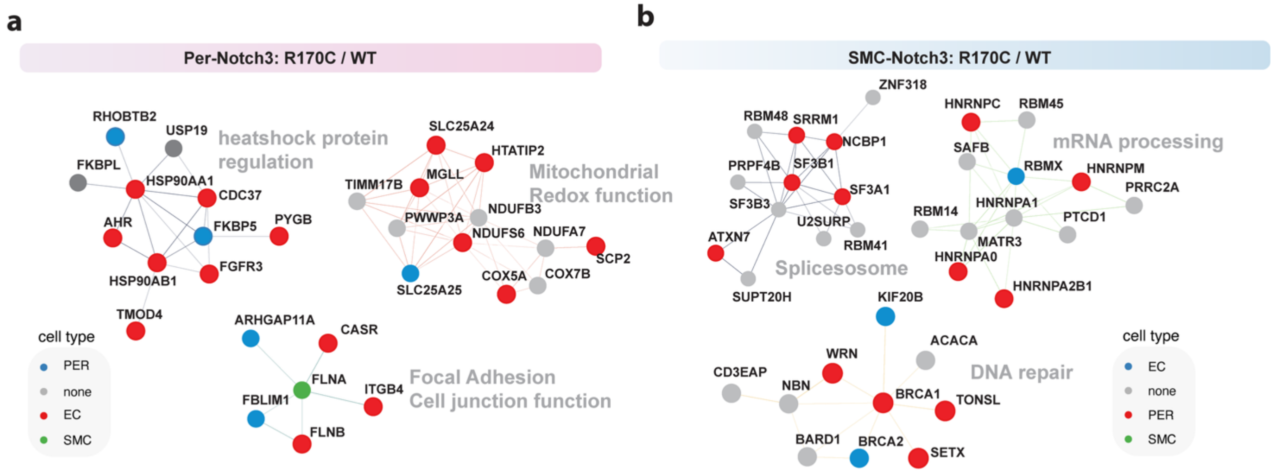


**Supplementary Figure 5. Networks analysis of phosphoproteomic changes in mural cell-specific Notch3$^{R170C}$ mice annotated by predicted vascular cell type.** (a,b) Protein interaction network of differentially phosphorylated proteins in (a) Per-Notch3$^{R170C}$ mice and (b) SMC-Notch3$^{R170C}$ mice, annotated by predicted vascular cell type (EC, endothelial cell, red; PER, pericyte, blue; SMC , smooth muscle cell, green).
