## Supplementary material for "Multiomic profiling reveals pericyte and smooth muscle cell contributions to CADASIL pathology in cell-specific Notch3 mutant mice": Suppl. Data 1

**GeneAgent output (August 2025)**

https://doi.org/10.1038/s41592-025-02748-6

### 1. SMC-NOTCH3 (all vascular cells)

Analysis of gene set: Prl7a2, Tmem69, Ctsc, Iqcf5, Cebpg, Ddx51, Clec1b, Crlf3, Rubcn, Spi1 ...

Step 5: SummarizationProcess: Leukocyte-Mediated Cytotoxicity

Ctsc is essential for activating serine proteases in immune cells, supporting innate immunity and inflammatory responses critical for leukocyte-mediated cytotoxicity. It encodes a lysosomal cysteine proteinase that acts as a central coordinator for the activation of many serine proteinases in immune system cells.

Cebpg and Spi1 are transcription factors that regulate genes involved in myeloid differentiation, immune responses, and immune cell development. Cebpg is part of the C/EBP family, which regulates transcription mediated by CCAAT/enhancer elements, and is associated with leukocyte-mediated cytotoxicity. Spi1 encodes PU.1, an ETS-domain transcription factor that activates gene expression during myeloid and B-lymphoid cell development, contributing to immune cell differentiation.

Rubcn is a negative regulator of autophagy and endocytic trafficking, controlling endosome maturation. It is involved in autophagy regulation, which is essential for immune homeostasis and cellular processes during leukocyte-mediated cytotoxicity.

Mymx is involved in cellular fusion processes, specifically myoblast fusion during skeletal muscle regeneration. This function may have implications for macrophage multinucleation, although direct evidence linking it to leukocyte-mediated cytotoxicity is not established.

Ddx51 is involved in RNA processing and ribosome biogenesis, enabling RNA binding activity and rRNA processing. These processes are essential for protein synthesis in immune cells, although its direct role in leukocyte-mediated cytotoxicity remains unclear.

Clec1b is a C-type lectin-like receptor expressed in myeloid and natural killer cells. It interacts with major histocompatibility complex class I molecules and can modulate cytotoxicity and cytokine secretion, contributing to immune responses.

Irak1bp1 is predicted to enable signaling adaptor activity and is involved in canonical NF-kappaB signal transduction, which may modulate innate immune responses and inflammation. However, its specific role in leukocyte-mediated cytotoxicity is not definitively established.

Tmem69 and Iqcf5 lack direct evidence linking them to immune cell communication or activation during leukocyte-mediated cytotoxicity. Tmem69 is located in the mitochondrion, and Iqcf5 is predicted to enable calmodulin binding activity, which is related to calcium signaling.

Crlf3 is a cytokine receptor-like factor that may negatively regulate cell cycle progression at the G0/G1 phase. Its role in immune responses or leukocyte-mediated cytotoxicity is not clearly defined.

Prl7a2 is expressed in the placenta and involved in pregnancy-related processes, such as positive regulation of lactation and receptor signaling pathways via JAK-STAT. While it may contribute to maternal-fetal immune interactions, there is no direct evidence linking it to leukocyte-mediated cytotoxicity.Step 4: Self-verification of the analytical narrativesOriginal_claim:Prl7a2 is involved in immune modulation, particularly at the maternal-fetal interface, contributing to immune tolerance and regulation in the context of leukocyte-mediated cytotoxicity.

Verified_claim:The claim that Prl7a2 is involved in immune modulation, particularly at the maternal-fetal interface, contributing to immune tolerance and regulation in the context of leukocyte-mediated cytotoxicity is weakly supported.

Evidence:

1. Gene Summary: Prl7a2 is predicted to enable hormone activity and prolactin receptor binding activity. It is involved in processes such as female pregnancy, positive regulation of lactation, and positive regulation of receptor signaling pathways via JAK-STAT. It is expressed in the placenta and other extraembryonic components, which aligns with its potential role at the maternal-fetal interface. However, there is no direct mention of immune modulation or leukocyte-mediated cytotoxicity in the summary.

2. Biological Domains: No specific biological domains related to immune modulation or leukocyte-mediated cytotoxicity were identified for Prl7a2.

3. Diseases: No diseases associated with Prl7a2 were found, which could have provided indirect evidence for its role in immune-related processes.

While the gene's expression in the placenta and involvement in pregnancy-related processes suggest a possible connection to maternal-fetal immune interactions, there is no direct evidence linking Prl7a2 to immune tolerance or leukocyte-mediated cytotoxicity. Thus, the support for the claim is weak and requires further evidence for validation.

Original_claim:Tmem69 and Iqcf5 are implicated in signaling pathways and calcium signaling, respectively, which may influence immune cell communication and activation during leukocyte-mediated cytotoxicity.

Verified_claim:Weak support.

1. TMEM69:

- The gene summary indicates that TMEM69 is located in the mitochondrion, but there is no direct evidence linking it to signaling pathways or immune cell communication and activation during leukocyte-mediated cytotoxicity.

2. IQCF5:

- The gene summary states that IQCF5 is predicted to enable calmodulin binding activity, which is related to calcium signaling. However, there is no direct evidence connecting IQCF5 to immune cell communication and activation during leukocyte-mediated cytotoxicity.

3. Enrichment and Pathway Analysis:

- No enrichment functions or biological pathways were identified for the gene set "Tmem69, Iqcf5," which limits the ability to confirm their roles in signaling pathways or immune-related processes.

4. Biological Domains:

- No specific biological domains were identified for either TMEM69 or IQCF5.

While IQCF5's association with calmodulin binding suggests a potential role in calcium signaling, there is insufficient evidence to confirm the claim that these genes influence immune cell communication and activation during leukocyte-mediated cytotoxicity. The support for the claim is weak and requires further evidence.

Original_claim:Ctsc is essential for activating serine proteases in immune cells, supporting innate immunity and inflammatory responses critical for leukocyte-mediated cytotoxicity.

Verified_claim:The claim that "Ctsc is essential for activating serine proteases in immune cells, supporting innate immunity and inflammatory responses critical for leukocyte-mediated cytotoxicity" is **supported**.

1. Gene Summary Evidence:

- The gene summary for CTSC (cathepsin C) states that it encodes a lysosomal cysteine proteinase that acts as a central coordinator for the activation of many serine proteinases in immune system cells. This aligns with the claim that CTSC is essential for activating serine proteases in immune cells.

- The summary also mentions that defects in CTSC are associated with Papillon-Lefevre syndrome, which involves immune dysfunction, further supporting its role in immune responses.

2. Enrichment Evidence:

- The enrichment analysis for CTSC indicates its association with biological features, though the specific enrichment result provided does not directly address immune or inflammatory responses.

No additional evidence was found regarding specific diseases or biological domains directly linked to CTSC in this context. However, the gene summary provides strong support for the claim.

Original_claim:Cebpg and Spi1 are transcription factors that regulate genes involved in myeloid differentiation, immune responses, and immune cell development, contributing to leukocyte-mediated cytotoxicity.

Verified_claim:The claim that Cebpg and Spi1 are transcription factors that regulate genes involved in myeloid differentiation, immune responses, and immune cell development, contributing to leukocyte-mediated cytotoxicity, is **supported** by the following evidence:

1. **Cebpg (CEBPG)**:

- Cebpg is part of the C/EBP family of transcription factors, which regulate transcription mediated by CCAAT/enhancer elements. These transcription factors are involved in various cellular processes, including immune responses. (Source: Gene summary for CEBPG)

- Enrichment analysis indicates that Cebpg is associated with "leukocyte-mediated cytotoxicity" (GO:0001909), which directly supports its role in immune responses and cytotoxicity.

2. **Spi1 (SPI1)**:

- Spi1 encodes an ETS-domain transcription factor (PU.1) that activates gene expression during myeloid and B-lymphoid cell development. This directly supports its involvement in myeloid differentiation and immune cell development. (Source: Gene summary for SPI1)

- Enrichment analysis shows that Spi1 is involved in "erythrocyte differentiation" (GO:0030218) and "B cell differentiation" (GO:0030183), further supporting its role in immune cell development.

3. **Combined Evidence**:

- Enrichment analysis for the gene set (Cebpg and Spi1) highlights their involvement in "leukocyte-mediated cytotoxicity" (GO:0001909), "erythrocyte differentiation" (GO:0030218), and "B cell differentiation" (GO:0030183). These processes are integral to immune responses and immune cell development.

The evidence collectively supports the claim that Cebpg and Spi1 are transcription factors regulating genes involved in myeloid differentiation, immune responses, and immune cell development, contributing to leukocyte-mediated cytotoxicity.

Original_claim:Ddx51 and Rubcn are involved in RNA processing, ribosome biogenesis, and autophagy regulation, processes that are essential for protein synthesis and antigen presentation in leukocyte-mediated cytotoxicity.

Verified_claim:The claim that Ddx51 and Rubcn are involved in RNA processing, ribosome biogenesis, and autophagy regulation, processes essential for protein synthesis and antigen presentation in leukocyte-mediated cytotoxicity, is partially supported with evidence, but some aspects remain ambiguous.

1. **Ddx51**:

- **RNA Processing**: The gene summary for Ddx51 indicates that it enables RNA binding activity and is predicted to be involved in rRNA processing. This supports its role in RNA processing and ribosome biogenesis.

- **Autophagy Regulation**: No direct evidence was found linking Ddx51 to autophagy regulation.

2. **Rubcn**:

- **Autophagy Regulation**: The gene summary for Rubcn describes it as a negative regulator of autophagy and endocytic trafficking, controlling endosome maturation. This supports its involvement in autophagy regulation.

- **RNA Processing and Ribosome Biogenesis**: No evidence was found linking Rubcn to RNA processing or ribosome biogenesis.

3. **Pathway Analysis**:

- The pathway analysis for the gene set "Ddx51, Rubcn" identified "Autophagy" as a related biological pathway, with Rubcn being the overlapping gene. This further supports Rubcn's role in autophagy regulation.

4. **Protein Synthesis and Antigen Presentation in Leukocyte-Mediated Cytotoxicity**:

- No direct evidence was found linking Ddx51 or Rubcn to protein synthesis or antigen presentation in leukocyte-mediated cytotoxicity.

**Conclusion**: The claim is partially supported. Ddx51 is involved in RNA processing and ribosome biogenesis, while Rubcn is involved in autophagy regulation. However, there is no evidence directly linking these genes to protein synthesis or antigen presentation in leukocyte-mediated cytotoxicity.

Original_claim:Clec1b and Crlf3 are receptors involved in immune responses, including pathogen recognition and signaling, which are relevant to leukocyte-mediated cytotoxicity.

Verified_claim:The claim that Clec1b and Crlf3 are receptors involved in immune responses, including pathogen recognition and signaling, which are relevant to leukocyte-mediated cytotoxicity, is weakly supported.

Evidence:

1. CLEC1B (Clec1b) is described as a C-type lectin-like receptor expressed in myeloid cells and natural killer (NK) cells. It interacts with major histocompatibility complex class I molecules and can either inhibit or activate cytotoxicity and cytokine secretion. This suggests a role in immune responses and leukocyte-mediated cytotoxicity (Source: Gene summary for CLEC1B).

2. CRLF3 (Crlf3) encodes a cytokine receptor-like factor that may negatively regulate cell cycle progression at the G0/G1 phase. There is no direct evidence from the gene summary linking CRLF3 to immune responses, pathogen recognition, or leukocyte-mediated cytotoxicity (Source: Gene summary for CRLF3).

3. No specific biological domains or diseases were identified for either CLEC1B or CRLF3 that directly confirm their involvement in the claimed immune functions.

While CLEC1B has a clear connection to immune responses and cytotoxicity, the evidence for CRLF3's involvement in these processes is lacking. Therefore, the claim is only weakly supported.

Original_claim:Mymx and Rubcn are associated with cellular fusion processes and autophagy regulation, respectively, which may play roles in macrophage multinucleation and immune homeostasis during leukocyte-mediated cytotoxicity.

Verified_claim:The claim that Mymx and Rubcn are associated with cellular fusion processes and autophagy regulation, respectively, is supported by evidence.

1. Mymx (MYMX) is described as "myomixer, myoblast fusion factor" and is involved in myoblast fusion during skeletal muscle regeneration. This aligns with the claim that Mymx is associated with cellular fusion processes.

2. Rubcn (RUBCN) is described as "rubicon autophagy regulator" and is a negative regulator of autophagy and endocytic trafficking. This supports the claim that Rubcn is associated with autophagy regulation.

However, the claim further suggests that these roles may contribute to macrophage multinucleation and immune homeostasis during leukocyte-mediated cytotoxicity. While the gene functions align with the claim, there is no direct evidence provided here linking these genes to macrophage multinucleation or immune homeostasis during leukocyte-mediated cytotoxicity. This part of the claim remains ambiguous and unverified.

Original_claim:Irak1bp1 acts as a negative regulator of Toll-like receptor signaling, modulating innate immune responses and inflammation in the context of leukocyte-mediated cytotoxicity.

Verified_claim:The claim that Irak1bp1 acts as a negative regulator of Toll-like receptor signaling, modulating innate immune responses and inflammation in the context of leukocyte-mediated cytotoxicity is weakly supported.

Evidence:

1. Gene Summary: Irak1bp1 (interleukin 1 receptor associated kinase 1 binding protein 1) is predicted to enable signaling adaptor activity and is involved in canonical NF-kappaB signal transduction. It is located in the cytoplasm and nucleus. While this suggests a role in signaling pathways, it does not explicitly confirm its function as a negative regulator of Toll-like receptor signaling or its involvement in leukocyte-mediated cytotoxicity.

2. Domain Information: No specific biological domains were identified for Irak1bp1, which limits further insights into its functional mechanisms related to Toll-like receptor signaling or immune responses.

Further evidence is required to conclusively verify the claim.

Original_claim:

Verified_claim:Failed.

Step 3: ModificationProcess: Leukocyte Mediated Cytotoxicity

The system of interacting proteins encoded by this gene set appears to play a prominent role in leukocyte-mediated cytotoxicity, as supported by enrichment analysis. This biological process involves the directed killing of target cells by leukocytes, which is critical for immune defense mechanisms.

Prl7a2: Prolactin family member 7A2 is associated with immune modulation, particularly in the maternal-fetal interface, suggesting a role in immune tolerance and regulation.

Tmem69: While its specific function is less characterized, transmembrane proteins often contribute to signaling pathways, potentially influencing immune cell communication or membrane-associated processes.

Ctsc: Cathepsin C is a lysosomal protease critical for activating serine proteases in immune cells, such as neutrophils and mast cells, which are essential for innate immunity and inflammatory responses.

Iqcf5: IQ motif-containing F5 is poorly characterized, but IQ domain proteins often interact with calmodulin, suggesting a potential role in calcium signaling, which is critical for immune cell activation.

Cebpg: CCAAT/enhancer-binding protein gamma is a transcription factor that regulates the expression of genes involved in myeloid differentiation and immune responses, particularly in macrophages and granulocytes.

Ddx51: DEAD-box helicase 51 is involved in RNA processing and ribosome biogenesis, which are essential for the high protein synthesis demands of activated immune cells.

Clec1b: C-type lectin domain family 1 member B is a receptor involved in platelet activation and immune responses, particularly in recognizing pathogens and mediating thrombosis-inflammation interactions.

Crlf3: Cytokine receptor-like factor 3 is implicated in neuronal survival but may also have roles in immune signaling, as cytokine receptors are often multifunctional.

Rubcn: Rubicon is a negative regulator of autophagy and endosomal trafficking, processes critical for antigen presentation and immune homeostasis.

Spi1: SPI1 (PU.1) is a master transcription factor for myeloid and lymphoid lineage commitment, directly regulating genes involved in immune cell development and function.

Mymx: Myomixer is primarily known for its role in muscle cell fusion, but its expression in this context may suggest a secondary, less-characterized role in cellular fusion processes relevant to immune cells, such as macrophage multinucleation.

Irak1bp1: IRAK1-binding protein 1 is a negative regulator of Toll-like receptor (TLR) signaling, which modulates innate immune responses and inflammation.

The collective activity of these proteins suggests a system that integrates transcriptional regulation (Cebpg, Spi1), proteolytic activation (Ctsc), signaling modulation (Irak1bp1, Clec1b), and cellular processes like autophagy (Rubcn) and RNA processing (Ddx51). These functions converge on the regulation of leukocyte-mediated cytotoxicity, which is essential for immune defense and pathogen elimination.Step 2: Self-verification of the process nameOriginal_claim:The entire gene set is involved in immune regulation and myeloid differentiation. Here is the entire gene set used for verification:

##Prl7a2,Tmem69,Ctsc,Iqcf5,Cebpg,Ddx51,Clec1b,Crlf3,Rubcn,Spi1,Mymx,Irak1bp1##

Verified_claim:Failed.

Original_claim:The gene set plays a critical role in the regulation of immune responses and the differentiation of myeloid cells. Here is the entire gene set used for verification:

##Prl7a2,Tmem69,Ctsc,Iqcf5,Cebpg,Ddx51,Clec1b,Crlf3,Rubcn,Spi1,Mymx,Irak1bp1##

Verified_claim:The claim that the gene set plays a critical role in the regulation of immune responses and the differentiation of myeloid cells is **weakly supported** based on the following evidence:

1. **Enrichment Analysis**:

- The gene set is associated with "leukocyte mediated cytotoxicity," which is a process related to immune responses. This provides some evidence of the gene set's involvement in immune regulation.

2. **Pathway Analysis**:

- The gene SPI1 in the set is linked to pathways such as "Type II interferon signaling (interferon-gamma)" and "RANKL signaling pathway," both of which are relevant to immune responses and cellular differentiation.

- The gene CLEC1B is associated with the "GPVI-mediated Activation Cascade," which is involved in platelet activation and immune responses.

3. **Gene-Specific Evidence**:

- SPI1 (PU.1) is a transcription factor that plays a significant role in myeloid and B-lymphoid cell development. It regulates gene expression during the differentiation of these immune cells, supporting the claim regarding myeloid cell differentiation.

While there is evidence linking the gene set to immune regulation and myeloid cell differentiation, the support is not comprehensive for the entire gene set. The evidence primarily revolves around specific genes like SPI1 and CLEC1B, and their roles in immune-related pathways. Therefore, the claim is weakly supported.

Original_claim:The entire gene set contributes to processes related to immune system modulation and myeloid lineage development. Here is the entire gene set used for verification:

##Prl7a2,Tmem69,Ctsc,Iqcf5,Cebpg,Ddx51,Clec1b,Crlf3,Rubcn,Spi1,Mymx,Irak1bp1##

Verified_claim:The claim that the gene set contributes to processes related to immune system modulation and myeloid lineage development is weakly supported based on the following evidence:

1. Enrichment Analysis:

- The gene set is associated with "leukocyte mediated cytotoxicity," which is a process involving immune system modulation. This provides some support for the claim regarding immune system involvement.

2. Pathway Analysis:

- The gene set overlaps with pathways such as "Type II interferon signaling (interferon-gamma)" and "RANKL signaling pathway," both of which are related to immune system processes. Additionally, "GPVI-mediated Activation Cascade" involves CLEC1B, which is relevant to platelet activation and immune responses.

- SPI1, a gene in the set, is involved in pathways related to immune signaling and myeloid lineage development.

3. Gene Summary:

- TMEM69 is located in the mitochondrion, but its specific role in immune system modulation or myeloid lineage development is not directly indicated in the summary.

While there is evidence linking some genes in the set to immune system processes and myeloid lineage development, the association is not comprehensive for the entire gene set. Therefore, the claim is weakly supported.

Original_claim: Here is the entire gene set used for verification:

##Prl7a2,Tmem69,Ctsc,Iqcf5,Cebpg,Ddx51,Clec1b,Crlf3,Rubcn,Spi1,Mymx,Irak1bp1##

Verified_claim:

Decision: Weak support.

Evidences:

1. No complex protocol IDs or corresponding complex names were found for the gene set provided.

2. Enrichment analysis of the gene set revealed a significant association with the biological process "leukocyte mediated cytotoxicity" (GO:0001909), which involves the directed killing of a target cell by a leukocyte. This suggests a potential functional role in immune response or cytotoxic activity.

While the enrichment analysis provides some functional context, the lack of additional supporting evidence (e.g., complex associations) makes the claim only weakly supported. Further investigation or specific claims about the gene set's role would require more detailed evidence.

Step 1: Initial generation for gene setProcess: Immune Regulation and Myeloid Differentiation

The system of interacting proteins encoded by this gene set appears to play a prominent role in immune regulation and myeloid cell differentiation. This conclusion is supported by the functional annotations and known roles of the individual genes.

Prl7a2: Prolactin family member 7A2 is associated with immune modulation, particularly in the maternal-fetal interface, suggesting a role in immune tolerance and regulation.

Tmem69: While its specific function is less characterized, transmembrane proteins often contribute to signaling pathways, potentially influencing immune cell communication or membrane-associated processes.

Ctsc: Cathepsin C is a lysosomal protease critical for activating serine proteases in immune cells, such as neutrophils and mast cells, which are essential for innate immunity and inflammatory responses.

Iqcf5: IQ motif-containing F5 is poorly characterized, but IQ domain proteins often interact with calmodulin, suggesting a potential role in calcium signaling, which is critical for immune cell activation.

Cebpg: CCAAT/enhancer-binding protein gamma is a transcription factor that regulates the expression of genes involved in myeloid differentiation and immune responses, particularly in macrophages and granulocytes.

Ddx51: DEAD-box helicase 51 is involved in RNA processing and ribosome biogenesis, which are essential for the high protein synthesis demands of activated immune cells.

Clec1b: C-type lectin domain family 1 member B is a receptor involved in platelet activation and immune responses, particularly in recognizing pathogens and mediating thrombosis-inflammation interactions.

Crlf3: Cytokine receptor-like factor 3 is implicated in neuronal survival but may also have roles in immune signaling, as cytokine receptors are often multifunctional.

Rubcn: Rubicon is a negative regulator of autophagy and endosomal trafficking, processes critical for antigen presentation and immune homeostasis.

Spi1: SPI1 (PU.1) is a master transcription factor for myeloid and lymphoid lineage commitment, directly regulating genes involved in immune cell development and function.

Mymx: Myomixer is primarily known for its role in muscle cell fusion, but its expression in this context may suggest a secondary, less-characterized role in cellular fusion processes relevant to immune cells, such as macrophage multinucleation.

Irak1bp1: IRAK1-binding protein 1 is a negative regulator of Toll-like receptor (TLR) signaling, which modulates innate immune responses and inflammation.

The collective activity of these proteins suggests a system that integrates transcriptional regulation (Cebpg, Spi1), proteolytic activation (Ctsc), signaling modulation (Irak1bp1, Clec1b), and cellular processes like autophagy (Rubcn) and RNA processing (Ddx51). These functions converge on the regulation of immune responses and the differentiation of myeloid cells, which are critical for both innate and adaptive immunity.

### 2. Per-NOTCH3 (all vascular cells)

Analysis of gene set: Elmo3, Camk2a, Degs1, Pkm, Ptk6, Timm44, Nectin1, Suclg1, Atp6v0c, Ugt2a3 ...

Step 5: SummarizationProcess: Synaptic Signaling and Energy Metabolism Coordination

The system of interacting proteins primarily supports synaptic signaling and energy metabolism coordination, with additional roles in cellular stress responses, structural integrity, protein homeostasis, and intracellular signaling. Below is a critical analysis of the biological processes performed by this system:

Snap25, Dlg4, Rac1, Cdc42, and Nectin1 are involved in synaptic signaling. Snap25 facilitates neurotransmitter release by mediating synaptic vesicle membrane docking and fusion through SNARE complexes. Dlg4 organizes postsynaptic density by forming a multimeric scaffold for clustering receptors, ion channels, and signaling proteins. Rac1 and Cdc42 regulate actin cytoskeleton dynamics, essential for synaptic plasticity, with Rac1 promoting cytoskeletal reorganization and Cdc42 controlling actin polymerization via N-WASP and the Arp2/3 complex. Nectin1 contributes to synapse adhesion and stabilization by organizing adherens junctions and tight junctions.

Pkm, Suclg1, Suclg2, Mdh2, Idh3a, Cox5a, Ndufs8, and Atp5mc2 are involved in energy metabolism. Pkm catalyzes the final step of glycolysis, converting phosphoenolpyruvate to pyruvate. Suclg1 and Suclg2 participate in the TCA cycle, with Suclg1 located in the mitochondrial inner membrane and Suclg2 forming part of the succinate-CoA ligase complex. Mdh2 is active in the mitochondrial matrix and contributes to the TCA cycle, gluconeogenesis, and the malate-aspartate shuttle. Idh3a is involved in isocitrate metabolism and the TCA cycle within the mitochondrion. Cox5a and Ndufs8 are components of the electron transport chain, with Cox5a facilitating cytochrome c to oxygen transport in complex IV and Ndufs8 enabling NADH to ubiquinone transport in complex I. Atp5mc2 drives ATP synthesis through the proton motive force in the mitochondrion.

Grpel1, Dele1, Bcl2l15, and Aifm3 are involved in cellular stress responses and apoptosis regulation. Grpel1 mediates mitochondrial stress signaling by enabling unfolded protein binding and protein import into the mitochondrial matrix. Dele1 facilitates HRI-mediated signaling, mitophagy, and responses to iron ion starvation, acting at the mitochondrial inner and outer membranes. Bcl2l15 regulates apoptotic processes in the cytosol and nucleus. Aifm3 contributes to apoptosis execution and cellular stress responses through oxidoreductase activity at the mitochondrial inner membrane.

Tuba1b, Tuba4a, Map6d1, and Dpysl2 are involved in structural integrity and cytoskeletal dynamics. Tuba1b and Tuba4a are alpha tubulin isoforms essential for microtubule assembly and cytoskeletal organization. Map6d1 stabilizes microtubules, functioning as a calmodulin-regulated neuronal protein. Dpysl2 promotes microtubule assembly and regulates axonal growth, guidance, and polarity, playing a role in neuron development and synaptic signaling.

Vcp, Uchl1, and Psmc6 are involved in protein trafficking and degradation. Vcp mediates protein quality control and ERAD by extracting ubiquitinated proteins from membranes or complexes. Uchl1 functions as a ubiquitin hydrolase, hydrolyzing peptide bonds at the C-terminal glycine of ubiquitin. Psmc6 is part of the 26S proteasome complex, facilitating ATP/ubiquitin-dependent protein degradation.

Camk2a, Ppp3r1, and Ptk6 are involved in calcium signaling and kinase activity. Camk2a is critical for synaptic plasticity, hippocampal long-term potentiation, and spatial learning, functioning as a calcium/calmodulin-dependent protein kinase. Ppp3r1 is part of the calcineurin complex, regulating calcium-dependent signaling cascades. Ptk6 acts as a tyrosine kinase, transducing intracellular signals in epithelial tissues.

In summary, this system of interacting proteins coordinates synaptic signaling, energy metabolism, cellular stress responses, structural integrity, protein homeostasis, and intracellular signaling, collectively supporting neuronal function and adaptability.Step 4: Self-verification of the analytical narrativesOriginal_claim:Snap25, Dlg4, Rac1, Cdc42, and Nectin1 are involved in synaptic signaling, with Snap25 facilitating neurotransmitter release, Dlg4 organizing postsynaptic density, Rac1 and Cdc42 regulating actin cytoskeleton dynamics for synaptic plasticity, and Nectin1 contributing to synapse adhesion and stabilization.

Verified_claim:The claim that Snap25, Dlg4, Rac1, Cdc42, and Nectin1 are involved in synaptic signaling is strongly supported by evidence. Here are the findings:

1. Snap25:

- Summary: Snap25 is a presynaptic plasma membrane protein involved in the regulation of neurotransmitter release. It mediates synaptic vesicle membrane docking and fusion through SNARE complexes, which are essential for neurotransmitter release.

- This supports the claim that Snap25 facilitates neurotransmitter release.

2. Dlg4:

- Summary: Dlg4 encodes a member of the membrane-associated guanylate kinase (MAGUK) family. It interacts at postsynaptic sites to form a multimeric scaffold for clustering receptors, ion channels, and associated signaling proteins.

- This supports the claim that Dlg4 organizes postsynaptic density.

3. Rac1:

- Summary: Rac1 is a GTPase that regulates cytoskeletal reorganization, which is crucial for cellular events such as synaptic plasticity.

- This supports the claim that Rac1 regulates actin cytoskeleton dynamics for synaptic plasticity.

4. Cdc42:

- Summary: Cdc42 is a small GTPase that regulates signaling pathways controlling diverse cellular functions, including actin polymerization through its binding to N-WASP, which activates the Arp2/3 complex.

- This supports the claim that Cdc42 regulates actin cytoskeleton dynamics for synaptic plasticity.

5. Nectin1:

- Summary: Nectin1 is an adhesion protein that plays a role in the organization of adherens junctions and tight junctions, contributing to cell-cell adhesion and stabilization.

- This supports the claim that Nectin1 contributes to synapse adhesion and stabilization.

6. Enrichment Analysis:

- The gene set (Snap25, Dlg4, Rac1, Cdc42, Nectin1) is significantly associated with biological processes such as synapse organization, neuron projection morphogenesis, and cell morphogenesis involved in neuron differentiation. These processes are directly related to synaptic signaling.

Decision: The claim is strongly supported by multiple lines of evidence, including gene summaries and enrichment analysis.

Original_claim:Pkm, Suclg1, Suclg2, Mdh2, Idh3a, Cox5a, Ndufs8, and Atp5mc2 are involved in energy metabolism, with Pkm catalyzing the final step of glycolysis, Suclg1 and Suclg2 participating in the TCA cycle, and Cox5a, Ndufs8, and Atp5mc2 playing roles in oxidative phosphorylation and ATP synthesis.

Verified_claim:The claim that Pkm, Suclg1, Suclg2, Mdh2, Idh3a, Cox5a, Ndufs8, and Atp5mc2 are involved in energy metabolism is strongly supported by the evidence. Here are the findings:

1. **Pkm**: The gene enables pyruvate kinase activity and is involved in canonical glycolysis, specifically catalyzing the final step of glycolysis. This aligns with the claim.

2. **Suclg1 and Suclg2**: Both genes are involved in the tricarboxylic acid (TCA) cycle and succinate metabolic processes. Suclg1 is located in the mitochondrial inner membrane, and Suclg2 is part of the succinate-CoA ligase complex. This supports their roles in the TCA cycle.

3. **Mdh2**: The gene is involved in the TCA cycle, gluconeogenesis, and the malate-aspartate shuttle. It is active in the mitochondrial matrix, confirming its role in energy metabolism.

4. **Idh3a**: The gene is involved in the TCA cycle and isocitrate metabolic processes. It is located in the mitochondrion, supporting its role in energy metabolism.

5. **Cox5a**: The gene is part of respiratory chain complex IV and is involved in mitochondrial electron transport, cytochrome c to oxygen. This supports its role in oxidative phosphorylation.

6. **Ndufs8**: The gene contributes to NADH dehydrogenase activity and is involved in mitochondrial electron transport, NADH to ubiquinone. It is part of respiratory chain complex I, confirming its role in oxidative phosphorylation.

7. **Atp5mc2**: The gene is involved in proton motive force-driven ATP synthesis and is located in the mitochondrion. This supports its role in ATP synthesis.

Decision: The claim is verified and strongly supported by the evidence provided.

Original_claim:Grpel1, Dele1, Bcl2l15, and Aifm3 are involved in cellular stress responses and apoptosis regulation, with Grpel1 and Dele1 mediating mitochondrial stress signaling, and Bcl2l15 and Aifm3 regulating apoptotic pathways to protect cells under stress conditions.

Verified_claim:The claim that Grpel1, Dele1, Bcl2l15, and Aifm3 are involved in cellular stress responses and apoptosis regulation, with Grpel1 and Dele1 mediating mitochondrial stress signaling, and Bcl2l15 and Aifm3 regulating apoptotic pathways to protect cells under stress conditions, is supported by the following evidence:

1. **Grpel1**:

- Summary: Grpel1 (GrpE like 1, mitochondrial) enables identical protein binding activity and unfolded protein binding activity. It is predicted to be involved in protein import into the mitochondrial matrix and is located in the mitochondrial matrix and nucleoplasm. This aligns with its role in mitochondrial stress signaling.

- Enrichment: Grpel1 is associated with the mitochondrial inner membrane, mitochondrial membrane, and mitochondrial envelope, which are critical for mitochondrial stress signaling.

2. **Dele1**:

- Summary: Dele1 (DAP3 binding cell death enhancer 1) enables protein kinase binding activity and protein serine/threonine kinase activator activity. It is involved in HRI-mediated signaling, positive regulation of mitophagy, and response to iron ion starvation. It is located in the mitochondrial inner membrane, cytosol, and mitochondrial outer membrane. These functions support its role in mitochondrial stress signaling.

- Enrichment: Dele1 is also associated with the mitochondrial inner membrane and mitochondrial envelope, further supporting its involvement in mitochondrial stress responses.

3. **Bcl2l15**:

- Summary: Bcl2l15 (BCL2 like 15) is predicted to be involved in the apoptotic process and regulation of apoptotic processes. It is active in the cytosol and nucleus, which supports its role in regulating apoptotic pathways.

- Enrichment: No specific mitochondrial association was found, but its role in apoptosis regulation is consistent with the claim.

4. **Aifm3**:

- Summary: Aifm3 (AIF family member 3) is predicted to enable oxidoreductase activity and is involved in the execution phase of apoptosis. It is located in the cytosol, endoplasmic reticulum, and mitochondrial inner membrane. This supports its role in apoptosis regulation and cellular stress responses.

- Enrichment: Aifm3 is associated with the mitochondrial inner membrane, which is relevant to its role in apoptosis and stress responses.

Decision: The claim is **strongly supported** by the evidence. Grpel1 and Dele1 are involved in mitochondrial stress signaling, while Bcl2l15 and Aifm3 are involved in apoptotic pathways, aligning with the described roles in cellular stress responses and apoptosis regulation.

Original_claim:Tuba1b, Tuba4a, Map6d1, and Dpysl2 are involved in structural integrity and cytoskeletal dynamics, with Tuba1b and Tuba4a contributing to microtubule assembly, Map6d1 stabilizing microtubules, and Dpysl2 regulating axonal growth and guidance.

Verified_claim:The claim that Tuba1b, Tuba4a, Map6d1, and Dpysl2 are involved in structural integrity and cytoskeletal dynamics is strongly supported by the evidence. Here are the findings:

1. Tuba1b:

- Summary: TUBA1B is described as a structural constituent of the cytoskeleton and is involved in microtubule cytoskeleton organization. It is active in the microtubule cytoskeleton, supporting its role in microtubule assembly and structural integrity.

2. Tuba4a:

- Summary: TUBA4A encodes an alpha tubulin that is a major component of microtubules, which are essential for the eukaryotic cytoskeleton. This supports its contribution to microtubule assembly and cytoskeletal dynamics.

3. Map6d1:

- Summary: MAP6D1 is related to STOP proteins and may function as a calmodulin-regulated neuronal protein that binds and stabilizes microtubules. This aligns with the claim that it stabilizes microtubules.

4. Dpysl2:

- Summary: DPYSL2 promotes microtubule assembly and is required for neuron guidance, growth, and polarity. It also plays a role in synaptic signaling and growth cone collapse, supporting its role in regulating axonal growth and guidance.

Decision: The claim is verified and strongly supported by the evidence provided.

Original_claim:Vcp, Uchl1, and Psmc6 are involved in protein trafficking and degradation, with Vcp mediating protein quality control and ERAD, Uchl1 functioning as a ubiquitin hydrolase for protein turnover, and Psmc6 being part of the proteasome complex.

Verified_claim:The claim that Vcp, Uchl1, and Psmc6 are involved in protein trafficking and degradation, with specific roles for each gene, is strongly supported by the evidence.

1. Vcp (valosin containing protein):

- Summary: Vcp is a member of the AAA ATPase family and plays a role in protein degradation, intracellular membrane fusion, and regulation of the cell cycle. It forms a homohexameric complex that extracts ubiquitinated proteins from lipid membranes or protein complexes. This aligns with its role in protein quality control and ERAD (endoplasmic reticulum-associated degradation).

- Enrichment: Vcp is associated with processes like "proteasome-mediated ubiquitin-dependent protein catabolic process," which supports its involvement in protein degradation.

2. Uchl1 (ubiquitin C-terminal hydrolase L1):

- Summary: Uchl1 is a thiol protease that hydrolyzes peptide bonds at the C-terminal glycine of ubiquitin, confirming its function as a ubiquitin hydrolase for protein turnover.

- Enrichment: Uchl1 is linked to "Deubiquitination," further supporting its role in ubiquitin-related processes.

3. Psmc6 (proteasome 26S subunit, ATPase 6):

- Summary: Psmc6 is part of the 26S proteasome complex, which is involved in ATP/ubiquitin-dependent protein degradation. This supports its role as part of the proteasome complex.

- Enrichment: Psmc6 is associated with the "proteasome complex" and "proteasome-mediated ubiquitin-dependent protein catabolic process," confirming its involvement in protein degradation.

Overall, the evidence from gene summaries and enrichment analysis strongly supports the claim regarding the roles of Vcp, Uchl1, and Psmc6 in protein trafficking and degradation.

Original_claim:Camk2a, Ppp3r1, and Ptk6 are involved in calcium signaling and kinase activity, with Camk2a playing a role in synaptic plasticity, Ppp3r1 being part of calcineurin for calcium-dependent signaling, and Ptk6 functioning as a tyrosine kinase in signal transduction.

Verified_claim:The claim is strongly supported by the evidence provided.

1. Camk2a:

- The summary for CAMK2A indicates that it belongs to the serine/threonine protein kinases family and the Ca(2+)/calmodulin-dependent protein kinases subfamily. It is involved in calcium signaling, which is crucial for synaptic plasticity at glutamatergic synapses. The alpha chain encoded by this gene is required for hippocampal long-term potentiation (LTP) and spatial learning, supporting its role in synaptic plasticity.

2. Ppp3r1:

- The summary for PPP3R1 shows that it is part of the calcineurin complex and is involved in the calcineurin-NFAT signaling cascade, which is calcium-dependent. This supports the claim that PPP3R1 is part of calcineurin for calcium-dependent signaling.

3. Ptk6:

- The summary for PTK6 describes it as a cytoplasmic nonreceptor protein tyrosine kinase that functions as an intracellular signal transducer in epithelial tissues. This aligns with the claim that PTK6 functions as a tyrosine kinase in signal transduction.

Original_claim:

Verified_claim:Failed.

Step 3: ModificationProcess: Synaptic Signaling and Energy Metabolism Coordination

The system of interacting proteins primarily supports synaptic signaling and energy metabolism coordination, with additional roles in cellular stress responses and structural integrity. Below is a critical analysis of the biological processes performed by this system:

1. Synaptic Signaling

Proteins such as Snap25, Dlg4, Rac1, Cdc42, and Nectin1 are central to synaptic signaling and neuronal communication. Snap25 and Dlg4 are involved in neurotransmitter release and postsynaptic density organization, respectively. Rac1 and Cdc42 regulate actin cytoskeleton dynamics, essential for synaptic plasticity and dendritic spine formation. Nectin1 contributes to synapse adhesion and stabilization. These proteins collectively ensure efficient signal transmission and structural maintenance of synapses.

Reasoning: Synaptic signaling requires precise coordination of neurotransmitter release, receptor localization, and cytoskeletal remodeling. The presence of these genes indicates a focus on maintaining synaptic integrity and adaptability.

2. Energy Metabolism

Proteins such as Pkm, Suclg1, Suclg2, Mdh2, Idh3a, Cox5a, Ndufs8, and Atp5mc2 are involved in glycolysis, the tricarboxylic acid (TCA) cycle, and oxidative phosphorylation. Pkm catalyzes the final step of glycolysis, while Suclg1 and Suclg2 are components of the TCA cycle. Cox5a and Ndufs8 are integral to the electron transport chain, and Atp5mc2 is part of ATP synthase, which generates cellular energy. These proteins ensure efficient energy production to meet the high metabolic demands of neurons.

Reasoning: Neurons require substantial energy for synaptic signaling and maintenance. The presence of these metabolic genes highlights the system's role in energy generation and utilization.

3. Stress Response and Apoptosis Regulation

Proteins such as Grpel1, Dele1, Bcl2l15, and Aifm3 are involved in cellular stress responses and apoptosis regulation. Grpel1 and Dele1 mediate mitochondrial stress signaling, while Bcl2l15 and Aifm3 regulate apoptotic pathways. These proteins protect cells from damage and ensure survival under stress conditions.

Reasoning: Neurons are highly sensitive to stress, and mechanisms to mitigate damage are critical for their function and longevity. These genes suggest a role in maintaining cellular homeostasis.

4. Structural Integrity and Cytoskeletal Dynamics

Proteins such as Tuba1b, Tuba4a, Map6d1, and Dpysl2 are involved in microtubule organization and cytoskeletal dynamics. Tuba1b and Tuba4a are tubulin isoforms essential for microtubule assembly, while Map6d1 stabilizes microtubules. Dpysl2 regulates axonal growth and guidance. These proteins contribute to neuronal structure and intracellular transport.

Reasoning: Proper cytoskeletal organization is essential for neuronal morphology, intracellular trafficking, and synaptic function. These genes indicate a role in maintaining structural integrity.

5. Protein Trafficking and Degradation

Proteins such as Vcp, Uchl1, and Psmc6 are involved in protein trafficking and degradation. Vcp mediates protein quality control and endoplasmic reticulum-associated degradation (ERAD). Uchl1 is a ubiquitin hydrolase critical for protein turnover, and Psmc6 is part of the proteasome complex. These proteins ensure proper protein homeostasis.

Reasoning: Neurons rely on efficient protein turnover to maintain function and prevent aggregation. These genes highlight the system's role in protein quality control.

6. Calcium Signaling and Kinase Activity

Proteins such as Camk2a, Ppp3r1, and Ptk6 are involved in calcium signaling and kinase activity. Camk2a is a calcium/calmodulin-dependent kinase critical for synaptic plasticity. Ppp3r1 is part of calcineurin, which regulates calcium-dependent signaling pathways. Ptk6 is a tyrosine kinase involved in signal transduction. These proteins modulate intracellular signaling cascades.

Reasoning: Calcium signaling is essential for synaptic activity and plasticity. The presence of these genes indicates a role in regulating intracellular signaling.

In summary, this system of interacting proteins coordinates synaptic signaling, energy metabolism, stress responses, structural integrity, protein homeostasis, and intracellular signaling. These processes collectively support neuronal function and adaptability.Step 2: Self-verification of the process nameOriginal_claim:The entire gene set is involved in synaptic signaling and energy metabolism coordination. Here is the entire gene set used for verification:

##Elmo3,Camk2a,Degs1,Pkm,Ptk6,Timm44,Nectin1,Suclg1,Atp6v0c,Ugt2a3,Grpel1,Bcl2l15,Tm4sf1,Il10,Prrt2,Vcp,Dpysl2,Nxt1,Ppp1cc,Nptn,Tssk1b,Aspa,Clta,Cldn11,Tas2r109,Tppp,Mboat1,Cend1,Stxbp1,Eif3f,Mrpl35,Dele1,Anapc15,Tspan7,Mtg1,Atp5pf,Riiad1,Tmprss9,H2aj,Grb2,Aldh5a1,Ndufs8,Snrpg,Phb2,Pusl1,Gspt1,Gabarapl1,Chchd2,Psmc6,Ldaf1,Snap25,Dlg4,Nherf1,Uchl1,Isg20,Slc1a4,Gpm6b,Cdc42,Atp5mc2,Cnrip1,Ywhag,Atp6v0d1,Acyp1,Maz,Map6d1,Slco1a6,Dlx4,Prdx6,Myl1,Hspe1,Tcl1a,Suclg2,Ndufa4,Araf,Ptgs1,Efhc1,Mapk7,Sptlc1,Mdh2,Tuba1b,Mamstr,Ppp3r1,Dok6,Rac1,Cox5a,Creb3l3,Bhlha9,Sltm,Tnnc1,Heatr5a,Nrgn,Aifm3,Gnao1,Idh3a,Cwf19l1,Kcnj10,Gpr182,Atp5if1,Ap2s1,Slc66a2,Gpr108,Cox20,Tuba4a,Slc25a3,Farsa##

Verified_claim:The claim that the entire gene set is involved in synaptic signaling and energy metabolism coordination is strongly supported by the evidence.

1. Enrichment Analysis:

- The gene set is significantly associated with "mitochondrion" (GO:0005739), which is a key organelle involved in energy metabolism.

- It is also linked to "aerobic respiration" (GO:0009060) and "generation of precursor metabolites and energy" (GO:0006091), both of which are central processes in energy metabolism.

2. Pathway Analysis:

- The gene set overlaps with pathways such as "Oxidative Phosphorylation," "Citric Acid (TCA) Cycle And Respiratory Electron Transport," and "Pathways of neurodegeneration," which are directly related to energy metabolism and synaptic signaling.

- Specific genes like ATP5PF, NDUFA4, COX5A, and others are involved in oxidative phosphorylation and the TCA cycle, which are critical for energy production.

- Pathways of neurodegeneration and Huntington disease also suggest involvement in synaptic signaling, as these pathways are often associated with neuronal function and signaling.

3. Disease and Functional Context:

- The gene set is associated with diseases like Huntington disease and Amyotrophic lateral sclerosis, which are known to involve synaptic signaling dysfunction and energy metabolism impairments.

Overall, the evidence from enrichment and pathway analyses, as well as disease associations, strongly supports the claim that the gene set is involved in synaptic signaling and energy metabolism coordination.

Original_claim:The gene set plays a critical role in the regulation of synaptic signaling and energy metabolism. Here is the entire gene set used for verification:

##Elmo3,Camk2a,Degs1,Pkm,Ptk6,Timm44,Nectin1,Suclg1,Atp6v0c,Ugt2a3,Grpel1,Bcl2l15,Tm4sf1,Il10,Prrt2,Vcp,Dpysl2,Nxt1,Ppp1cc,Nptn,Tssk1b,Aspa,Clta,Cldn11,Tas2r109,Tppp,Mboat1,Cend1,Stxbp1,Eif3f,Mrpl35,Dele1,Anapc15,Tspan7,Mtg1,Atp5pf,Riiad1,Tmprss9,H2aj,Grb2,Aldh5a1,Ndufs8,Snrpg,Phb2,Pusl1,Gspt1,Gabarapl1,Chchd2,Psmc6,Ldaf1,Snap25,Dlg4,Nherf1,Uchl1,Isg20,Slc1a4,Gpm6b,Cdc42,Atp5mc2,Cnrip1,Ywhag,Atp6v0d1,Acyp1,Maz,Map6d1,Slco1a6,Dlx4,Prdx6,Myl1,Hspe1,Tcl1a,Suclg2,Ndufa4,Araf,Ptgs1,Efhc1,Mapk7,Sptlc1,Mdh2,Tuba1b,Mamstr,Ppp3r1,Dok6,Rac1,Cox5a,Creb3l3,Bhlha9,Sltm,Tnnc1,Heatr5a,Nrgn,Aifm3,Gnao1,Idh3a,Cwf19l1,Kcnj10,Gpr182,Atp5if1,Ap2s1,Slc66a2,Gpr108,Cox20,Tuba4a,Slc25a3,Farsa##

Verified_claim:The claim that the gene set plays a critical role in the regulation of synaptic signaling and energy metabolism is **strongly supported** by the evidence.

1. **Energy Metabolism**:

- The enrichment analysis identified significant associations with terms related to energy metabolism, such as "aerobic respiration" (p-value: 6.11e-06) and "generation of precursor metabolites and energy" (p-value: 1.69e-05). These terms directly relate to the processes of energy production and metabolism.

- Pathway analysis revealed associations with "Oxidative Phosphorylation" and "Citric Acid (TCA) Cycle And Respiratory Electron Transport," both of which are central to cellular energy metabolism.

2. **Synaptic Signaling**:

- The pathway analysis highlighted "Pathways of neurodegeneration" and "Huntington disease," which involve synaptic signaling and neuronal function. Genes such as CAMK2A, DLG4, and RAC1 are known to play roles in synaptic signaling and plasticity.

- The enrichment analysis also identified "Huntington disease" as a significant term (p-value: 1.78e-05), which is associated with synaptic dysfunction.

These findings collectively indicate that the gene set is involved in both energy metabolism and synaptic signaling, aligning with the claim.

Original_claim:The entire gene set contributes to the coordination between synaptic signaling and energy metabolism processes. Here is the entire gene set used for verification:

##Elmo3,Camk2a,Degs1,Pkm,Ptk6,Timm44,Nectin1,Suclg1,Atp6v0c,Ugt2a3,Grpel1,Bcl2l15,Tm4sf1,Il10,Prrt2,Vcp,Dpysl2,Nxt1,Ppp1cc,Nptn,Tssk1b,Aspa,Clta,Cldn11,Tas2r109,Tppp,Mboat1,Cend1,Stxbp1,Eif3f,Mrpl35,Dele1,Anapc15,Tspan7,Mtg1,Atp5pf,Riiad1,Tmprss9,H2aj,Grb2,Aldh5a1,Ndufs8,Snrpg,Phb2,Pusl1,Gspt1,Gabarapl1,Chchd2,Psmc6,Ldaf1,Snap25,Dlg4,Nherf1,Uchl1,Isg20,Slc1a4,Gpm6b,Cdc42,Atp5mc2,Cnrip1,Ywhag,Atp6v0d1,Acyp1,Maz,Map6d1,Slco1a6,Dlx4,Prdx6,Myl1,Hspe1,Tcl1a,Suclg2,Ndufa4,Araf,Ptgs1,Efhc1,Mapk7,Sptlc1,Mdh2,Tuba1b,Mamstr,Ppp3r1,Dok6,Rac1,Cox5a,Creb3l3,Bhlha9,Sltm,Tnnc1,Heatr5a,Nrgn,Aifm3,Gnao1,Idh3a,Cwf19l1,Kcnj10,Gpr182,Atp5if1,Ap2s1,Slc66a2,Gpr108,Cox20,Tuba4a,Slc25a3,Farsa##

Verified_claim:The claim that the entire gene set contributes to the coordination between synaptic signaling and energy metabolism processes is supported by multiple pieces of evidence.

1. Enrichment Analysis:

- The gene set is significantly associated with "mitochondrion" (GO:0005739), which is a key organelle involved in energy metabolism.

- It is also linked to "aerobic respiration" (GO:0009060) and "generation of precursor metabolites and energy" (GO:0006091), both of which are central to energy metabolism processes.

2. Pathway Analysis:

- The gene set overlaps with pathways such as "Oxidative Phosphorylation" and "Citric Acid (TCA) Cycle And Respiratory Electron Transport," which are critical for energy metabolism.

- It is also associated with "Pathways of neurodegeneration" and "Huntington disease," which involve synaptic signaling and energy metabolism dysfunction.

3. Specific Genes:

- Genes like ATP5PF, NDUFA4, COX5A, and others are involved in mitochondrial function and energy production.

- Genes such as CAMK2A and DLG4 are known to play roles in synaptic signaling.

Decision: Strong support for the claim. The gene set is significantly involved in processes and pathways that coordinate synaptic signaling and energy metabolism.

Original_claim: Here is the entire gene set used for verification:

##Elmo3,Camk2a,Degs1,Pkm,Ptk6,Timm44,Nectin1,Suclg1,Atp6v0c,Ugt2a3,Grpel1,Bcl2l15,Tm4sf1,Il10,Prrt2,Vcp,Dpysl2,Nxt1,Ppp1cc,Nptn,Tssk1b,Aspa,Clta,Cldn11,Tas2r109,Tppp,Mboat1,Cend1,Stxbp1,Eif3f,Mrpl35,Dele1,Anapc15,Tspan7,Mtg1,Atp5pf,Riiad1,Tmprss9,H2aj,Grb2,Aldh5a1,Ndufs8,Snrpg,Phb2,Pusl1,Gspt1,Gabarapl1,Chchd2,Psmc6,Ldaf1,Snap25,Dlg4,Nherf1,Uchl1,Isg20,Slc1a4,Gpm6b,Cdc42,Atp5mc2,Cnrip1,Ywhag,Atp6v0d1,Acyp1,Maz,Map6d1,Slco1a6,Dlx4,Prdx6,Myl1,Hspe1,Tcl1a,Suclg2,Ndufa4,Araf,Ptgs1,Efhc1,Mapk7,Sptlc1,Mdh2,Tuba1b,Mamstr,Ppp3r1,Dok6,Rac1,Cox5a,Creb3l3,Bhlha9,Sltm,Tnnc1,Heatr5a,Nrgn,Aifm3,Gnao1,Idh3a,Cwf19l1,Kcnj10,Gpr182,Atp5if1,Ap2s1,Slc66a2,Gpr108,Cox20,Tuba4a,Slc25a3,Farsa##

Verified_claim:The gene set provided has significant associations with several biological processes and cellular components. Here are the findings:

1. **Mitochondrion (GO:0005739)**:

- Description: A semiautonomous, self-replicating organelle involved in tissue respiration.

- Intersection Size: 33 genes from the set are associated with this term.

- P-value: 6.20e-10, indicating strong statistical significance.

- Precision: 0.33, suggesting moderate relevance.

2. **Cytoplasm (GO:0005737)**:

- Description: The contents of a cell excluding the plasma membrane and nucleus.

- Intersection Size: 84 genes from the set are associated with this term.

- P-value: 1.27e-06, indicating strong statistical significance.

- Precision: 0.84, suggesting high relevance.

3. **Aerobic Respiration (GO:0009060)**:

- Description: Enzymatic release of energy requiring oxygen as the terminal electron acceptor.

- Intersection Size: 11 genes from the set are associated with this term.

- P-value: 6.11e-06, indicating strong statistical significance.

- Precision: 0.11, suggesting low relevance.

4. **Generation of Precursor Metabolites and Energy (GO:0006091)**:

- Description: Formation of precursor metabolites and liberation of energy.

- Intersection Size: 15 genes from the set are associated with this term.

- P-value: 1.69e-05, indicating strong statistical significance.

- Precision: 0.15, suggesting low relevance.

5. **Huntington Disease (KEGG:05016)**:

- Description: A neurodegenerative disorder.

- Intersection Size: 13 genes from the set are associated with this pathway.

- P-value: 1.78e-05, indicating strong statistical significance.

- Precision: 0.21, suggesting moderate relevance.

Decision: The gene set is significantly associated with mitochondrial function, cytoplasmic processes, energy metabolism, and Huntington disease pathways. These findings provide strong support for claims related to these biological contexts.

Step 1: Initial generation for gene setProcess: Synaptic Signaling and Energy Metabolism Coordination

The system of interacting proteins primarily supports synaptic signaling and energy metabolism coordination, with additional roles in cellular stress responses and structural integrity. Below is a critical analysis of the biological processes performed by this system:

1. **Synaptic Signaling**

Proteins such as Snap25, Dlg4, Rac1, Cdc42, and Nectin1 are central to synaptic signaling and neuronal communication. Snap25 and Dlg4 are involved in neurotransmitter release and postsynaptic density organization, respectively. Rac1 and Cdc42 regulate actin cytoskeleton dynamics, essential for synaptic plasticity and dendritic spine formation. Nectin1 contributes to synapse adhesion and stabilization. These proteins collectively ensure efficient signal transmission and structural maintenance of synapses.

Reasoning: Synaptic signaling requires precise coordination of neurotransmitter release, receptor localization, and cytoskeletal remodeling. The presence of these genes indicates a focus on maintaining synaptic integrity and adaptability.

2. **Energy Metabolism**

Proteins such as Pkm, Suclg1, Suclg2, Mdh2, Idh3a, Cox5a, Ndufs8, and Atp5mc2 are involved in glycolysis, the tricarboxylic acid (TCA) cycle, and oxidative phosphorylation. Pkm catalyzes the final step of glycolysis, while Suclg1 and Suclg2 are components of the TCA cycle. Cox5a and Ndufs8 are integral to the electron transport chain, and Atp5mc2 is part of ATP synthase, which generates cellular energy. These proteins ensure efficient energy production to meet the high metabolic demands of neurons.

Reasoning: Neurons require substantial energy for synaptic signaling and maintenance. The presence of these metabolic genes highlights the system's role in energy generation and utilization.

3. **Stress Response and Apoptosis Regulation**

Proteins such as Grpel1, Dele1, Bcl2l15, and Aifm3 are involved in cellular stress responses and apoptosis regulation. Grpel1 and Dele1 mediate mitochondrial stress signaling, while Bcl2l15 and Aifm3 regulate apoptotic pathways. These proteins protect cells from damage and ensure survival under stress conditions.

Reasoning: Neurons are highly sensitive to stress, and mechanisms to mitigate damage are critical for their function and longevity. These genes suggest a role in maintaining cellular homeostasis.

4. **Structural Integrity and Cytoskeletal Dynamics**

Proteins such as Tuba1b, Tuba4a, Map6d1, and Dpysl2 are involved in microtubule organization and cytoskeletal dynamics. Tuba1b and Tuba4a are tubulin isoforms essential for microtubule assembly, while Map6d1 stabilizes microtubules. Dpysl2 regulates axonal growth and guidance. These proteins contribute to neuronal structure and intracellular transport.

Reasoning: Proper cytoskeletal organization is essential for neuronal morphology, intracellular trafficking, and synaptic function. These genes indicate a role in maintaining structural integrity.

5. **Protein Trafficking and Degradation**

Proteins such as Vcp, Uchl1, and Psmc6 are involved in protein trafficking and degradation. Vcp mediates protein quality control and endoplasmic reticulum-associated degradation (ERAD). Uchl1 is a ubiquitin hydrolase critical for protein turnover, and Psmc6 is part of the proteasome complex. These proteins ensure proper protein homeostasis.

Reasoning: Neurons rely on efficient protein turnover to maintain function and prevent aggregation. These genes highlight the system's role in protein quality control.

6. **Calcium Signaling and Kinase Activity**

Proteins such as Camk2a, Ppp3r1, and Ptk6 are involved in calcium signaling and kinase activity. Camk2a is a calcium/calmodulin-dependent kinase critical for synaptic plasticity. Ppp3r1 is part of calcineurin, which regulates calcium-dependent signaling pathways. Ptk6 is a tyrosine kinase involved in signal transduction. These proteins modulate intracellular signaling cascades.

Reasoning: Calcium signaling is essential for synaptic activity and plasticity. The presence of these genes indicates a role in regulating intracellular signaling.

In summary, this system of interacting proteins coordinates synaptic signaling, energy metabolism, stress responses, structural integrity, protein homeostasis, and intracellular signaling. These processes collectively support neuronal function and adaptability.

### 3. SMC-NOTCH3 (EC only)

Analysis of gene set: Ctsc, Cebpg, Ido1, Slc11a2Step 5: SummarizationProcess: Immune Regulation and Iron Metabolism

The system of interacting proteins encoded by the genes Ctsc, Cebpg, Ido1, and Slc11a2 contributes to immune regulation and iron metabolism, with distinct roles for each gene in these processes.

Ctsc encodes cathepsin C, a lysosomal cysteine protease that acts as a central coordinator for the activation of serine proteases in immune cells. These proteases, such as neutrophil elastase and cathepsin G, are critical for innate immunity, pathogen clearance, and tissue remodeling during inflammation. Defects in Ctsc are associated with immune-related conditions such as Papillon-Lefevre syndrome, further supporting its role in immune regulation.

Cebpg encodes CCAAT/enhancer-binding protein gamma, a transcription factor that regulates genes involved in immune responses and hematopoiesis. It is part of the C/EBP family, which modulates transcription mediated by CCAAT/enhancer elements. Cebpg is also implicated in leukocyte-mediated cytotoxicity, highlighting its importance in immune system function.

Ido1 encodes indoleamine 2,3-dioxygenase 1, a heme enzyme that catalyzes the first step in the kynurenine pathway of tryptophan catabolism. This pathway is essential for immune tolerance, as it depletes tryptophan and generates metabolites that suppress T-cell proliferation and promote regulatory T-cell differentiation. Ido1 plays a key role in immunoregulation and modulating T-cell behavior, particularly during inflammation.

Slc11a2 encodes solute carrier family 11 member 2, also known as DMT1, a divalent metal transporter responsible for iron uptake and homeostasis. Iron is critical for numerous cellular processes, including heme synthesis and immune cell function. Slc11a2 ensures proper iron absorption and transport, which is essential for maintaining iron balance and preventing oxidative stress or pathogen exploitation.

In summary, this system of proteins collectively contributes to immune regulation and iron metabolism, with Ctsc and Cebpg primarily involved in immune responses, and Ido1 and Slc11a2 playing roles in immune modulation and iron homeostasis.Step 4: Self-verification of the analytical narrativesOriginal_claim:The genes Ctsc, Cebpg, Ido1, and Slc11a2 collectively contribute to the regulation of the paraferritin complex and tryptophan catabolism, which are essential for maintaining iron homeostasis and modulating immune responses.

Verified_claim:The claim that the genes Ctsc, Cebpg, Ido1, and Slc11a2 collectively contribute to the regulation of the paraferritin complex and tryptophan catabolism, which are essential for maintaining iron homeostasis and modulating immune responses, is partially supported by the evidence.

1. **Paraferritin Complex**:

- The enrichment analysis indicates that the paraferritin complex is associated with the gene set. The paraferritin complex is described as being involved in iron transport and is essential for iron homeostasis. However, the specific contribution of each gene (Ctsc, Cebpg, Ido1, Slc11a2) to this complex is not explicitly detailed in the evidence.

2. **Tryptophan Catabolism**:

- The pathway analysis shows that Ido1 is directly involved in tryptophan catabolism, as evidenced by its association with pathways like "Tryptophan catabolism" (BioPlanet_2019) and "Tryptophan Catabolism R-HSA-71240" (Reactome_2022). This supports the claim that tryptophan catabolism is relevant to the gene set.

3. **Iron Homeostasis and Immune Responses**:

- While the paraferritin complex is linked to iron homeostasis, and tryptophan catabolism is known to modulate immune responses, the evidence does not explicitly connect all four genes collectively to these processes. The role of Slc11a2 in iron transport and the involvement of Cebpg in heme deficiency response (Reactome_2022) suggest potential relevance, but the evidence is not comprehensive for all genes.

4. **Interactions**:

- No direct interactions among the genes in the set were identified, which limits the ability to confirm their collective contribution to the processes mentioned.

Decision: The claim is weakly supported. While there is evidence linking individual genes to the paraferritin complex, tryptophan catabolism, iron homeostasis, and immune responses, the collective contribution of all four genes to these processes is not conclusively demonstrated.

Original_claim:Ctsc and Cebpg are involved in immune regulation, with Ctsc encoding a lysosomal cysteine protease that activates serine proteases critical for innate immunity, and Cebpg encoding a transcription factor that regulates genes involved in immune responses and hematopoiesis.

Verified_claim:The claim that Ctsc and Cebpg are involved in immune regulation is strongly supported by evidence.

1. Ctsc (cathepsin C) is described as a lysosomal cysteine proteinase that acts as a central coordinator for the activation of many serine proteinases in immune system cells. This aligns with the claim that Ctsc activates serine proteases critical for innate immunity. Additionally, defects in Ctsc are associated with Papillon-Lefevre syndrome, which involves immune-related symptoms such as periodontitis.

2. Cebpg (CCAAT enhancer binding protein gamma) belongs to the C/EBP family of transcription factors, which regulate viral and cellular transcription mediated by CCAAT/enhancer elements. These transcription factors are involved in immune responses and hematopoiesis, supporting the claim that Cebpg regulates genes involved in immune responses and hematopoiesis.

3. Enrichment analysis for the gene set Ctsc and Cebpg reveals significant involvement in "leukocyte mediated cytotoxicity," a biological process directly related to immune regulation.

The evidence collectively supports the claim that Ctsc and Cebpg are involved in immune regulation, with their respective roles in activating serine proteases and regulating immune-related genes.

Original_claim:Ido1 and Slc11a2 play roles in immune modulation and iron metabolism, with Ido1 catalyzing the first step in the kynurenine pathway of tryptophan catabolism to regulate immune tolerance, and Slc11a2 encoding a divalent metal transporter critical for iron uptake and homeostasis.

Verified_claim:The claim is **supported** by the evidence.

1. For **Ido1**:

The gene summary confirms that Ido1 encodes indoleamine 2,3-dioxygenase 1, which is a heme enzyme catalyzing the first and rate-limiting step in tryptophan catabolism to N-formyl-kynurenine. This enzyme is involved in immunoregulation and modulates T-cell behavior, supporting its role in immune modulation and the kynurenine pathway of tryptophan catabolism.

2. For **Slc11a2**:

The gene summary confirms that Slc11a2 encodes a member of the solute carrier family 11 protein family, which transports divalent metals and is involved in iron absorption. This supports its role as a divalent metal transporter critical for iron uptake and homeostasis.

Original_claim:

Verified_claim:Failed.

Step 3: ModificationProcess: Paraferritin Complex and Tryptophan Catabolism

The system of interacting proteins encoded by the genes Ctsc, Cebpg, Ido1, and Slc11a2 appears to coordinate processes related to the paraferritin complex and tryptophan catabolism. These processes are critical for maintaining iron homeostasis and modulating immune responses.

Ctsc encodes cathepsin C, a lysosomal cysteine protease involved in the activation of serine proteases in immune cells. This protein plays a key role in regulating the activity of neutrophil elastase, cathepsin G, and other proteases critical for innate immune responses. Its function is essential for pathogen clearance and tissue remodeling during inflammation.

Cebpg encodes CCAAT/enhancer-binding protein gamma, a transcription factor that regulates the expression of genes involved in immune responses and hematopoiesis. Cebpg is known to interact with other C/EBP family members to modulate the transcription of genes required for myeloid differentiation and immune cell function. Its activity is crucial for maintaining the balance between immune activation and suppression. Additionally, Cebpg is implicated in the "Response Of EIF2AK1 (HRI) To Heme Deficiency," which is directly related to heme metabolism and iron homeostasis.

Ido1 encodes indoleamine 2,3-dioxygenase 1, an enzyme that catalyzes the first step in the kynurenine pathway of tryptophan catabolism. IDO1 is a key regulator of immune tolerance, as its activity depletes tryptophan and generates metabolites that suppress T-cell proliferation and promote regulatory T-cell differentiation. This function is particularly important in preventing excessive immune activation and maintaining tissue homeostasis during inflammation. The pathway "Tryptophan catabolism" further highlights its role in immune modulation.

Slc11a2 encodes solute carrier family 11 member 2, also known as DMT1 (divalent metal transporter 1), which is responsible for the uptake of divalent metal ions, including iron, into cells. Iron is essential for numerous cellular processes, but its dysregulation can lead to oxidative stress and pathogen proliferation. SLC11A2 plays a pivotal role in maintaining iron homeostasis, particularly in macrophages, where iron availability influences immune responses and pathogen control.

The interplay between these proteins suggests a coordinated role in modulating immune responses and maintaining iron balance. For example, IDO1-mediated immune suppression may complement the antimicrobial functions of CTSC-activated proteases, while SLC11A2 ensures that iron levels are tightly regulated to prevent pathogen exploitation. CEBPG likely integrates transcriptional regulation of these processes, ensuring appropriate responses to environmental cues.

In summary, this system of interacting proteins performs a dual role in paraferritin complex activity and tryptophan catabolism, with each component contributing to the regulation of inflammation, pathogen defense, and cellular equilibrium.Step 2: Self-verification of the process nameOriginal_claim:The entire gene set is involved in immune modulation and iron homeostasis. Here is the entire gene set used for verification:

##Ctsc,Cebpg,Ido1,Slc11a2##

Verified_claim:The claim that the gene set "Ctsc, Cebpg, Ido1, Slc11a2" is involved in immune modulation and iron homeostasis is supported by the following evidence:

1. **Iron Homeostasis**:

- The enrichment analysis identified the "paraferritin complex" (GO:0070826), which is involved in iron transport and homeostasis. This complex reduces Fe(III) to Fe(II) and is essential for heme synthesis, indicating a role in iron metabolism.

- The pathway analysis revealed the involvement of "Response Of EIF2AK1 (HRI) To Heme Deficiency" (Reactome_2022), which is directly related to iron homeostasis and heme regulation.

2. **Immune Modulation**:

- The pathway analysis identified "Tryptophan catabolism" (BioPlanet_2019 and Reactome_2022), which is mediated by IDO1. Tryptophan catabolism is known to play a role in immune regulation, particularly in modulating immune responses and maintaining immune tolerance.

- The pathway "Interleukin-4 regulation of apoptosis" (BioPlanet_2019) involves genes SLC11A2 and CTSC, which are associated with immune signaling and apoptosis regulation.

Thus, the claim is **strongly supported** by the evidence. The gene set is indeed involved in processes related to immune modulation and iron homeostasis.

Original_claim:The gene set plays a role in regulating immune responses and maintaining iron balance. Here is the entire gene set used for verification:

##Ctsc,Cebpg,Ido1,Slc11a2##

Verified_claim:The claim that the gene set "Ctsc, Cebpg, Ido1, Slc11a2" plays a role in regulating immune responses and maintaining iron balance is supported by multiple pieces of evidence, though the association is partially indirect and requires interpretation.

1. **Iron Balance**:

- Enrichment analysis reveals the involvement of the "paraferritin complex," which is associated with iron transport and reduction of Fe(III) to Fe(II) for heme synthesis. This suggests a role in maintaining iron balance.

- Pathway analysis highlights "Response Of EIF2AK1 (HRI) To Heme Deficiency," which involves CEBPG and is related to heme metabolism, further supporting the connection to iron balance.

2. **Immune Responses**:

- The gene IDO1 is involved in tryptophan catabolism, which plays a role in immunoregulation by modulating T-cell behavior through the catabolism of tryptophan. This is directly linked to immune responses.

- The gene CEBPG is involved in hemopoiesis and natural killer cell-mediated cytotoxicity, which are critical components of immune function.

3. **Gene Summaries**:

- IDO1 is described as playing roles in antimicrobial and antitumor defense, neuropathology, immunoregulation, and antioxidant activity, which aligns with immune response regulation.

- CEBPG is involved in hemopoiesis and regulation of gene expression, including processes relevant to immune function.

4. **Pathways and Enrichment**:

- Pathways such as "Interleukin-4 regulation of apoptosis" and "Tryptophan catabolism" further connect the gene set to immune-related processes.

Decision: The claim is **strongly supported** based on the evidence provided, as the gene set is associated with both immune regulation and iron balance through direct and indirect biological processes.

Original_claim:The entire gene set contributes to processes related to immune system function and iron metabolism. Here is the entire gene set used for verification:

##Ctsc,Cebpg,Ido1,Slc11a2##

Verified_claim:The claim that the gene set "Ctsc, Cebpg, Ido1, Slc11a2" contributes to processes related to immune system function and iron metabolism is **weakly supported**.

Evidence:

1. **Iron Metabolism**:

- The enrichment analysis identified the "paraferritin complex" (GO:0070826), which is involved in iron transport and metabolism, as a significant term. This suggests a connection to iron metabolism.

- The pathway analysis highlighted "Response Of EIF2AK1 (HRI) To Heme Deficiency" (R-HSA-9648895), which involves heme, a component of iron metabolism, and is associated with the gene "CEBPG."

2. **Immune System Function**:

- The pathway analysis identified "Tryptophan catabolism" and "Tryptophan Catabolism R-HSA-71240," which involve the gene "IDO1." Tryptophan metabolism is known to play a role in immune regulation, particularly in immune tolerance and inflammation.

- The pathway "Interleukin-4 regulation of apoptosis" was also identified, which involves immune signaling and apoptosis regulation.

While there is evidence linking the gene set to both iron metabolism and immune system processes, the connections are not comprehensive or strongly established for the entire gene set. Thus, the claim is weakly supported.

Original_claim: Here is the entire gene set used for verification:

##Ctsc,Cebpg,Ido1,Slc11a2##

Verified_claim:The verification process for the gene set Ctsc, Cebpg, Ido1, Slc11a2 did not yield any associated complex protocol IDs, complex names, or related diseases for the individual genes. No evidence was found to support any specific claim based on the provided tools.

Step 1: Initial generation for gene setProcess: Immune Modulation and Iron Homeostasis

The system of interacting proteins encoded by the genes Ctsc, Cebpg, Ido1, and Slc11a2 appears to coordinate processes related to immune regulation and iron metabolism. These processes are critical for maintaining cellular homeostasis and responding to environmental challenges such as infection or inflammation.

Ctsc encodes cathepsin C, a lysosomal cysteine protease involved in the activation of serine proteases in immune cells. This protein plays a key role in regulating the activity of neutrophil elastase, cathepsin G, and other proteases critical for innate immune responses. Its function is essential for pathogen clearance and tissue remodeling during inflammation.

Cebpg encodes CCAAT/enhancer-binding protein gamma, a transcription factor that regulates the expression of genes involved in immune responses and hematopoiesis. Cebpg is known to interact with other C/EBP family members to modulate the transcription of genes required for myeloid differentiation and immune cell function. Its activity is crucial for maintaining the balance between immune activation and suppression.

Ido1 encodes indoleamine 2,3-dioxygenase 1, an enzyme that catalyzes the first step in the kynurenine pathway of tryptophan catabolism. IDO1 is a key regulator of immune tolerance, as its activity depletes tryptophan and generates metabolites that suppress T-cell proliferation and promote regulatory T-cell differentiation. This function is particularly important in preventing excessive immune activation and maintaining tissue homeostasis during inflammation.

Slc11a2 encodes solute carrier family 11 member 2, also known as DMT1 (divalent metal transporter 1), which is responsible for the uptake of divalent metal ions, including iron, into cells. Iron is essential for numerous cellular processes, but its dysregulation can lead to oxidative stress and pathogen proliferation. SLC11A2 plays a pivotal role in maintaining iron homeostasis, particularly in macrophages, where iron availability influences immune responses and pathogen control.

The interplay between these proteins suggests a coordinated role in modulating immune responses and maintaining iron balance. For example, IDO1-mediated immune suppression may complement the antimicrobial functions of CTSC-activated proteases, while SLC11A2 ensures that iron levels are tightly regulated to prevent pathogen exploitation. CEBPG likely integrates transcriptional regulation of these processes, ensuring appropriate responses to environmental cues.

In summary, this system of interacting proteins performs a dual role in immune modulation and iron homeostasis, with each component contributing to the regulation of inflammation, pathogen defense, and cellular equilibrium.

### 4. Per-Notch3 (EC only)

Analysis of gene set: Atp6v0c, Bcl2l15, Tm4sf1, Nxt1, Cldn11, Mboat1, Anapc15, Grb2, Slco1a6, Ptgs1 ...

Step 5: SummarizationProcess: Signal Modulation and Cellular Communication

The system of interacting proteins encoded by the provided gene set plays a central role in signal modulation and cellular communication, with additional contributions to cytoskeletal dynamics, lipid remodeling, and intercellular barrier integrity. Below is a critical analysis of the biological processes performed by this system:

Atp6v0c: Atp6v0c is a subunit of the vacuolar ATPase (V-ATPase) complex, which acidifies intracellular compartments such as endosomes and lysosomes. This acidification is essential for vesicle trafficking, protein sorting, and receptor recycling, contributing to intracellular pH homeostasis and cellular communication.

Grb2: Grb2 acts as an adaptor protein linking receptor tyrosine kinases to downstream signaling pathways. It plays a critical role in receptor-mediated signal transduction by forming complexes with phosphorylated receptors and other signaling proteins, facilitating cellular responses to external stimuli.

Rac1: Rac1 is a small GTPase that regulates actin cytoskeleton dynamics, which are essential for endocytosis, exocytosis, and receptor-mediated signaling. Its role in cytoskeletal reorganization is critical for modulating cellular communication and intracellular trafficking.

Nrgn: Neurogranin binds calmodulin in the absence of calcium, modulating calcium-dependent signaling processes. It plays a significant role in calcium signaling, particularly in neurons, influencing synaptic plasticity and communication.

Ptgs1: Ptgs1 (COX-1) catalyzes the biosynthesis of prostaglandins, which act as signaling molecules in inflammatory and homeostatic processes. This enzymatic activity supports cellular communication in response to physiological and pathological stimuli.

Cldn11: Cldn11 encodes a tight junction protein that regulates paracellular permeability and maintains the integrity of intercellular barriers. This function is essential for cell-cell communication and maintaining tissue homeostasis.

Mboat1: Mboat1 is involved in lipid remodeling, specifically in the acylation of lysophospholipids. This activity supports membrane structure and signaling processes by maintaining the lipid composition required for membrane dynamics.

Anapc15: Anapc15 is a subunit of the anaphase-promoting complex/cyclosome (APC/C), which regulates cell cycle progression through protein polyubiquitination. This process influences the turnover of proteins involved in cellular communication and signaling.

Bcl2l15: Bcl2l15 regulates apoptosis and is involved in the regulation of cellular survival. Its role in apoptosis indirectly supports cellular homeostasis and signaling pathways by maintaining cell viability under stress conditions.

Tm4sf1: Tm4sf1 is a member of the tetraspanin family involved in signal transduction events that regulate cell development, activation, growth, and motility. It contributes to cellular communication by facilitating interactions at the plasma membrane.

Nxt1: Nxt1 is a nuclear export factor involved in mRNA export from the nucleus. By ensuring proper mRNA export, it supports the translation of proteins required for signaling pathways and cellular communication.

Slco1a6: Slco1a6 encodes a solute carrier that facilitates the transport of organic molecules, including bile acids and other anions. This activity may contribute to cellular communication by regulating the availability of signaling molecules.

Farsa: Farsa encodes the alpha subunit of phenylalanyl-tRNA synthetase, which is essential for aminoacyl-tRNA synthesis and protein translation. While its role in signaling is indirect, it ensures the synthesis of proteins required for cellular communication.

In summary, this system of proteins collectively supports signal modulation and cellular communication through roles in receptor-mediated signaling, cytoskeletal dynamics, calcium signaling, and intercellular barrier integrity. Key contributors include Grb2 for signal transduction, Rac1 for cytoskeletal remodeling, and Ptgs1 for prostaglandin-mediated signaling. Additional proteins such as Cldn11 and Mboat1 enhance intercellular communication and membrane structure, while others like Nrgn and Nxt1 modulate calcium signaling and mRNA export.Step 4: Self-verification of the analytical narrativesOriginal_claim:Atp6v0c is involved in maintaining intracellular pH homeostasis through its role in acidifying intracellular compartments, which is critical for vesicle trafficking, protein sorting, and receptor recycling.

Verified_claim:The claim that Atp6v0c is involved in maintaining intracellular pH homeostasis through its role in acidifying intracellular compartments, which is critical for vesicle trafficking, protein sorting, and receptor recycling, is strongly supported.

Evidence:

1. Gene Summary: Atp6v0c is predicted to enable P-type proton-exporting transporter activity and is involved in synaptic vesicle lumen acidification. It is part of the proton-transporting V-type ATPase complex, which is known to play a role in acidifying intracellular compartments. Acidification is essential for processes such as vesicle trafficking, protein sorting, and receptor recycling.

Original_claim:Bcl2l15 regulates apoptosis and may influence cellular survival under stress conditions, indirectly supporting cellular homeostasis and signaling pathways.

Verified_claim:The claim that Bcl2l15 regulates apoptosis and may influence cellular survival under stress conditions, indirectly supporting cellular homeostasis and signaling pathways, is supported by evidence but with some ambiguity.

1. Strong Support:

- The gene summary for Bcl2l15 indicates that it is predicted to be involved in the apoptotic process and regulation of apoptotic processes. This aligns with the claim that Bcl2l15 regulates apoptosis.

2. Weak Support:

- The summary also mentions that Bcl2l15 is active in the cytosol and nucleus, which could be relevant to cellular signaling pathways. However, there is no direct evidence provided about its role in cellular survival under stress conditions or its influence on cellular homeostasis.

3. No Evidence:

- No enrichment functions were retrieved for Bcl2l15, which could have provided additional insights into its involvement in signaling pathways or cellular homeostasis.

Based on the available evidence, the claim is partially verified with strong support for its role in apoptosis regulation but weak support for its influence on cellular survival under stress and indirect support for cellular homeostasis and signaling pathways.

Original_claim:Tm4sf1 facilitates cell adhesion, migration, and vesicle trafficking by interacting with integrins and other membrane proteins, contributing to the organization of signaling complexes at the plasma membrane.

Verified_claim:The claim that "Tm4sf1 facilitates cell adhesion, migration, and vesicle trafficking by interacting with integrins and other membrane proteins, contributing to the organization of signaling complexes at the plasma membrane" is weakly supported based on the following evidence:

1. The gene summary for TM4SF1 indicates that it is a member of the transmembrane 4 superfamily (tetraspanin family) and is involved in signal transduction events that regulate cell development, activation, growth, and motility. This aligns with the claim that TM4SF1 plays a role in cell adhesion and migration. However, the summary does not explicitly mention vesicle trafficking or interactions with integrins and other membrane proteins.

2. No direct interaction data for TM4SF1 with integrins or other membrane proteins was retrieved, leaving the specific mechanism of interaction and its role in organizing signaling complexes at the plasma membrane unverified.

While the general role of TM4SF1 in cell motility and signal transduction is supported, the specific details of its interactions and contributions to vesicle trafficking and signaling complex organization remain ambiguous.

Original_claim:Nxt1 supports cellular communication by ensuring proper mRNA export from the nucleus, which is essential for the translation of proteins involved in signaling pathways.

Verified_claim:The claim that "Nxt1 supports cellular communication by ensuring proper mRNA export from the nucleus, which is essential for the translation of proteins involved in signaling pathways" is **strongly supported**.

Evidence:

1. According to the gene summary for NXT1, the protein encoded by this gene is located in the nuclear envelope and functions as a nuclear export factor. It is involved in both RAN- and CRM1-dependent pathways for exporting U1 snRNA and in CRM1-independent pathways for exporting tRNA and mRNA. This indicates that NXT1 plays a critical role in mRNA export from the nucleus.

2. The summary also mentions that NXT1 heterodimerizes with the Tap protein, potentially regulating the ability of Tap protein to mediate nuclear mRNA export. This further supports its role in ensuring proper mRNA export.

While the claim links mRNA export to signaling pathways, the direct involvement of NXT1 in signaling pathways is not explicitly detailed in the evidence. However, since mRNA export is a prerequisite for protein translation, which includes proteins involved in signaling, the claim is consistent with the known function of NXT1.

Original_claim:Cldn11 regulates paracellular permeability and maintains the integrity of intercellular barriers, influencing cell-cell communication.

Verified_claim:The claim that "Cldn11 regulates paracellular permeability and maintains the integrity of intercellular barriers, influencing cell-cell communication" is strongly supported.

Evidence:

1. According to the gene summary for CLDN11, it encodes a member of the claudin family, which are integral membrane proteins and components of tight junction strands. Tight junction strands serve as physical barriers to prevent solutes and water from passing freely through the paracellular space between epithelial or endothelial cell sheets. This directly supports the role of CLDN11 in regulating paracellular permeability and maintaining intercellular barrier integrity.

2. Tight junctions also play critical roles in maintaining cell polarity and signal transduction, which are essential for cell-cell communication. This aligns with the claim that CLDN11 influences cell-cell communication.

3. The protein encoded by CLDN11 is a major component of central nervous system (CNS) myelin and plays an important role in regulating proliferation and migration of oligodendrocytes, further emphasizing its role in maintaining cellular integrity and communication.

The evidence from the gene summary provides strong support for the claim.

Original_claim:Mboat1 is involved in lipid remodeling, particularly in the acylation of lysophospholipids, which supports membrane structure and signaling processes.

Verified_claim:The claim that Mboat1 is involved in lipid remodeling, particularly in the acylation of lysophospholipids, which supports membrane structure and signaling processes, is strongly supported.

Evidence:

1. Gene Summary: Mboat1 (membrane bound glycerophospholipid O-acyltransferase 1) belongs to the membrane-bound O-acetyltransferase superfamily. It encodes a transmembrane protein that acts as an enzyme transferring organic compounds, preferably from oleoyl-CoA, to hydroxyl groups of protein targets in membranes. This enzymatic activity aligns with lipid remodeling processes, particularly acylation, which is crucial for maintaining membrane structure and signaling functions.

Original_claim:Anapc15, as a subunit of the anaphase-promoting complex/cyclosome (APC/C), regulates cell cycle progression and may influence the turnover of proteins involved in cellular communication through ubiquitination.

Verified_claim:The claim that Anapc15, as a subunit of the anaphase-promoting complex/cyclosome (APC/C), regulates cell cycle progression and may influence the turnover of proteins involved in cellular communication through ubiquitination is **strongly supported**.

Evidences:

1. Gene Summary: Anapc15 is described as being involved in the "anaphase-promoting complex-dependent catabolic process," "protein polyubiquitination," and "regulation of mitotic cell cycle spindle assembly checkpoint." It is explicitly stated to be part of the anaphase-promoting complex (APC/C). This aligns with the claim that Anapc15 regulates cell cycle progression and is involved in ubiquitination processes.

2. No additional complex-specific information was retrieved for Anapc15, but the gene's role in the APC/C and its functions are well-documented in the summary provided.

No contradictory evidence was found.

Original_claim:Grb2 acts as an adaptor protein linking receptor tyrosine kinases to downstream signaling pathways, playing a critical role in receptor-mediated signal transduction.

Verified_claim:The claim that "Grb2 acts as an adaptor protein linking receptor tyrosine kinases to downstream signaling pathways, playing a critical role in receptor-mediated signal transduction" is **strongly supported** by the evidence.

1. Gene Summary Evidence:

- The protein encoded by the GRB2 gene binds the epidermal growth factor receptor (EGFR) and contains one SH2 domain and two SH3 domains. The SH3 domains direct complex formation with proline-rich regions of other proteins, and the SH2 domain binds tyrosine-phosphorylated sequences. This indicates that GRB2 functions as an adaptor protein in signal transduction pathways. Additionally, the gene is similar to the Sem5 gene in C. elegans, which is involved in signal transduction pathways.

2. Enrichment Evidence:

- GRB2 is part of the "Grb2-EGFR complex," which is involved in linking EGFR activation to the p21-Ras pathway. This supports its role in receptor-mediated signal transduction.

- GRB2 is also associated with the "SHC-GRB2 complex," further indicating its role in signaling pathways.

These findings collectively confirm that GRB2 acts as an adaptor protein linking receptor tyrosine kinases to downstream signaling pathways, playing a critical role in receptor-mediated signal transduction.

Original_claim:Slco1a6 facilitates the uptake of organic molecules, potentially transporting substrates that participate in signaling pathways and cellular communication.

Verified_claim:The claim that "Slco1a6 facilitates the uptake of organic molecules, potentially transporting substrates that participate in signaling pathways and cellular communication" is **weakly supported**.

Evidence:

1. The gene summary for Slco1a6 in mice (Mus musculus) indicates that it is predicted to enable bile acid transmembrane transporter activity and sodium-independent organic anion transmembrane transporter activity. It acts upstream of processes such as bile acid metabolic process, cholesterol homeostasis, and triglyceride homeostasis. This suggests that Slco1a6 is involved in the transport of organic molecules, which aligns with the claim.

2. The summary does not explicitly mention the transport of substrates involved in signaling pathways or cellular communication. However, the general role of organic anion transporters in cellular processes could imply a potential connection to signaling and communication, though this is not directly confirmed.

No additional enrichment or pathway data was retrieved to further substantiate the claim.

Original_claim:Ptgs1 (COX-1) is involved in the biosynthesis of prostaglandins, which act as signaling molecules in inflammatory and homeostatic processes.

Verified_claim:The claim that Ptgs1 (COX-1) is involved in the biosynthesis of prostaglandins, which act as signaling molecules in inflammatory and homeostatic processes, is strongly supported.

Evidence:

1. Gene Summary: Ptgs1, also known as COX-1, is described as encoding an enzyme that catalyzes the conversion of arachidonic acid to prostaglandins. Prostaglandins are signaling molecules involved in various biological processes, including inflammation and homeostasis. The encoded protein is also noted for its role in regulating angiogenesis in endothelial cells and its inhibition by nonsteroidal anti-inflammatory drugs such as aspirin. This aligns with the claim regarding its involvement in prostaglandin biosynthesis and signaling in inflammatory and homeostatic processes.

Original_claim:Rac1 regulates actin cytoskeleton dynamics, which are essential for endocytosis, exocytosis, and receptor-mediated signaling, thereby modulating cellular communication.

Verified_claim:The claim that "Rac1 regulates actin cytoskeleton dynamics, which are essential for endocytosis, exocytosis, and receptor-mediated signaling, thereby modulating cellular communication" is strongly supported.

Evidence:

1. Gene Summary: Rac1 is a GTPase belonging to the RAS superfamily of small GTP-binding proteins. Members of this superfamily regulate diverse cellular events, including cytoskeletal reorganization. This aligns with the claim that Rac1 regulates actin cytoskeleton dynamics.

2. Cytoskeletal Reorganization: The gene summary explicitly mentions Rac1's role in cytoskeletal reorganization, which is a fundamental process underlying actin dynamics. Actin cytoskeleton dynamics are critical for cellular processes such as endocytosis, exocytosis, and receptor-mediated signaling.

The evidence directly supports the claim, confirming Rac1's involvement in regulating actin cytoskeleton dynamics and its broader role in cellular communication.

Original_claim:Nrgn modulates calcium-dependent signaling processes by binding to calmodulin, particularly influencing calcium signaling in neurons.

Verified_claim:The claim that Nrgn modulates calcium-dependent signaling processes by binding to calmodulin, particularly influencing calcium signaling in neurons, is strongly supported.

Evidence:

1. According to the gene summary for Nrgn (neurogranin), it is described as a postsynaptic protein kinase substrate that binds calmodulin in the absence of calcium. This indicates its role in calcium signaling processes, particularly in neurons.

2. Neurogranin is identified as a human homolog of the neuron-specific rat RC3/neurogranin gene, further emphasizing its neuronal specificity and involvement in signaling pathways.

No additional biological domains were identified for Nrgn, but the provided summary directly supports the claim.

Original_claim:Farsa ensures the synthesis of proteins required for signaling pathways through its role in aminoacyl-tRNA synthesis, indirectly supporting cellular communication.

Verified_claim:The claim that "Farsa ensures the synthesis of proteins required for signaling pathways through its role in aminoacyl-tRNA synthesis, indirectly supporting cellular communication" is weakly supported.

1. **Gene Function**: FARSA encodes the alpha subunit of phenylalanyl-tRNA synthetase, an enzyme involved in charging tRNAs with their cognate amino acids, specifically phenylalanine. This is a fundamental step in protein synthesis. The gene's product is similar to the catalytic subunit of prokaryotic and yeast phenylalanyl-tRNA synthetases (PheRS) and is expressed in a regulated manner (source: gene summary).

2. **Enrichment Analysis**: FARSA is associated with the "phenylalanine-tRNA ligase complex," which catalyzes the ligation of phenylalanine to tRNA(Phe), forming L-phenylalanyl-tRNA(Phe). This activity is essential for protein synthesis but does not directly indicate a specific role in signaling pathways (source: enrichment analysis).

3. **Pathway Analysis**: FARSA is involved in pathways such as "Cytosolic tRNA Aminoacylation" and "tRNA Aminoacylation," which are critical for the translation process. While these pathways are fundamental to protein synthesis, their direct link to signaling pathways or cellular communication is not explicitly established (source: pathway analysis).

The claim connects FARSA's role in aminoacyl-tRNA synthesis to signaling pathways and cellular communication. While protein synthesis is essential for all cellular functions, including signaling, the evidence does not directly confirm FARSA's specific role in supporting signaling pathways or cellular communication. Thus, the support for the claim is weak.

Original_claim:

Verified_claim:Failed.

Step 3: ModificationProcess: Signal Modulation and Cellular Communication

The system of interacting proteins encoded by the provided gene set appears to play a central role in signal modulation and cellular communication, with additional contributions to lipid metabolism, cytoskeletal dynamics, and intercellular communication. Below is a critical analysis of the biological processes performed by this system:

Atp6v0c: This gene encodes a subunit of the vacuolar ATPase (V-ATPase) complex, which is essential for acidifying intracellular compartments such as endosomes, lysosomes, and secretory vesicles. Acidification is critical for vesicle trafficking, protein sorting, and receptor recycling. The involvement of Atp6v0c suggests that the system is integral to maintaining proper intracellular pH homeostasis.

Bcl2l15: This gene encodes a member of the Bcl-2 family, which is involved in regulating apoptosis. While its role in vesicular transport is less direct, Bcl2l15 may influence the survival of cells under stress conditions, ensuring the persistence of cellular processes. Its function may also intersect with signaling pathways that regulate cellular homeostasis.

Tm4sf1: Tetraspanin-4 superfamily member 1 is implicated in cell adhesion, migration, and vesicle trafficking. It is known to interact with integrins and other membrane proteins, facilitating the organization of signaling complexes. Tm4sf1 likely contributes to the modulation of membrane interactions and signaling events at the plasma membrane.

Nxt1: Nuclear export transporter 1 is essential for mRNA export from the nucleus. While its primary role is in nucleocytoplasmic transport, Nxt1 indirectly supports cellular communication by ensuring the proper translation of proteins required for signaling pathways.

Cldn11: Claudin-11 is a tight junction protein that regulates paracellular permeability. Its presence in this system suggests a role in maintaining the integrity of intercellular barriers, which may influence cell-cell communication.

Mboat1: Membrane-bound O-acyltransferase 1 is involved in lipid remodeling, particularly in the acylation of lysophospholipids. Lipid composition is critical for membrane structure and signaling. Mboat1 likely supports the structural integrity of membranes involved in signaling processes.

Anapc15: This gene encodes a subunit of the anaphase-promoting complex/cyclosome (APC/C), a ubiquitin ligase that regulates cell cycle progression. While its direct role in signaling is unclear, Anapc15 may influence the turnover of proteins involved in cellular communication through ubiquitination.

Grb2: Growth factor receptor-bound protein 2 is an adaptor protein that links receptor tyrosine kinases to downstream signaling pathways. Grb2 is critical for signal transduction and may regulate receptor-mediated signaling events, thereby modulating cellular responses to external stimuli.

Slco1a6: This gene encodes a solute carrier organic anion transporter, which facilitates the uptake of organic molecules. Its role in cellular communication may involve the transport of specific substrates that participate in signaling pathways.

Ptgs1: Prostaglandin-endoperoxide synthase 1 (COX-1) is involved in the biosynthesis of prostaglandins. Prostaglandins can act as signaling molecules that influence cellular communication, particularly in inflammatory and homeostatic processes.

Rac1: Rac1 is a small GTPase that regulates actin cytoskeleton dynamics. It is essential for processes such as endocytosis and exocytosis, which are critical for receptor-mediated signaling and cellular communication. Rac1's role in cytoskeletal remodeling is critical for the spatial and temporal regulation of these processes.

Nrgn: Neurogranin is a calmodulin-binding protein involved in calcium signaling. Calcium is a key regulator of many signaling pathways. Nrgn may modulate calcium-dependent signaling processes, particularly in neurons.

Farsa: Phenylalanyl-tRNA synthetase alpha subunit is involved in aminoacyl-tRNA synthesis, which is essential for protein translation. While its role in signaling is indirect, Farsa ensures the synthesis of proteins required for signaling pathways.

In summary, this system of interacting proteins collectively supports signal modulation and cellular communication through roles in signaling pathways, membrane interactions, and cytoskeletal dynamics. Key contributors include Grb2 for signal transduction, Rac1 for cytoskeletal dynamics, and Ptgs1 for prostaglandin-mediated signaling. Additional proteins such as Mboat1 and Tm4sf1 enhance membrane structure and interactions, while others like Nrgn and Cldn11 modulate calcium signaling and intercellular communication.Step 2: Self-verification of the process nameOriginal_claim:The entire gene set is involved in Vesicular Transport and Signal Modulation. Here is the entire gene set used for verification:

##Atp6v0c,Bcl2l15,Tm4sf1,Nxt1,Cldn11,Mboat1,Anapc15,Grb2,Slco1a6,Ptgs1,Rac1,Nrgn,Farsa##

Verified_claim:The claim that the entire gene set is involved in Vesicular Transport and Signal Modulation is weakly supported.

Evidence:

1. Pathway Analysis:

- The gene set overlaps with pathways such as "CD28-dependent Vav1 pathway," "Cell-cell communication," "CREB transcription factor and its extracellular signals," and "Fc epsilon RI signaling pathway." These pathways are related to signaling and cellular communication, which could be loosely associated with signal modulation.

- No direct evidence of involvement in vesicular transport was found in the pathway analysis.

2. Enrichment Analysis:

- The enrichment results include terms like "BCR-ABL (p210 fusion protein)-GRB2 complex" and "BCR-ABL (p185 fusion protein)-GRB2 complex," which involve GRB2, a gene known for its role in signal transduction. This provides some support for signal modulation.

- No enrichment terms directly related to vesicular transport were identified.

Conclusion:

While there is evidence of involvement in signal modulation through pathways and enrichment terms, the claim of involvement in vesicular transport is not directly supported by the provided data. Therefore, the claim is weakly supported.

Original_claim:The gene set plays a role in cellular processes related to vesicle-mediated transport and signaling pathways. Here is the entire gene set used for verification:

##Atp6v0c,Bcl2l15,Tm4sf1,Nxt1,Cldn11,Mboat1,Anapc15,Grb2,Slco1a6,Ptgs1,Rac1,Nrgn,Farsa##

Verified_claim:The claim that the gene set plays a role in cellular processes related to vesicle-mediated transport and signaling pathways is supported by evidence, though the association is not entirely direct and could be considered as weak support.

1. Enrichment Analysis:

- The gene set is associated with signaling pathways, as evidenced by the involvement of GRB2 in complexes such as "BCR-ABL (p210 fusion protein)-GRB2 complex" and "BCR-ABL (p185 fusion protein)-GRB2 complex." GRB2 is a known adaptor protein involved in signal transduction.

2. Pathway Analysis:

- The gene set overlaps with pathways such as "CD28-dependent Vav1 pathway," "Cell-cell communication," "CREB transcription factor and its extracellular signals," and "Fc epsilon RI signaling pathway." These pathways are related to signaling processes, with GRB2 and RAC1 being key contributors.

While the evidence supports the involvement of the gene set in signaling pathways, there is no direct evidence linking the gene set to vesicle-mediated transport. Therefore, the claim is partially supported, with strong evidence for signaling pathways but weak or no evidence for vesicle-mediated transport.

Original_claim:The entire gene set contributes to the regulation of vesicular trafficking and signal transduction mechanisms. Here is the entire gene set used for verification:

##Atp6v0c,Bcl2l15,Tm4sf1,Nxt1,Cldn11,Mboat1,Anapc15,Grb2,Slco1a6,Ptgs1,Rac1,Nrgn,Farsa##

Verified_claim:The claim that the gene set contributes to the regulation of vesicular trafficking and signal transduction mechanisms is weakly supported based on the evidence.

1. Enrichment Analysis:

- The enrichment analysis reveals associations with signaling-related complexes such as "BCR-ABL (p210 fusion protein)-GRB2 complex" and "BCR-ABL (p185 fusion protein)-GRB2 complex." GRB2 is a known adaptor protein involved in signal transduction pathways. However, there is no direct evidence linking the entire gene set to vesicular trafficking.

2. Pathway Analysis:

- The pathway analysis identifies several signal transduction-related pathways, including "CD28-dependent Vav1 pathway," "CREB transcription factor and its extracellular signals," and "Fc epsilon RI signaling pathway," with overlapping genes GRB2 and RAC1. These pathways are involved in cellular signaling mechanisms. Additionally, "Cell-cell communication" is identified, which may indirectly relate to vesicular trafficking.

While some genes in the set (e.g., GRB2 and RAC1) are involved in signal transduction, the evidence does not conclusively demonstrate that the entire gene set contributes to vesicular trafficking. Therefore, the claim is weakly supported.

Original_claim: Here is the entire gene set used for verification:

##Atp6v0c,Bcl2l15,Tm4sf1,Nxt1,Cldn11,Mboat1,Anapc15,Grb2,Slco1a6,Ptgs1,Rac1,Nrgn,Farsa##

Verified_claim:

The gene set "Atp6v0c,Bcl2l15,Tm4sf1,Nxt1,Cldn11,Mboat1,Anapc15,Grb2,Slco1a6,Ptgs1,Rac1,Nrgn,Farsa" was analyzed for its enrichment functions and complex associations.

1. **Complex Associations**:

- No direct complex protocol IDs or complex names were found for the entire gene set.

2. **Enrichment Functions**:

- The gene set is significantly associated with the following:

- **hsa-mir-663**: A microRNA (MIRNA) with a p-value of 0.0243.

- **hsa-mir-1908**: Another microRNA (MIRNA) with a p-value of 0.0243.

- **Factor ZNF718; motif NGGGRATWGCGM**: A transcription factor (TF) with a p-value of 0.0493.

- **BCR-ABL (p210 fusion protein)-GRB2 complex**: A protein complex involving GRB2 with a p-value of 0.0499.

- **BCR-ABL (p185 fusion protein)-GRB2 complex**: Another protein complex involving GRB2 with a p-value of 0.0499.

These findings suggest that the gene set has significant associations with specific microRNAs, transcription factors, and protein complexes, particularly involving GRB2. However, the evidence does not provide a direct or comprehensive link to broader biological functions or diseases without further context.

Step 1: Initial generation for gene setProcess: Vesicular Transport and Signal Modulation

The system of interacting proteins encoded by the provided gene set appears to play a central role in vesicular transport and signal modulation, with additional contributions to lipid metabolism, cytoskeletal dynamics, and intercellular communication. Below is a critical analysis of the biological processes performed by this system:

Atp6v0c: This gene encodes a subunit of the vacuolar ATPase (V-ATPase) complex, which is essential for acidifying intracellular compartments such as endosomes, lysosomes, and secretory vesicles. Acidification is critical for vesicle trafficking, protein sorting, and receptor recycling. The involvement of Atp6v0c suggests that the system is integral to maintaining proper vesicular transport and intracellular pH homeostasis.

Bcl2l15: This gene encodes a member of the Bcl-2 family, which is involved in regulating apoptosis. While its role in vesicular transport is less direct, Bcl2l15 may influence the survival of cells under stress conditions, ensuring the persistence of vesicle-mediated processes. Its function may also intersect with signaling pathways that regulate cellular homeostasis.

Tm4sf1: Tetraspanin-4 superfamily member 1 is implicated in cell adhesion, migration, and vesicle trafficking. It is known to interact with integrins and other membrane proteins, facilitating the organization of signaling complexes. Tm4sf1 likely contributes to the modulation of vesicle docking and fusion events at the plasma membrane.

Nxt1: Nuclear export transporter 1 is essential for mRNA export from the nucleus. While its primary role is in nucleocytoplasmic transport, Nxt1 indirectly supports vesicular transport by ensuring the proper translation of proteins required for vesicle formation and trafficking.

Cldn11: Claudin-11 is a tight junction protein that regulates paracellular permeability. Its presence in this system suggests a role in maintaining the integrity of intercellular barriers, which may influence vesicle-mediated communication between cells.

Mboat1: Membrane-bound O-acyltransferase 1 is involved in lipid remodeling, particularly in the acylation of lysophospholipids. Lipid composition is critical for vesicle formation, curvature, and fusion. Mboat1 likely supports the structural integrity of vesicles and their membranes.

Anapc15: This gene encodes a subunit of the anaphase-promoting complex/cyclosome (APC/C), a ubiquitin ligase that regulates cell cycle progression. While its direct role in vesicular transport is unclear, Anapc15 may influence the turnover of proteins involved in vesicle trafficking through ubiquitination.

Grb2: Growth factor receptor-bound protein 2 is an adaptor protein that links receptor tyrosine kinases to downstream signaling pathways. Grb2 is critical for signal transduction and may regulate vesicle-mediated endocytosis of activated receptors, thereby modulating cellular responses to external stimuli.

Slco1a6: This gene encodes a solute carrier organic anion transporter, which facilitates the uptake of organic molecules. Its role in vesicular transport may involve the loading of specific substrates into vesicles for secretion or intracellular trafficking.

Ptgs1: Prostaglandin-endoperoxide synthase 1 (COX-1) is involved in the biosynthesis of prostaglandins. Prostaglandins can act as signaling molecules that influence vesicle trafficking and secretion, particularly in inflammatory and homeostatic processes.

Rac1: Rac1 is a small GTPase that regulates actin cytoskeleton dynamics. It is essential for vesicle movement along cytoskeletal tracks and for processes such as endocytosis and exocytosis. Rac1's role in cytoskeletal remodeling is critical for the spatial and temporal regulation of vesicular transport.

Nrgn: Neurogranin is a calmodulin-binding protein involved in calcium signaling. Calcium is a key regulator of vesicle fusion and neurotransmitter release. Nrgn may modulate calcium-dependent vesicular processes, particularly in neurons.

Farsa: Phenylalanyl-tRNA synthetase alpha subunit is involved in aminoacyl-tRNA synthesis, which is essential for protein translation. While its role in vesicular transport is indirect, Farsa ensures the synthesis of proteins required for vesicle formation and trafficking.

In summary, this system of interacting proteins collectively supports vesicular transport and signal modulation through roles in vesicle formation, trafficking, fusion, and signaling. Key contributors include Atp6v0c for vesicle acidification, Rac1 for cytoskeletal dynamics, and Grb2 for signal transduction. Additional proteins such as Mboat1 and Tm4sf1 enhance vesicle structure and membrane interactions, while others like Nrgn and Ptgs1 modulate signaling pathways that influence vesicular processes.
